# Programming Probiotics: Diet-responsive gene expression and colonization control in engineered *S. boulardii*

**DOI:** 10.1101/2023.11.17.567539

**Authors:** Deniz Durmusoglu, Daniel J. Haller, Ibrahim S. Al’Abri, Katie Day, Carmen Sands, Andrew Clark, Adriana San-Miguel, Ruben Vazquez-Uribe, Morten O. A. Sommer, Nathan C. Crook

**Affiliations:** North Carolina State University; Technical University of Denmark

## Abstract

*Saccharomyces boulardii* (*Sb*) is an emerging probiotic chassis for delivering biomolecules to the mammalian gut, offering unique advantages as the only eukaryotic probiotic. However, precise control over gene expression and gut residence time in *Sb* have remained challenging. To address this, we developed five ligand-responsive gene expression systems and repaired galactose metabolism in *Sb*, enabling inducible gene expression in this strain. Engineering these systems allowed us to construct AND logic gates, control the surface display of proteins, and turn on protein production in the mouse gut in response to a dietary sugar. Additionally, repairing galactose metabolism expanded *Sb*’s habitat within the intestines and resulted in galactose-responsive control over gut residence time. This work opens new avenues for precise dosing of therapeutics by *Sb* via control over its *in vivo* gene expression levels and localization within the gastrointestinal tract.

## Introduction

Engineered probiotics are microorganisms that are genetically engineered to deliver biotherapeutics, detect disease biomarkers, or perform other beneficial functions in the gut environment, therefore providing unique advantages for disease prevention and treatment (1). Specifically, engineered probiotics can convert nutrients available in the gut to molecules that are not, or cannot be readily biosynthesized by the host (1). Additionally, probiotics can be engineered to eliminate undesirable molecules, such as antinutrients or toxins. Recently, probiotics have been engineered to target pathogens such as *Clostridioides difficile*, treat metabolic diseases such as phenylketonuria, and deliver biotherapeutics such as interleukin-10 to treat inflammatory bowel disease (2, 3). Furthermore, engineered probiotics can sense and respond to their environment, improving targeting and dosage of therapeutics (4). Engineered probiotics can therefore potentially be used as a drug delivery platform for current therapies, or enable new treatments for unmet needs in the gut environment. While all engineered probiotics that are currently in clinical trials are bacterial probiotics, engineered yeast probiotics have seen growing preclinical interest due to their high rates of protein secretion, stability under lyophilization, and ability to function during antibiotic treatment (5–10).

*Saccharomyces boulardii* (*S. boulardii* or *Sb*) is a probiotic yeast that was first isolated from lychee and mangosteen by Henri Boulard in 1923 (11, 12). Compared to *Saccharomyces cerevisiae* (with which it shares 99% genomic relatedness) (13), *Sb* can better tolerate low pH and human body temperature, enabling improved survival in the human gut (14–16). *Sb* is currently used to treat ulcerative colitis, diarrhea, and recurrent *Clostridium difficile* infection (17–21). Wild-type *Sb* is generally recognized as safe (GRAS) and persists in conventionally-raised mice up to 24 hours, antibiotic-treated mice for up to 10 days, and in humans for approximately one week (22–24). Recently, a suite of genetic engineering tools for *Sb* have been developed, including transformation methods, constitutive promoters, genome editing protocols, genomic integration sites, G protein-coupled receptors, and secretion-enhancing gene knockouts (23, 25–27).

The overall activity of a probiotic-delivered function is the product of per-cell gene expression level and the abundance of the probiotic at the target site. A major challenge in the development of engineered probiotics is control over these two properties. If the expression of a therapeutic is constitutive, i.e. “always on”, then cessation of treatment becomes dependent on probiotic washout, which can be variable between individuals (4). Additionally, biosynthesis of a therapeutic is often energy- and nutrient-intensive, decreasing the fitness of the engineered probiotic and reducing its abundance in the gut (28). To date, all *in vivo* applications of *Sb* have used constitutive expression systems. Therefore, it would be advantageous to control the timing and dosage of the delivered therapeutic in *Sb.* To enable such control in other organisms, ligand-responsive gene expression systems are often employed (29, 30). Ligand-responsive gene expression systems can activate or deactivate transcription of a specific gene in the presence of a particular inducer molecule (31). For *Sb*, this molecule could be provided through the patient’s diet/medication (if *in vivo* expression is desired) or directly to the culture medium (if expression during production is desired). The level of gene expression is thus regulated by the concentration of inducer molecule provided, enabling custom dosage of a therapeutic. Ligand-responsive expression systems also enable construction of more complex circuits, such as logical AND and OR gates, enabling more control over the behavior of engineered probiotics (32, 33). Additionally, ligand-responsive expression systems can improve the biosafety of the therapeutic, as human-to-human spread of the engineered probiotic does not result in drug delivery of the therapeutic to unintended individuals (34, 35).

In this work, we sought to investigate the performance in *Sb* of ligand-responsive gene expression systems that were previously investigated in *S. cerevisiae*, and demonstrate their utility for several *in vivo* applications. We first constructed *SbGal⁺*, a galactose-competent strain of *Sb*, and demonstrated its improved growth relative to wild-type on galactose and other sugars. We next investigated the dose-response behavior in *SbGal⁺* of five inducible promoters that respond to galactose, xylose, lactose or IPTG (Isopropyl ß-D-1-thiogalactopyranoside), copper, and anhydrotetracycline, respectively, and span a variety of gene activation ranges under both aerobic and anaerobic (i.e. gut-like) conditions. Several of these inducible systems are regulated by non-toxic (e.g. galactose, IPTG, xylose) or minimally toxic (e.g. aTc at the tested concentrations) small molecules. We then demonstrated the applicability of this inducible promoter set for applications such as yeast surface display and logic gate construction. When delivered to the mouse gut, we showed that the *SbGal⁺* strain exhibits improved colonization when galactose is added, likely due to galactose’s dual role as a carbon source. Finally, we constructed a logical AND gate using these inducible promoters, and demonstrated programmable control of gene expression in probiotic yeast during passage through the mouse gut. This research therefore unveils enhanced, precision-controlled *Sb* expression systems, which will find utility in advancing personalized disease intervention strategies.

## Results and Discussion

### *Repair of S. boulardii PGM2* gene enables growth on galactose and raffinose

We first chose to investigate the galactose-inducible promoter *pGAL1* for our toolkit of inducible systems in *Sb*. *pGAL1* is one of the most commonly used promoters for inducible gene expression in *S. cerevisiae* due its extensive characterization and high dynamic range (36). Also, as one of the monosaccharides comprising the lactose disaccharide, it is generally non-toxic to humans, except in the case of galactosemia, a rare genetic disorder that prevents galactose metabolism (37). Liu et al demonstrated that the *PGM2* gene in *Sb* MYA-796 harbors a point mutation that introduces a premature stop codon (**Fig. 1A**), leading to a truncated phosphoglucomutase enzyme and a very slow growth rate of *Sb* on galactose (38). Liu et al also demonstrated that when *Sb*’s *PGM2* gene is reverted to the sequence found in *S. cerevisiae*, growth on galactose is restored (**Fig. 1B**). However, this study did not investigate the functionality of the *pGAL1* promoter in either wildtype *Sb* or in *Sb* with the *PGM2* gene repaired (henceforth *SbGal⁺*) (38).

**Figure 1.**
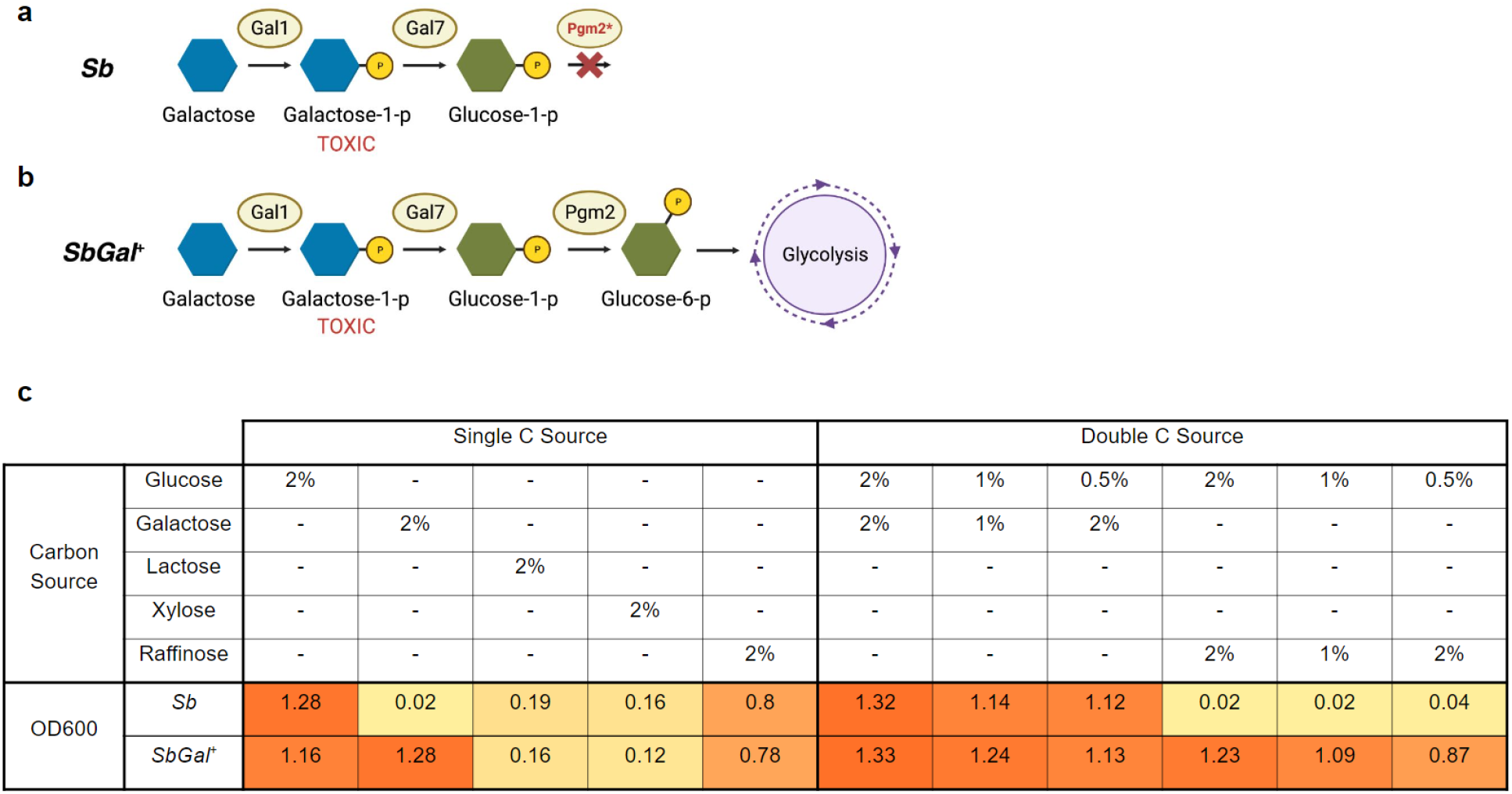
Repair of *PGM2* enables metabolism of galactose by *S. boulardii*. (**a**) Diagram illustrating the galactose utilization pathway in *Sb*, with an inactive *PGM2* enzyme leading to toxic intermediate accumulation. (**b**) The engineered *SbGal⁺* pathway, showing the restoration of *PGM2* activity, allowing for efficient galactose metabolism. (**c**) Growth comparison in complete synthetic media (CSM) with various carbon sources for wild-type *Sb* MYA-796 and the genetically repaired *Sb* MYA-796 ( *SbGal⁺*). The data illustrate the improved growth of *SbGal⁺* on 2% galactose, demonstrating the benefits of *PGM2* repair (orange highlighting). Minimal to no growth difference between *Sb* and *SbGal⁺* was observed on alternative sugars like xylose and lactose, which do not utilize the galactose metabolic pathway. *SbGal⁺* enhanced growth when raffinose is present with glucose, suggesting the strain’s potential for improved performance in complex sugar environments, such as the gut. Values represent the averages of endpoint optical densities of three biological replicates grown in the indicated media for 36h.

In order to enable cells to utilize galactose as both a nutrient source and as an inducer for gene expression, we first measured the growth of *Sb* and *SbGal⁺* on media containing a variety of carbon sources. As shown in **Fig. 1C**, wildtype *Sb* grows very slowly at 37°C on media with 2% galactose as the sole carbon source, while *SbGal⁺* grows at a much higher rate (**Fig. 1C**, **S1B**, and **S1M**), in agreement with Liu et al. Additionally, *SbGal⁺* appears to grow to a slightly lower optical density than wild-type *Sb* when grown on 2% glucose at 37°C (**Fig. 1C** and **S1A**). This is consistent with the hypothesis that the point mutation in *PGM2* enables better glucose utilization at 37°C in *Sb* (38). Notably, the growth advantage of *Sb* on glucose is eliminated at 30°C (**Fig. S1L**). Repair of the *PGM2* gene had minimal to no effect on the growth of *Sb* in media with xylose or lactose as the sole carbon source at 37°C or 30°C (**Fig. 1C**, **S1C-D**, and **S1N-O**). We also investigated the growth of *SbGal⁺* on raffinose, a trisaccharide containing units of galactose, glucose, and fructose. We found that *SbGal⁺* exhibits slightly improved growth relative to *Sb* at 37°C with raffinose as the sole carbon source (**Fig. 1C** and **S1E**) but similar growth as *Sb* at 30°C (**Fig. S1P**), possibly due to its improved ability to utilize the galactose monomer that may result from raffinose metabolism.

**Figure S1.**
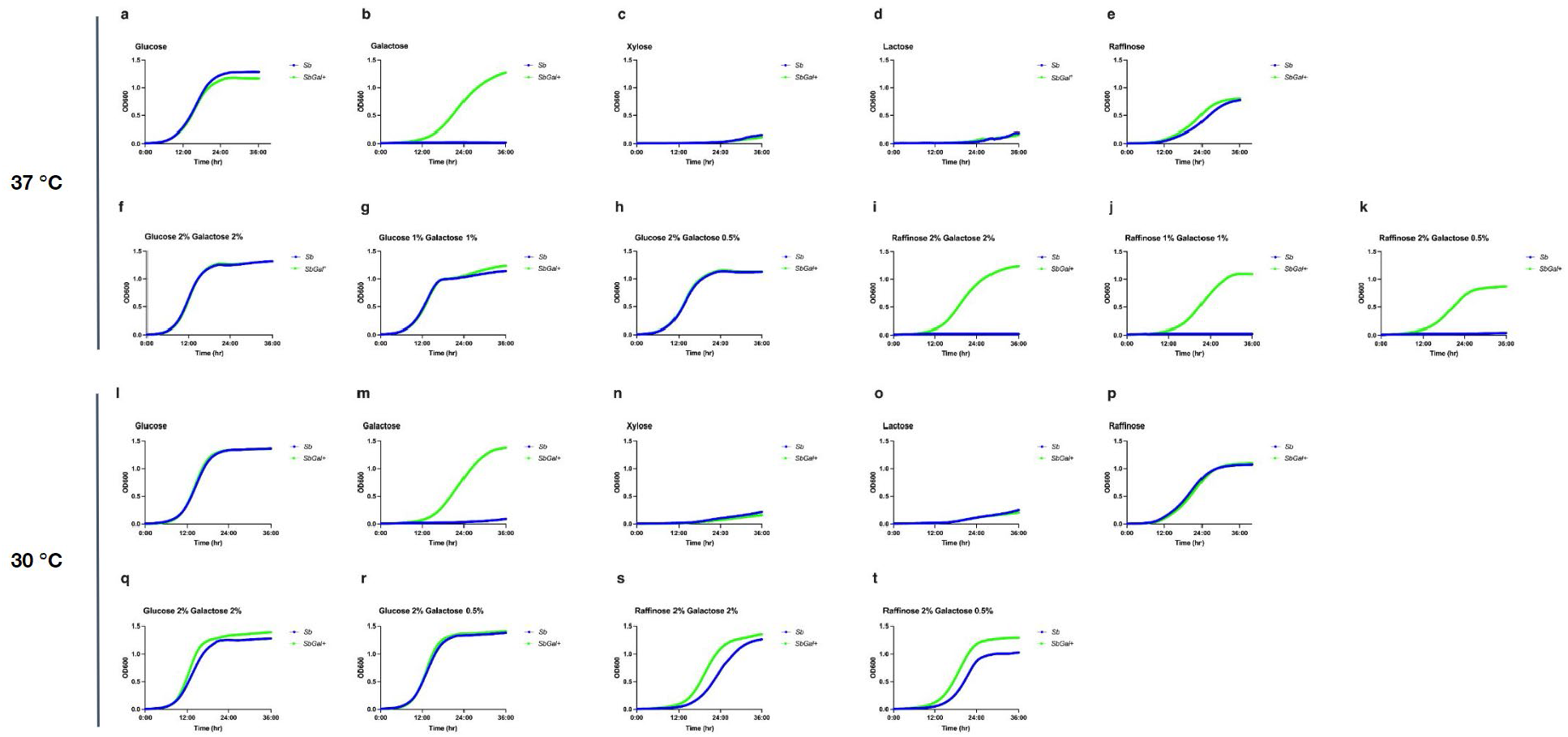
Repaired galactose metabolism enables growth in inducing sugars at 37 °C and 30 °C. (**a-k**) Growth of *Sb* and *SbGal⁺* in complete synthetic media (CSM) with various carbon sources at 37 °C. (**a**) *Sb* and *SbGal⁺* grow equivalently in glucose-containing media. (**b**) *SbGal⁺* exhibits a marked proliferation advantage in galactose-containing media. (**c-d**) Both strains show minimal growth on xylose or lactose, respectively. (**e**) *SbGal⁺* reveals a slightly enhanced ability to utilize raffinose at 37 °C. (**f-g**) Examination of growth in mixed glucose and galactose, underscoring the diauxic shift behavior of *SbGal⁺* when presented with limited glucose and abundant galactose. (**h**) Both strains grow equally well when glucose is in excess. (**i-k**) A stark contrast in growth is observed at 37°C with only raffinose and galactose available, highlighting the superior growth of *SbGal⁺*. (**l-t**) Comparative growth analyses of *Sb* and *SbGal⁺* in complete synthetic media (CSM) with various carbon sources at 30°C. (**l-p**) Growth *Sb* and *SbGal⁺* in complete synthetic media (CSM) with various single carbon sources. (**q-t**) Examination of growth in mixed carbon sources at 30°C, again showing underscoring the diauxic shift behavior of *SbGal⁺* when presented with limited glucose or raffinose and abundant galactose. Three biological replicates were used for all growth curve measurements.

To further characterize the carbon metabolism of *SbGal⁺*, we compared its growth in mixed carbon sources to that of *Sb.* We found no apparent differences in growth between the two strains when cultivated at 37°C under media containing 2% glucose and 2% galactose (**Fig. 1C** and **S1F**). When the concentration of glucose was more limited (1% glucose, 1% galactose), *Sb* and *SbGal⁺* exhibited similar growth for the first ∼18 hours (**Fig. S1G**). After this point, however, the growth rate of *Sb* slowed, likely due to the depletion of glucose in the media, while the growth rate of *SbGal⁺* remained high as it was able to switch to galactose metabolism after glucose depletion. This diauxic shift behavior indicates that glucose is the preferred carbon source for *SbGal⁺* (much like *S. cerevisiae*), but that *SbGal⁺* is capable of efficiently metabolizing galactose when glucose is scarce (39). No difference between the two strains was observed at 37°C when glucose was in excess of galactose (i.e. 2% glucose, 0.5% galactose, **Fig. 1C**, **S1H**). When only raffinose and galactose are available as carbon sources, *Sb* grows extremely slowly at 37°C, while *SbGal⁺* grows well (**Fig. S1I-K**). The growth of *Sb* on mixed raffinose/galactose media improved at 30°C, but the growth of *SbGal⁺* was still superior (**Fig. S1S-T**). These results demonstrate that *SbGal⁺* is able to efficiently utilize galactose and raffinose as carbon sources, particularly at human body temperature.

### Inducible systems enable tunable gene expression in *S. boulardii*

Having confirmed expanded carbohydrate metabolism in *SbGal⁺*, we next asked whether and to what extent gene expression could be induced by common small molecule inducers. For maximum compatibility with existing yeast toolkits, we used the galactose-inducible pG*AL1* promoter and the copper-inducible *pCUP1* promoter from the MoClo Yeast Toolkit (YTK) (40), as well as three engineered minimal promoters previously described: *pTET* (inducible by anhydrotetracycline (aTc)), *pLAC* (inducible by IPTG), and *pXYL* (inducible by xylose) (41). All three minimal promoters consist of two repressor-binding operator sequences, separated by spacers, placed upstream of a minimal *ADH2* transcriptional start site. To investigate the dose-response characteristics of these promoters in *SbGal⁺*, we constructed plasmids encoding yeast-enhanced green fluorescent protein (yeGFP) under the control of each of the inducible promoters and transformed these plasmids to *SbGal⁺*. For those strains harboring the *pTET*, *pLAC*, and *pXYL* promoters, we also introduced plasmids encoding the corresponding repressor proteins (*tetR*, *lacI*, and *xylR*, respectively) under the control of constitutive promoters. Glucose was used as the carbon source for all strains except the *pGAL1-yeGFP* strain, for which raffinose was used as a carbon source to avoid repression of *pGAL1* by glucose.

All five systems showed increasing fluorescence with increasing inducer concentration, demonstrating ligand-responsive gene expression in *Sb* (**Fig. 2A-E**). The *pCUP1* promoter exhibited the highest fluorescence level at its minimum inducer concentration (0.05 uM), matching previous reports that it is a “leaky” promoter (40). *pCUP1* also exhibited the highest maximal fluorescence, followed by *pGAL1*, *pLAC*, *pXYL*, and *pTET*. *pGAL1* and *pLAC* had tighter off-states, with negligible (no fluorescence above background) fluorescence detected for the lowest inducer concentrations tested (0.002% galactose and 0.5 mM IPTG). *pGAL1* had the highest fold change (226.9), followed by *pLAC* (17.9), *pTET (*16.3), *pXYL* (10.3) and *pCUP1* (3.3). Notably, *pLAC* required a high concentration of IPTG to reach high expression levels and did not reach saturation even at 500 mM IPTG. In contrast, the *pGAL1* promoter was sensitive to low concentrations of galactose, exhibiting an increase in fluorescence at less than 0.01% (w/v) galactose. While *pGAL1* does respond to galactose in *Sb*, *SbGal⁺* enables simultaneous growth and induction in galactose alone (**Fig. S2A-B**) or in the presence of both galactose and raffinose (**Fig. S2C**). Although raffinose does contain a galactose monosaccharide, raffinose alone does not significantly activate transcription from the *pGAL1* promoter in *Sb* or *SbGal⁺* (**Fig. S2C**). The fold induction, sensitivity, and basal/peak expression levels we observed for these inducible systems, as well as inducer characteristics such as cost, toxicity, and stability, indicates that these inducible systems could be applicable across a wide range of application areas.

**Figure 2.**
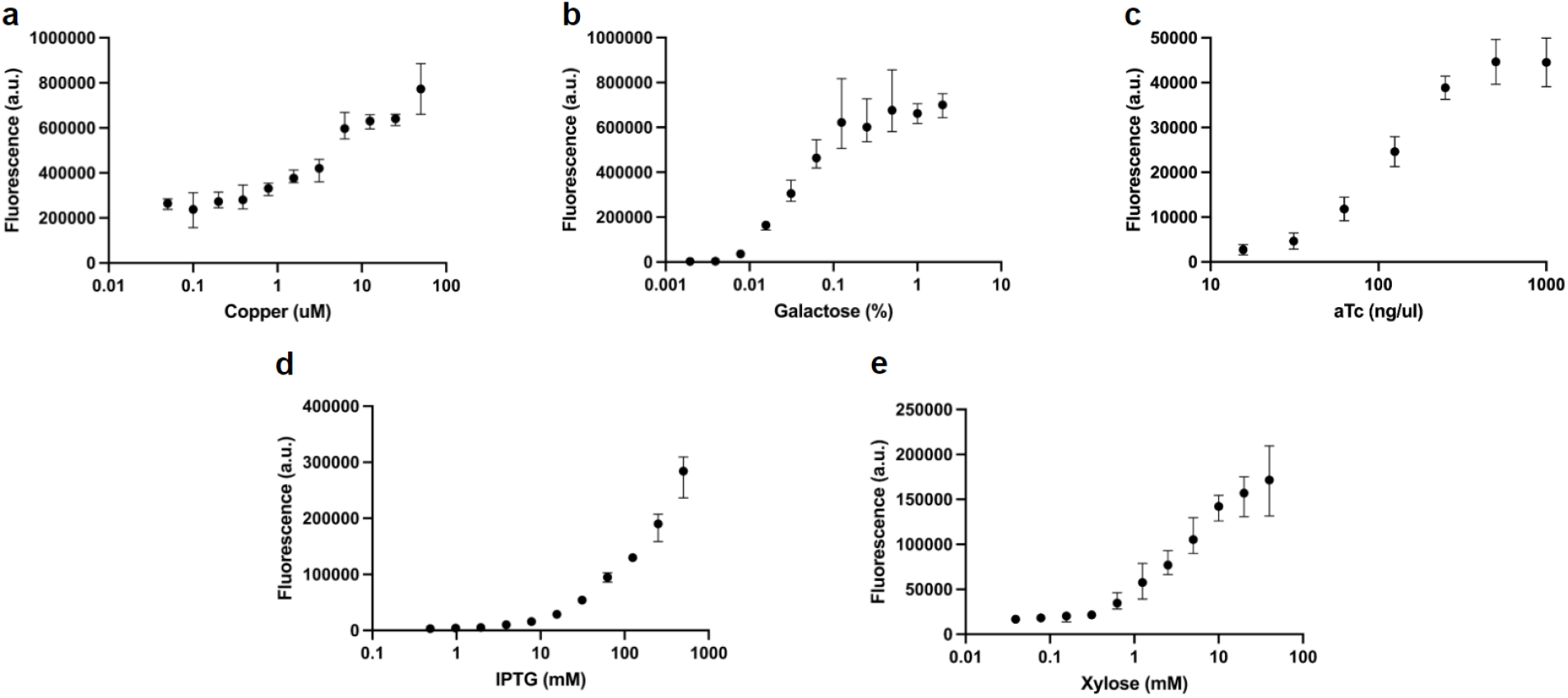
Inducible systems enable tunable gene expression in probiotic yeast. yeGFP was used to measure the activity of each promoter: *pCUP1* (**a**), *pGAL1* (**b**), *pTET* (**c**), *pLAC* (**d**) and *pXYL* (**e**) in response to different concentrations of galactose (**a**), copper (**b**), aTc (**c**), IPTG (**d**) and xylose (**e**). Inducible promoter-yeGFP constructs were placed on a high-copy (2µ) plasmid with a *URA3* selective marker. For systems using heterologous repressors (tetR, lacI, and xylR), these repressors were expressed from a low-copy (CEN) plasmid with a *HIS3* marker. Strains containing the *pGAL1* construct were cultured at 37 °C in CSM media lacking uracil and supplemented with raffinose (2%) as a carbon source and exposed to various concentrations of galactose as an inducer. Strains containing the *pCUP1*, *pTET*, *pLAC*, and *pXYL* constructs were cultured at 37 °C in CSM media lacking either uracil or uracil and histidine supplemented with glucose (2%) with a range of copper, aTc, IPTG, and xylose concentrations, respectively. Three biological replicates were used for all measurements.

**Figure S2.**
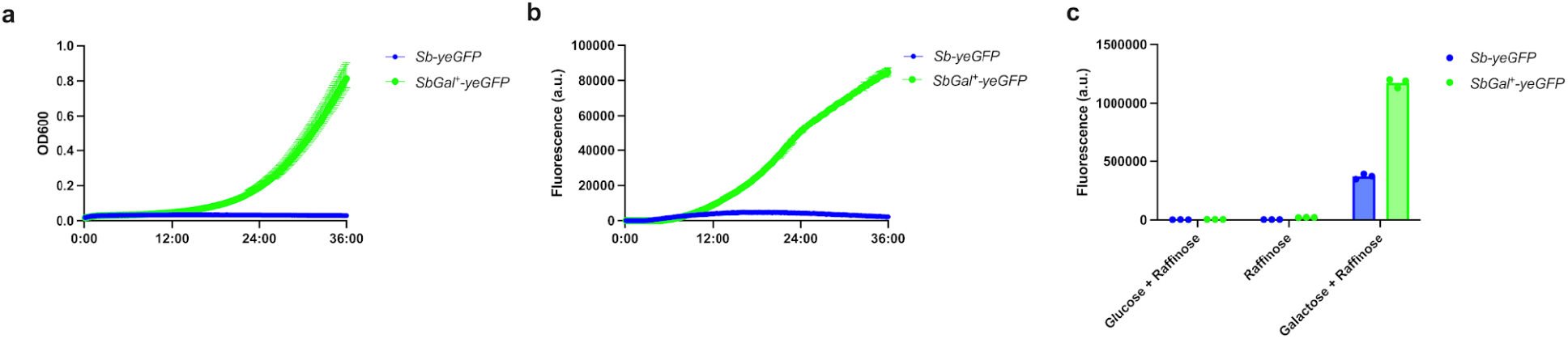
Comparison of growth and expression of yeGFP in *Sb* and *SbGal⁺*. (**a**) Growth of *Sb* and *SbGal⁺* containing *pGAL1-yeGFP* in complete synthetic media without uracil (CSM-U) with galactose. (**b**) Fluorescence levels over time of the strains in (**a**). (**c**) Per-cell fluorescence values (measured via flow cytometry) of both strains grown in glucose+raffinose, raffinose alone, or galactose+raffinose. Three biological replicates were used for all growth curve measurements.

### Inducible systems enable tunable gene expression in an anaerobic environment

We next chose to investigate the performance of *pXYL, pLAC,* and *pGAL* under gut-like (i.e. anaerobic) conditions, selecting these systems due to their favorable dynamic range, low leakiness, and non-toxicity. yeGFP requires oxygen to fluoresce, so we selected the fluorescent protein CaFbFP (*Candida albicans-*adapted flavin-based fluorescent protein) for use as an anaerobic reporter of gene expression (42, 43). Flavin-based fluorescent proteins do not require oxygen to fold (42). We placed *CaFbFP* under the control of the three inducible promoters selected, transformed these plasmids into *SbGal⁺* along with corresponding repressor plasmids as necessary, and cultivated the resulting strains under both aerobic and anaerobic conditions.

All three promoters demonstrated activation under anaerobic conditions (**Fig. S3**). Interestingly, *pLAC* and *pXYL* exhibited lower maximum activation under anaerobic conditions than in aerobic conditions (maximal fluorescence decreased by 78% for both systems), while *pGAL1* demonstrated higher maximum activation (61% increase in maximal fluorescence). The high maximal activation of *pGAL1* in the anaerobic environment, its high dynamic range in the anaerobic environment (169x), as well as the presence of galactose as a major component of human mucins (44), makes *pGAL1* an attractive candidate for applications in the gut.

**Figure S3.**
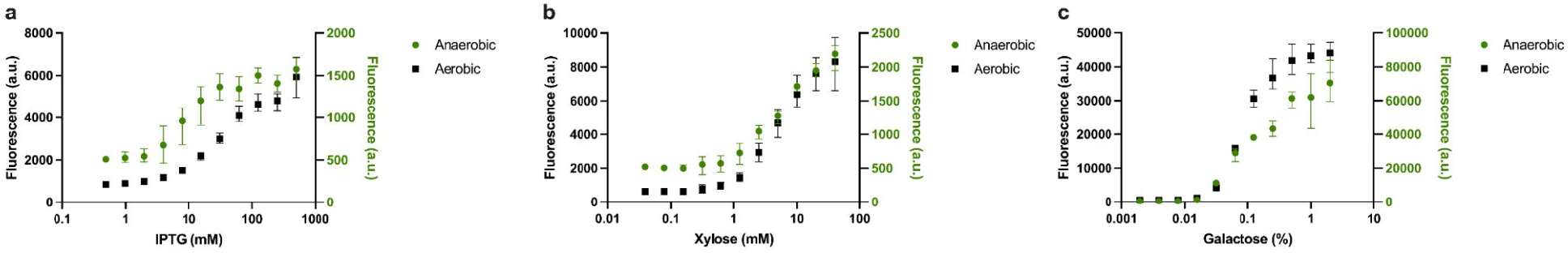
Inducible systems enable tunable gene expression in aerobic and anaerobic environments. Dose-response curves of yeast expressing CaFbfp under aerobic and anaerobic conditions using three inducible systems. Systems with the highest fold induction: *pLAC* (**a**), *pXYL* (**b**), and *pGAL1* (**c**) were cultured under aerobic and anaerobic conditions. (**a**) Strains expressing CaFbfp under the control of the *pLAC* promoter were cultured in synthetic media lacking uracil and histidine supplemented with glucose (2%) and 0.1-1000 mM IPTG. (**b**) Strains expressing CaFbfp under the control of the *pXYL* promoter were cultured in synthetic media lacking uracil and histidine supplemented with glucose (2%) and 0.01-100 mM xylose. (**c**) Strains expressing CaFbfp under the control of the *pGAL1* promoter were cultured in synthetic media lacking uracil supplemented with raffinose (2%) and 0.001-10% galactose. Three biological replicates were cultured for each condition and CaFbfp expression was measured for 24 hours.

Additionally, we observed that the CaFbFP dose-response curves for *pLAC* (under both aerobic and anaerobic conditions) approached saturation with high concentrations of IPTG. This was not observed in the dose-response curve produced using yeGFP (**Fig. 2**). The concentrations and fluorescence signals of flavin-based fluorescent proteins have previously been shown to decrease over time in *E. coli* cells growing exponentially, likely due to a depletion of flavin mononucleotide (FMN) necessary for fluorescence, as well as increased degradation of the proteins by proteases (45). We posit that the “early” saturation we observe in CaFBFP dose-response curves is a consequence of this same mechanism. Nevertheless, this data indicates that *pLAC*, *pXYL*, and *pGAL1* are functional under the anaerobic conditions that are present in the large intestine.

### Inducible systems enable ligand-responsive surface display in *S. boulardii*

Having demonstrated inducible gene expression in *Sb* under anaerobic conditions, we used these systems to enable cell surface display of a therapeutic peptide. Surface display of proteins on the cell surface of commensal bacteria has enabled discovery of host-microbiome interactions and has modulated their colonization/localization within the gut (46). Therefore, enabling surface display in *Sb* will add to its programmability as a therapeutic. In *S. cerevisiae*, surface display is often enabled by fusing the protein of interest to Aga2p (47). Upon expression of the fusion protein, the protein of interest localizes to the cell surface through disulfide bonds that form between Aga1p and Aga2p (48). To investigate the utility of inducible systems for enabling surface display in *Sb*, we engineered *SbGal⁺* to display a short peptide that binds to and inhibits the activity of Toxin A from *Clostridioides difficile* (SA1) (49)*. C. difficile* infections cause colon inflammation and diarrhea and are the most frequently reported nosocomial infection in the United States (50). The ability to inducibly display short peptides that block *C. difficile* toxin activity could enable programmable, targeted treatment and prevention of *C. difficile* infection using *Sb*.

Following the design of *S. cerevisiae* EBY100, a commonly-used yeast strain for surface display, we expressed the cell surface anchor protein Aga1p in the genome via *pGAL1*, and expressed an Aga2p-SA1 fusion protein with a V5 tag under inducible control of *pGAL1* on a low-copy plasmid (**Fig. 3A**). Because surface display of recombinant proteins is often toxic, inducible expression is preferable to constitutive expression because it enables the cell population to grow to the requisite size *in vivo* before initiating surface display. To verify the display of SA1, we exposed induced and uninduced cells to an anti-V5 antibody conjugated with FITC to enable detection of displayed SA1 via flow cytometry and fluorescence microscopy. Cell cultures exposed to galactose demonstrated a significant increase in mean fluorescence compared to uninduced cultures (**Fig. 3B**). Similarly, fluorescence microscopy of uninduced and induced samples exposed to the anti-V5 antibody demonstrated sharp localization of fluorescence to the cell wall, indicating successful display of the SA1 peptide (**Fig. 3C**). No fluorescence above background was detected for uninduced samples. The ability of *Sb* to inducibly display proteins enhances its potential utility as an engineered biotherapeutic.

**Figure 3.**
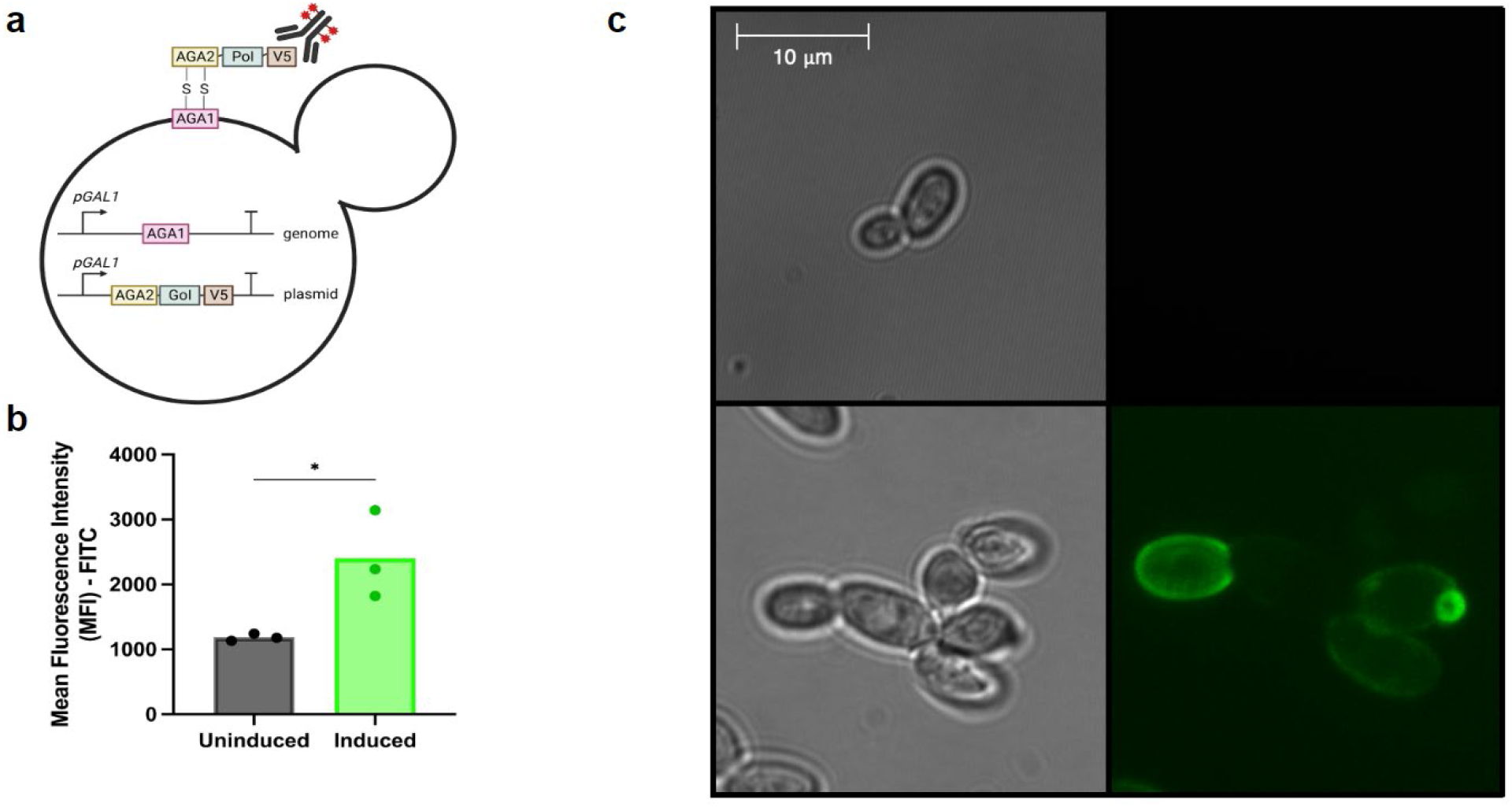
Inducible promoter-mediated surface display in *Sb*. (**a**) Schematic overview of surface display in yeast: *AGA1* is integrated into the genome, while *AGA2* and the Gene of Interest (*GoI*) are placed on a low copy number plasmid, both under the control of the *pGAL1* promoter. Upon exposure to galactose, the Protein of Interest (PoI) is expressed and anchored to the cell wall via disulfide bonds between Aga1p and Aga2p. (**b**) Display of SA1, a short peptide that binds to and inhibits the activity of Toxin A from *Clostridioides difficile* on the *Sb* surface. The expression levels of SA1 on the surface of *Sb* were compared between induced and non-induced conditions using immunoflow cytometry across three biological replicates. (**c**) Confocal microscopy images of *Sb* expressing SA1 without (top) and with (bottom) induction. Left panels show brightfield images, while right panels show fluorescence enabled by an antibody that binds to the V5 tag on SA1. Unpaired t-test was conducted between uninduced and induced groups (*=P< 0.05)

### Inducible systems enable orthogonal gene expression

Engineered microbes can be programmed to make use of multiple inducible promoters, each responding to different inducers, to control the expression of multiple genes. We wished to investigate whether any inducible promoters in our library responded to inducer molecules from other systems, as such “crosstalk” could present a barrier to engineering programmable expression and more complex circuit behaviors. To check for inducer crosstalk, we grew yeast cultures harboring each inducible system in cultures containing high concentrations of each of the five inducer molecules, as well as a no-inducer control. Flow cytometry was used to check for fluorescence in each culture. We found that two inducible systems, the *pGAL1* and *pLAC* system, exhibited significant crosstalk, with galactose inducing a 7.4-fold increase in *pLAC* expression and IPTG producing a 11.2-fold increase in *pGAL1* expression (**Fig. 4A**). This lack of orthogonality between these two systems most likely arises from the chemical similarity between IPTG and galactose. The *pGAL1* system exhibited the highest fold-change (667.6-fold) in expression following induction with the intended ligand, while *pCUP1* had the lowest fold-change (5.5-fold), followed by *pXYL* and *pTET*. This data demonstrates that although the five inducible systems are not fully orthogonal, several sets of two systems exist with minimal crosstalk between them, enabling construction of transcriptional logic gates.

**Figure 4.**
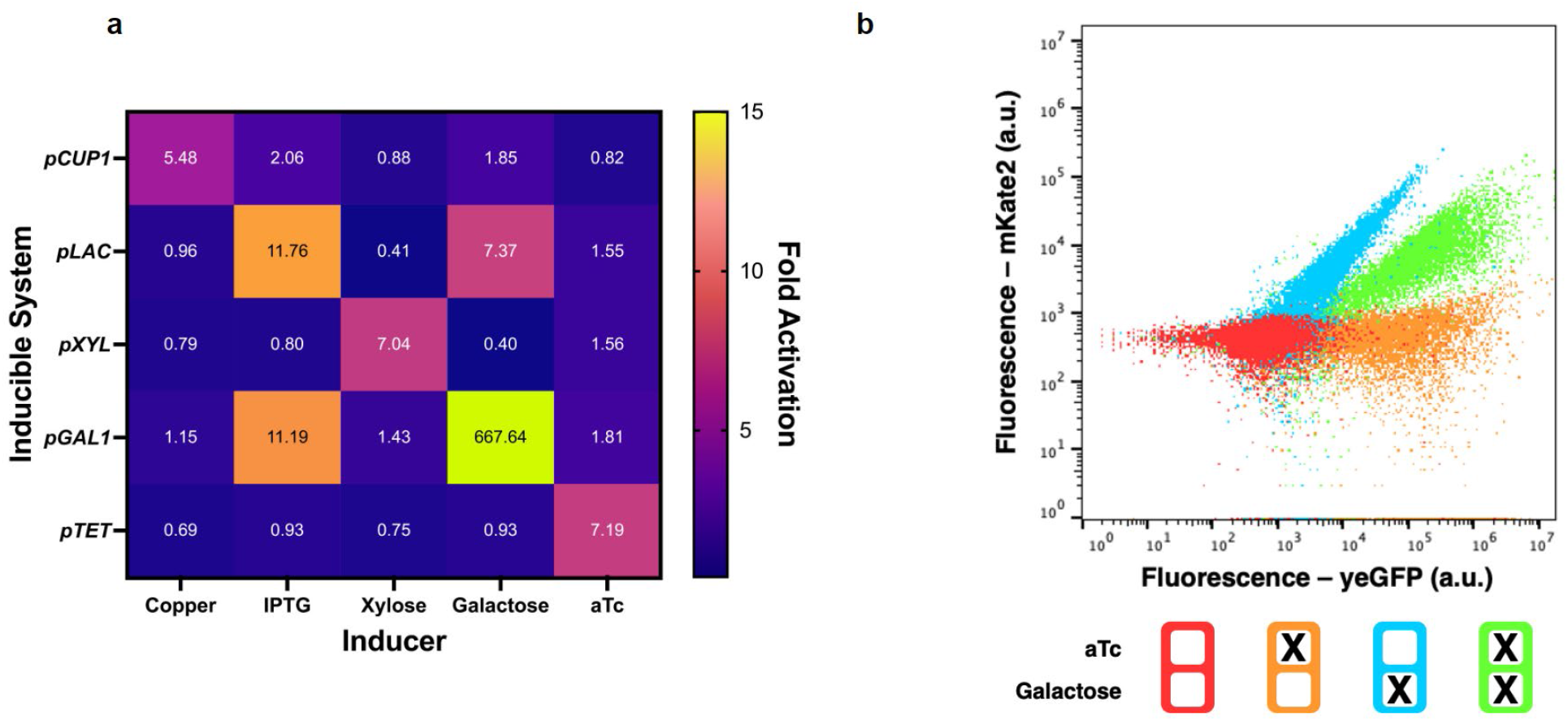
Orthogonality and composability of inducible promoters in *Sb*. (**a**) Evaluating the cross reactivity between the five inducible promoters driving yeGFP expression when exposed to various inducers. Each system was cultivated in CSM media lacking either only uracil or both uracil and histidine, and supplemented with 2% raffinose, in addition to the highest tested concentration of each inducer (as in Figure 2). Color intensity is represented as the average ratio of induced promoter activity to that of the reference (uninduced state). (**b**) A dual-fluorescence experiment demonstrating simultaneous expression of yeGFP and mKate, showcasing the potential of employing multiple inducible promoters within the same system. The orthogonal behavior of pTET and pGAL1 is evidenced by the distinct fluorescence patterns observed under different induction conditions: no inducer (red), aTc only (orange), galactose only (blue), and both aTc and galactose (green).

To further ensure that multiple inducible systems can operate simultaneously within the same cell, we next sought to simultaneously express two different fluorescent proteins (yeGFP and mKate2) using *pTET* and *pGAL1*, which exhibited low levels of crosstalk. yeGFP and mKate2 fluoresce at different wavelengths, enabling them to be detected separately by flow cytometry. We constructed a single plasmid containing *mKate2* under the control of *pGAL1* and *yeGFP* under the control of *pTET*. *TetR*, the cognate repressor of *pTet*, was placed under constitutive control on a separate plasmid. The two plasmids were co-transformed into *SbGal⁺*. To demonstrate simultaneous, independent activation of both inducible systems, the strain was exposed to four conditions (aTc, galactose, both inducers, and no inducers) and fluorescence of mKate2 (controlled by *pGAL1*) and yeGFP (controlled by *pTET*) was measured using flow cytometry. **Fig. 4B** shows a representative scatterplot of the simultaneous and independent activation of two reporter proteins controlled by two inducible systems in the same cell. Because galactose-only induction showed substantial signal in both fluorescence channels, we confirmed that mKate2 emits in both channels via constitutive expression of mKate2 alone, thereby ruling out leaky expression of GFP (**Fig. S4A**). Other pairs of inducible systems are also capable of simultaneous orthogonal induction (e.g. *pTET* and *pCUP1* or *pTET* and *pXYL*), demonstrating that this library of systems can be used to construct more complex circuits.

**Figure S4.**
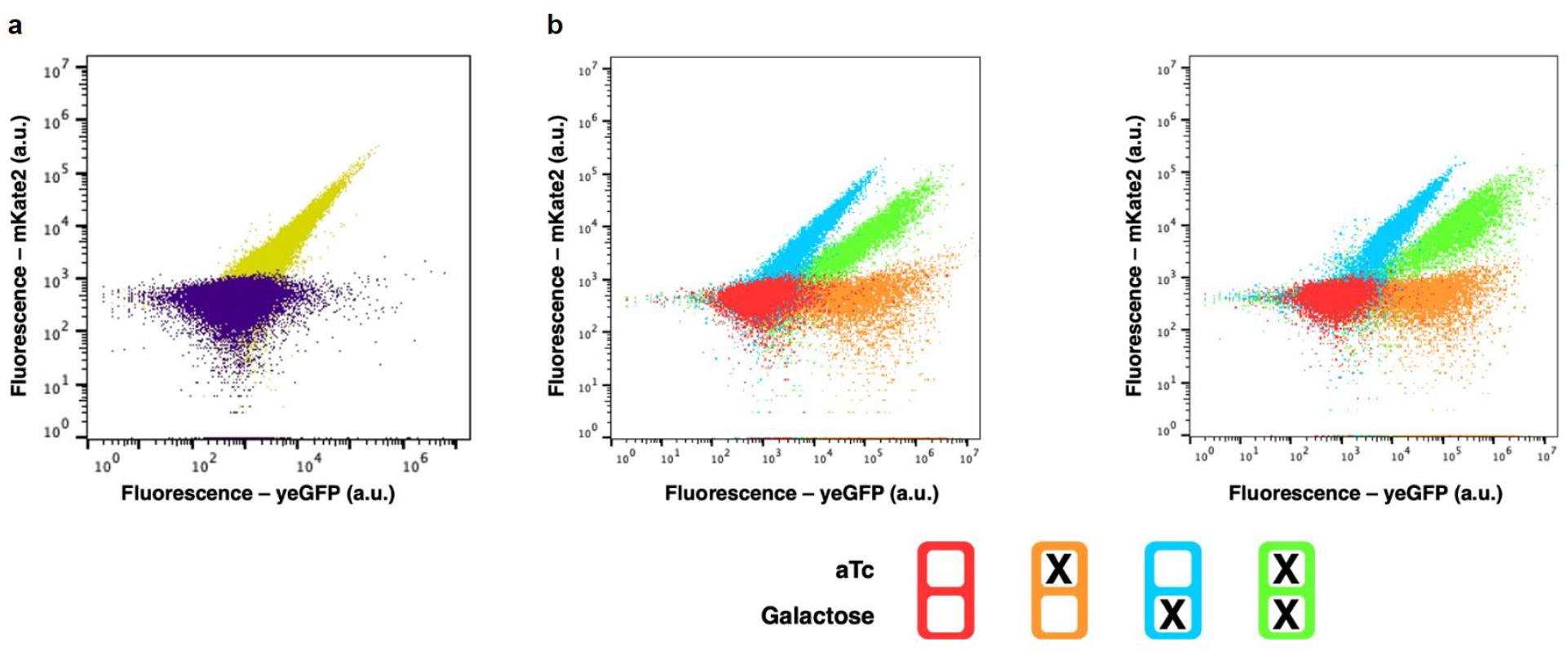
Validation and replication of independent expression of dual fluorescent reporters in *S. boulardii.* (**a**) Scatterplot of *Sb* harboring an empty vector(purple) or a constitutive mKate2 fluorescent reporter (**b**) Replicate scatter plots confirming the results shown in Figure 4.

### Galactose metabolism prolongs gut colonization by *Sb*

Gut mucus is decorated with glycans that serve as a primary carbon source for many intestinal microbes (51). These glycans are primarily composed of galactose, N-acetylglucosamine, N-acetylgalactosamine, fucose, and sialic acid (52). Due to galactose’s presence on intestinal mucus, we hypothesized that the ability to metabolize galactose would provide a colonization advantage to *SbGal⁺* over wild-type *Sb*. To test this hypothesis, we first integrated a nourseothricin resistance marker (*NatR*) to *SbGal⁺* and *Sb* to create strains that can grow on nourseothricin-containing selective plates (which eliminate bacterial and fungi present in fecal samples) (23). The two strains were then delivered to two different groups of antibiotic-treated mice (total of 4 groups). Both groups of mice consumed a noncaloric sweetener sucralose in their water (S), but one of these groups of mice also consumed galactose (S+G). After three consecutive days of *Sb* administration, fecal *Sb* levels were measured every day for one week (**Fig. 5a**). Subsequently, intestinal *Sb* levels were measured at the end of the experiment (day 9). We observed that both *Sb* and *SbGal⁺* attained roughly the same colonization level during the experiment, indicating that intestinal galactose is not sufficient to significantly enhance the colonization of *Sb*. However, upon administration of galactose, *SbGal⁺* maintained an increased abundance by more than three orders of magnitude on day 9, as compared to *Sb* (**Fig. 5b**). The half-life, as measured by the slope of an exponential decay curve, was also significantly higher for *SbGal⁺* than for *Sb* during galactose administration (**Fig. S5**). This increased colonization was also reflected in the intestinal contents, with *SbGal⁺* exhibiting higher levels of colonization of the small intestine, cecum, and colon, as compared to *Sb* during galactose administration (**Fig. 5c**). Additionally, during galactose administration *SbGal⁺* was recovered from the small intestine at levels comparable to the cecum and colon, in contrast to *Sb*, which was absent from the small intestine. These findings agree with those of Liu et al, which showed that galactose is toxic to wild-type *Sb* (38). Small intestinal colonization unveils expanded opportunities for disease treatment beyond those specific to the large intestine, including nutrient provision to the human host. The ability to tune the abundance of *SbGal⁺* using galactose promises enhanced bioavailability and dosage control of delivered therapeutics.

**Figure 5.**
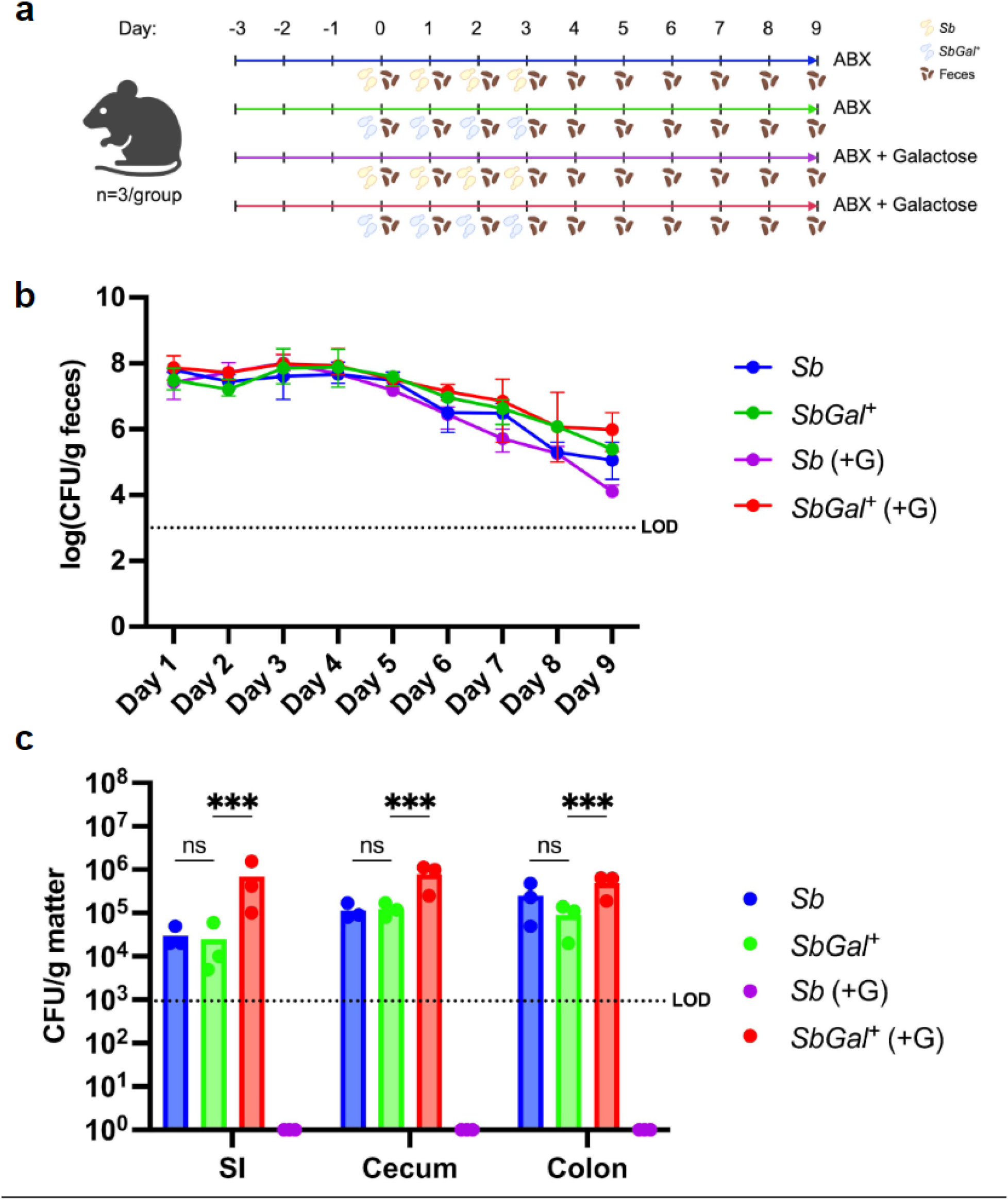
Modulation of *Sb* colonization profile and residence time in the mouse gut via addition of inducing sugars. (**a**) Schematic overview of the mice groups and timeline for antibiotics, galactose and *Sb* administration, and fecal sample collection. (**b**) Colonization profile of wild-type *Sb* and *SbGal⁺* in conventionally-raised mice that were given an antibiotic cocktail (ampicillin (0.5 mg/mL), gentamicin (0.5 mg/mL), metronidazole (0.5 mg/mL), neomycin (0.5 mg/mL), vancomycin (0.25 mg/mL), and sucralose (4 mg/mL)) throughout the experiment. Galactose (2 mg/mL) was administered in water starting from Day 0. 10^8 CFUs of *SbGal⁺* and *Sb* were given to the mice for 4 days. Fecal samples were collected daily and plated on YPD media containing antibiotics. (**c**) Colonization profiles of *Sb* and *SbGal⁺* strains in the small intestine (SI), cecum and colon in this antibiotic treated mouse model. Error bars indicate the SD among the 3 mice in each experimental arm. For each gastrointestinal section datasets, two-way ANOVA with Sidak’s multiple comparisons test was conducted between *Sb*, *SbGal^+^*, Sb (+G) and *SbGal^+^* (+G) treatment groups. (ns P> 0.05, * P< 0.05, ** P< 0.005, *** P< 0.0005)

**Figure S5:**
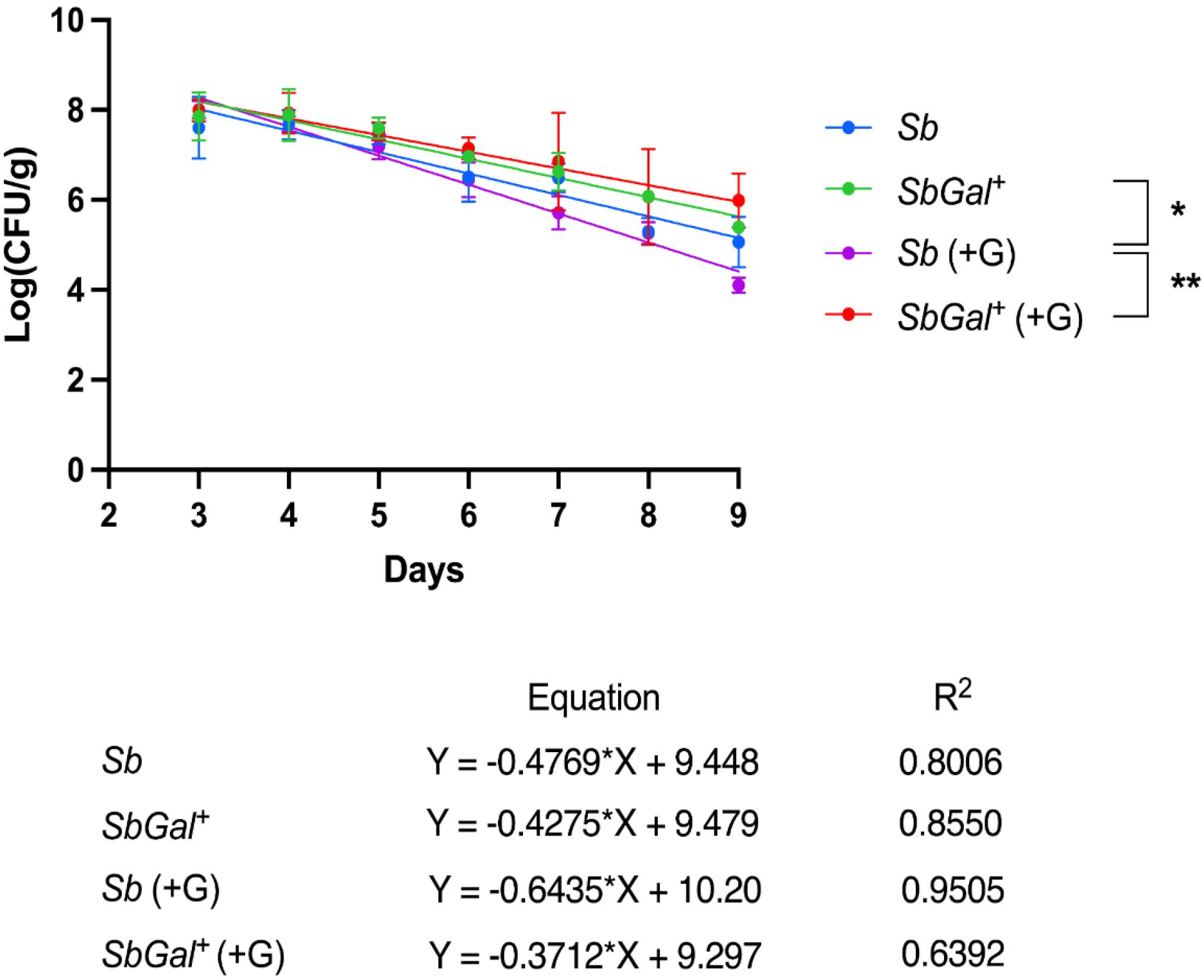
Longitudinal colonization dynamics of *Sb* and *SbGal^+^* in antibiotic-treated mice. This graph displays the average logarithmic counts of colony-forming units per gram (Log(CFU/g)) of fecal samples over time, depicting the colonization profiles of wild-type *Sb* and *SbGal^+^* in the presence and absence of galactose (G). Best fit lines to this logarithmically transformed data are provided below the plot. One-way ANOVA with Sidak’s multiple comparisons test were conducted between the slopes of *Sb*, *SbGal^+^*, Sb (+G) and *SbGal^+^* (+G) treatment groups. (ns P> 0.05, *P< 0.05, **P< 0.005).

### Inducible systems enable on-demand protein production in mouse models

To determine whether the *pGAL1* promoter can be employed to inducibly produce recombinant proteins in the mouse gut using *Sb*, we integrated nanoluciferase (*NanoLuc*) under the control of the *pGAL1* promoter into Site 5 of the *SbGal⁺* genome to produce the strain *SbGal⁺NanoLuc* (23). Nanoluciferase is a small ATP-independent luciferase that produces a luminescent signal upon exposure to furimazine (53). We first found that *SbGal⁺NanoLuc* was able to inducibly produce NanoLuc when cultured in synthetic media (CSM-URA), with maximum luminescence signal detected, in the cultures, 5 minutes after the addition of substrate to the sample (**Fig. S6A-B**). Prior to investigating the behavior of this system in the mouse gut, we investigated how mouse chow would affect *pGAL1* induction, under the suspicion that inducing sugars might be present. We cultured *SbGal⁺* expressing yeGFP from *pGAL1* in a mouse chow diet-derived medium with and without additional galactose. While some activation of *pGAL1* was observed in the chow diet without galactose, yeGFP fluorescence was 15.5-fold higher when the chow diet was supplemented with galactose, demonstrating that inducible activation of *pGAL1* is possible within the chow diet media (**Fig. S6C**).

**Figure S6.**
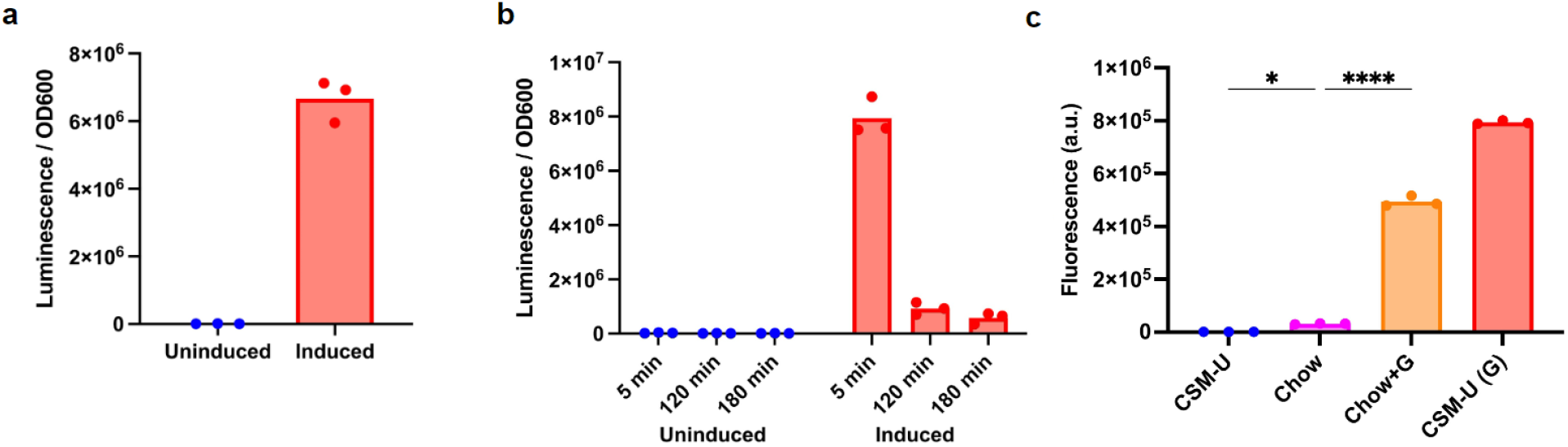
Gene expression in yeast synthetic media and mouse chow diet. (**a**) *Sb* expressing NanoLuciferase under the control of *pGAL1* in CSM-U with and without galactose. (**b**) Measurement of NanoLuciferase luminescence at 5, 120 and 180 minutes after Luciferase substrate addition. Each dot represents a biological replicate. (**c**) Fluorescence from *Sb* expressing yeGFP under the control of *pGAL1* in yeast complete synthetic media without uracil (CSM-U) and chow mice diet (Chow) with and without additional galactose. One-way ANOVA with Sidak’s multiple comparisons test were conducted between CSM-U, Chow and Chow+G groups. (ns P> 0.05, *P< 0.05, **P< 0.005, ***P< 0.0005, ****P< 0.0001)

**Figure S7.**
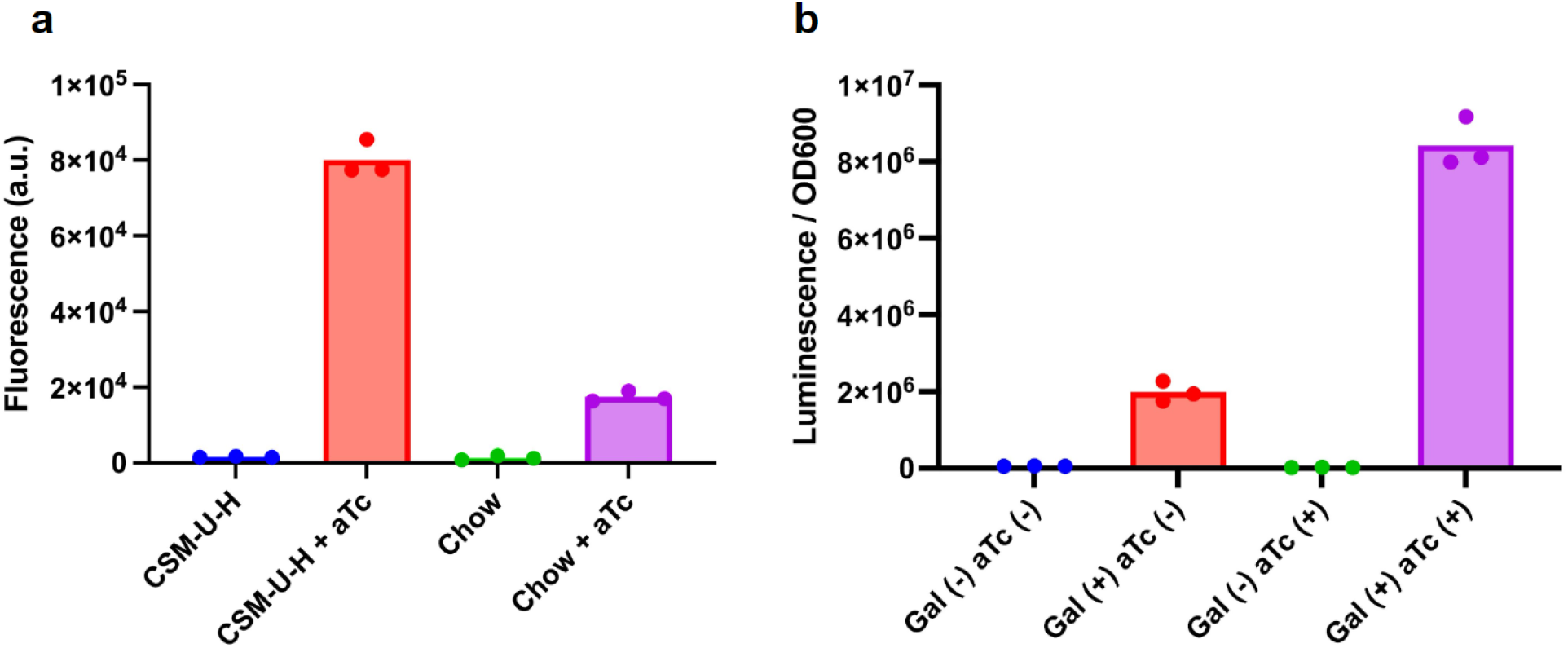
*in vitro* characterization of genetic logic gates in *Sb*. (**a**) *Sb* expressing yeGFP under the control of anhydrotetracycline inducible promoter (*pTET*) was tested in yeast complete synthetic media without uracil and histidine (CSM-U-H) and chow mice diet (Chow) with and without added aTc. (**b**) *Sb* expressing NanoLuciferase under the control of *pGAL-TET* in CSM-U-H in various combinations of inducers. Each dot represents a biological replicate.

Next, we sought to induce production of NanoLuc *in vivo* in the mouse gut using *pGAL1*. We exposed three groups of antibiotic-treated mice (n=4) to various *Sb* strains and sugars (in water) over the course of two days (Day 0 - Day 1) (**Fig. 6A**). The first group of mice received *SbGal⁺* and the non-caloric sweetener sucralose in their water to serve as a control group and provide background luminescence measurements. The remaining two groups both received *SbGal⁺NanoLuc*. One group received sucralose in their water, while the other received both sucralose and galactose as an inducer. Fecal samples were collected daily for two days and NanoLuc luminescence was measured. After two days, mouse intestinal tissues were collected for CFU count assays, NanoLuc activity assays, and imaging of NanoLuc luminescence. All luminescence values were normalized by CFU.

**Figure 6.**
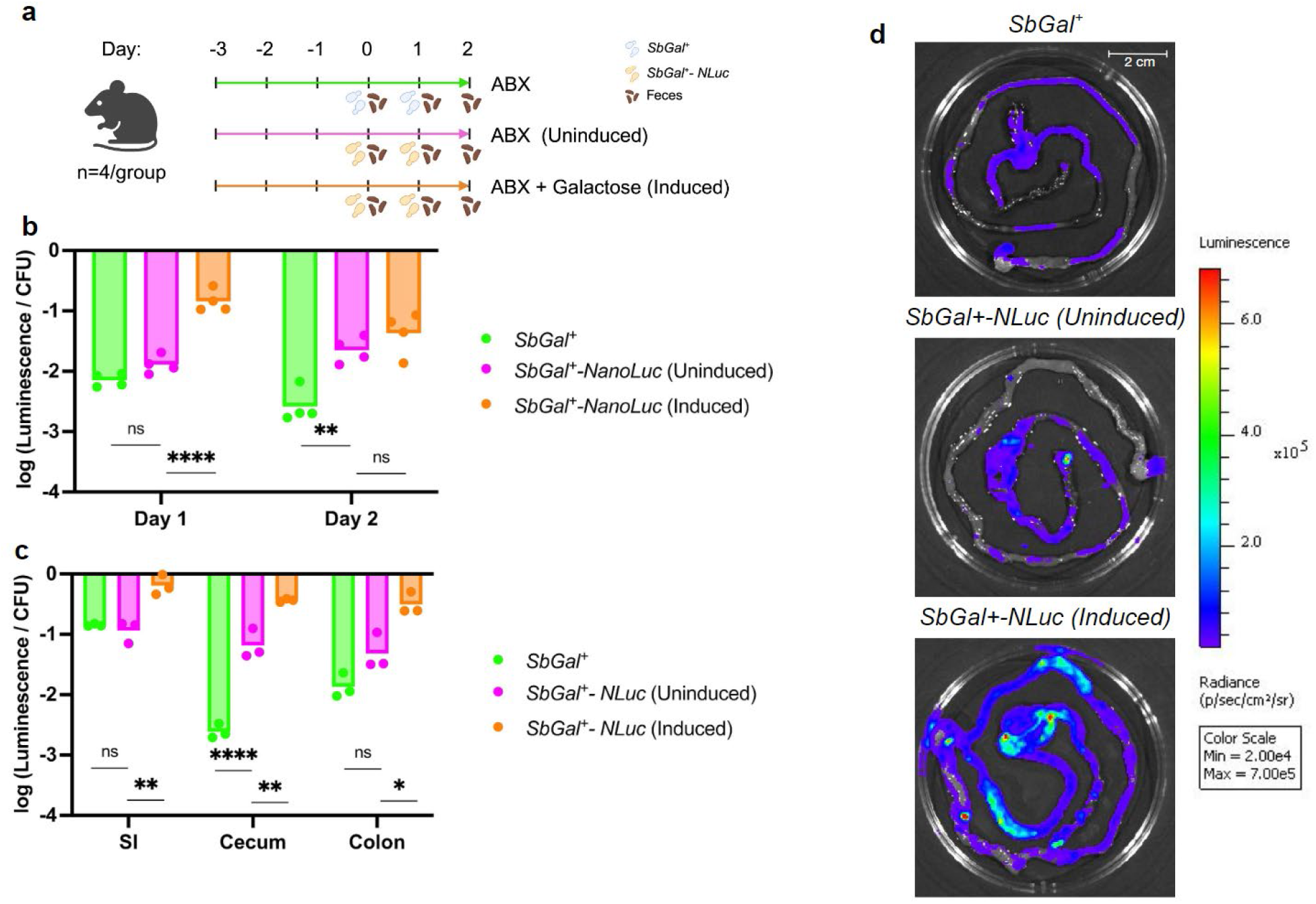
Inducible protein expression in the mammalian gut using *SbGal⁺*. NanoLuciferase expressed via *pGAL1* was used to evaluate *in vivo* protein expression. (**a**) Schematic overview of the mice groups and timeline for antibiotics, galactose and *Sb* administration, and fecal sample collection. (**b**) Detection of NanoLuciferase in fecal matter over days of *SbGal⁺* and galactose treatment. (**c**) Localization and distribution of NanoLuciferase throughout distinct sections of the lower GI tract, including the small intestine (SI), cecum, and colon. (**d**) Luminescence images of NanoLuciferase activity in the GI tract, demonstrating spatially distinct expression patterns across different GI tract regions. Each dot represents one mouse in each experimental arm. For each gastrointestinal section samples and day samples, one-way ANOVA with Sidak’s multiple comparisons test was conducted between *SbGal^+^*, *SbGal^+^* (uninduced) and *SbGal^+^* (induced) treatment groups. (ns P> 0.05, * P< 0.05, ** P< 0.005, *** P< 0.0005, **** P< 0.0001).

Fecal samples from mice treated with the *SbGal⁺NanoLuc* and galactose exhibited higher luminescence than both control groups on both days of the experiment, demonstrating greater levels of production of NanoLuc when galactose is present. Luminescence detected in fecal samples from induced mice bearing *SbGal⁺NanoLuc* was significantly higher than that of uninduced mice bearing *SbGal⁺NanoLuc* on Day 1, demonstrating successful induction of the *pGAL1* system *in vivo* (**Fig. 6B**). However, on Day 2 the luminescence observed from the uninduced condition was not significantly different from the induced condition. This observation, combined with the fact that the uninduced *SbGal⁺NanoLuc* condition enabled significantly higher luminescence than the *SbGal⁺*-only control on both days suggests that galactose present in the gut mucus and mouse diet is sufficient for some degree of activation of *pGAL1*, but that this basal level of activation can be enhanced through supplementation of additional galactose. Similarly, higher luminescence values were detected for the induced *SbGal⁺NanoLuc* condition than for the uninduced *SbGal⁺NanoLuc* condition and *SbGal⁺* control condition for all three locations tested (small intestine, cecum, and colon) (**Fig. 6C**). “Leaky” expression of NanoLuc in the uninduced condition relative to the *SbGal⁺* control condition appeared to be particularly high in the cecum. Bioluminescence imaging of the tissue samples confirmed that the small intestine and cecum exhibited the highest accumulation of NanoLuc upon galactose induction, followed by the colon (**Fig. 6D**). Because whole tissues were immersed in furimazine prior to imaging, the spatial patterns in luminescence we observed likely arise from local inhomogeneities in *SbGal⁺* abundance within the intestines, as “per-cell” luminescence was relatively constant across the gut in the induced condition (**Fig. 6C**). This data shows the spatial distribution of probiotic yeast in the gut at the millimeter scale and indicates that *pGAL1* enables inducible control of recombinant protein production in the mouse gut.

### Inducible systems enable *in vivo* logical operations in the mouse gut

We next sought to demonstrate *in vivo* programmable gene expression in *Sb* by constructing logic gates using two of these inducible systems. We chose aTc and galactose due to their low degree of crosstalk, non-toxicity, and previous use of aTc for activating transcription in the mammalian gut (54, 55). Additionally, the presence of galactose within the gut mucus may serve to restrict engineered gene expression to the gut, thus promoting biocontainment. We constructed a transcriptional AND gate in which both galactose and aTc are necessary for activation of transcription, by cloning two Tet operators (separated by a T-rich spacer) downstream of the *pGAL1* promoter, which contains an operator for the Gal4p activator. The spacer separates the operators to prevent steric competition between repressor and activator proteins, and the high T content facilitates nucleosome depletion, enabling polymerase access to the promoter region (41). In the absence of aTc, the TetR repressor is bound to the Tet operator sites, preventing readthrough, while in the absence of galactose, the Gal4p activator is absent, preventing transcription. Only when both inducers are present should transcription of downstream genes occur. Nanoluciferase (NanoLuc) was selected as the reporter for *pGalTet* activity. A plasmid containing the AND-gate NanoLuc unit was transformed to *SbGal⁺* along with a second plasmid encoding the corresponding TetR repressor to produce the strain *SbGal⁺AND*.

We first tested the behavior of the *pTET* promoter alone as well as the *pGalTet* transcriptional logic gate *in vitro* in both synthetic media and in mouse chow media. We found that *pTET* is functional in chow media, although the maximum fluorescence signal obtained from a *pTET-yeGFP* reporter was lower in chow media than in synthetic media (**Fig. S6A**). Having demonstrated *pTET* functionality in chow media, we next examined the behavior of the *pGalTet* transcriptional logic gate driving production of NanoLuc in synthetic media. We found high NanoLuc production when both inducers are present compared to lower production when only galactose is present, and no luminescence above background was observed under the “aTc only” and “no inducer” conditions (**Fig. S6B**). This behavior demonstrates that *pGalTet* is not a perfect AND gate, as some transcriptional activation occurs in the presence of galactose alone. Nevertheless, this *in vitro* characterization indicated that *SbGal⁺AND* would function predictably for *in vivo* experiments.

To examine the logic behavior of *SbGal⁺AND* in the mammalian gut, four groups of antibiotic-treated mice (n=4) were treated with *SbGal⁺AND* for 3 days (**Fig. 7A**). Starting on Day 0, each group of mice received either no inducer, galactose only, aTc only, or both galactose and aTc in their water. As before, fecal samples were collected daily for two days, after which samples of GI contents were taken. Fecal samples from mice exposed to at least one inducer demonstrated significantly higher NanoLuc production than mice exposed to no inducers, confirming that the logic gate does not exhibit pure *AND* behavior (**Fig. 7B**). However, when mice are treated with both inducers, we observed a more significant increase in luminescence. A similar relationship was observed for cells retrieved from the mouse GI tract (**Fig. 7C**). The synergistic interaction of both galactose and aTc was pronounced, with the luminescent output in the dual-inducer condition exceeding the cumulative effects of individual inducers. This experiment highlights the successful *in vivo* use of the AND gate logic system in probiotic yeast.

**Figure 7.**
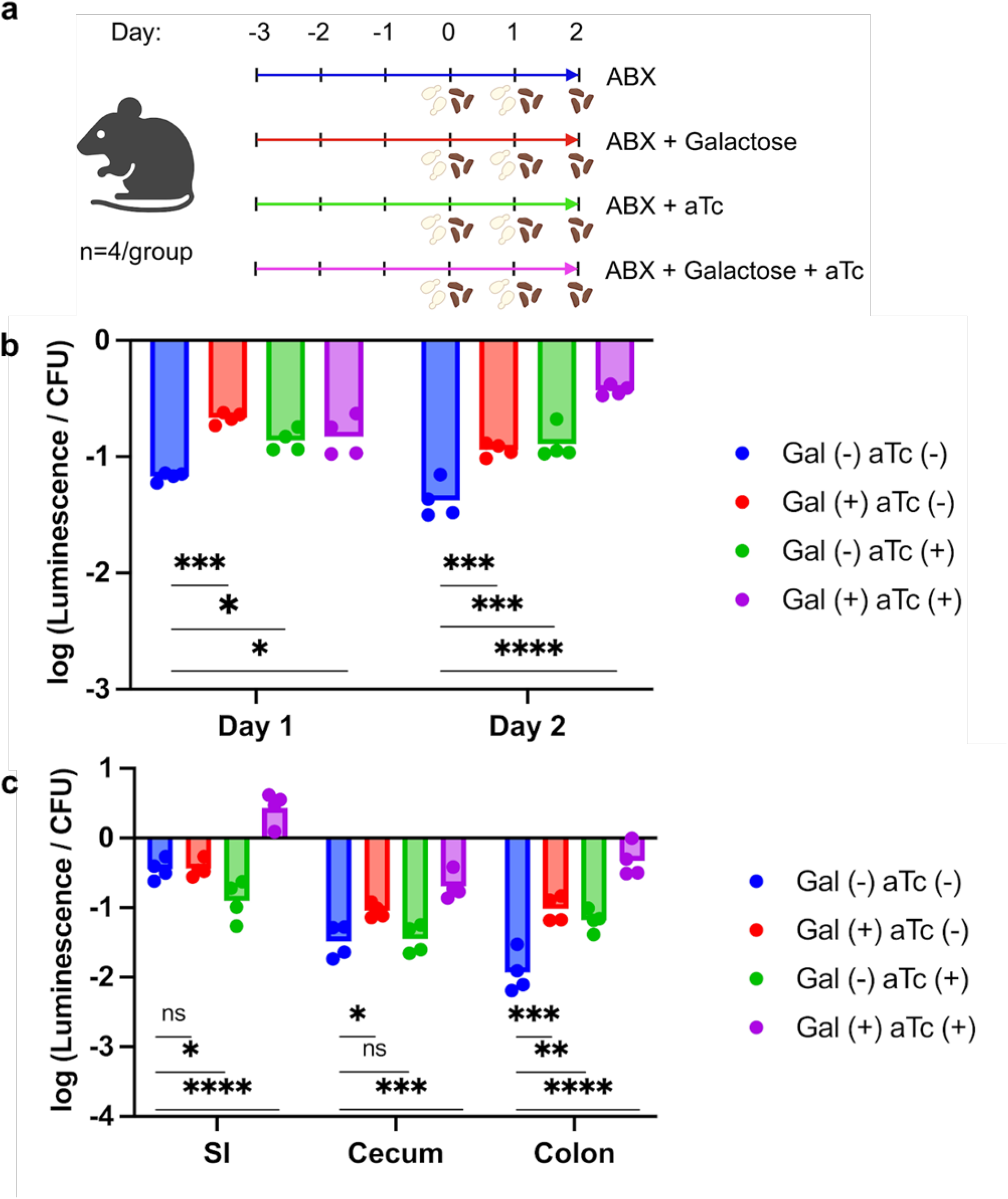
Development and validation of genetic logic gates for precision control of protein expression in the gut. (**a**) Schematic overview of the mouse groups and timeline for antibiotics, inducer and *Sb* administration, and fecal sample collection. (**b**) Comparative evaluation of NanoLuciferase activity under these four conditions in the mouse gut over two days, showing the functionality of the genetic logic gate *in vivo*. (**c**) NanoLuciferase expression across distinct sections of the lower GI tract, including the small intestine (SI), cecum, and colon, signifying the functionality of this gate throughout the gut. For each gastrointestinal section samples and day samples, one-way ANOVA with Sidak’s multiple comparisons test was conducted between SbGal+, SbGal+ (uninduced) and SbGal+ (induced) treatment groups. (ns P> 0.05, * P< 0.05, ** P< 0.005, *** P< 0.0005, **** P< 0.0001)

## Conclusion

Unlike nonliving therapeutics, engineered live biotherapeutics are subject to ecological forces that impact their abundance and product expression level after administration. Because the taxonomic and metabolomic composition of the gut can vary substantially between individuals, a lack of external control of recombinant gene expression and available nutrients can negatively impact the efficacy and safety of probiotic therapies. Here, we show that several inducible gene expression systems that are sensitive to nontoxic small molecules are suitable for controlled gene expression within *S. boulardii*, and that at least two of these (*pGAL1* and galactose, *pTET* and anhydrotetracycline) are functional in a live mammalian host. This inducible construct allowed us to visualize the local biodistribution of *S. boulardii* within the intestines, as its fitness prior to visualization was not perturbed by high-level expression of a recombinant gene. Further work is necessary to determine whether local inhomogeneities in *Sb* abundance are persistent or dynamic in the face of intestinal peristalsis. This galactose-enhanced colonization invites further exploration into modulation of probiotic metabolism to exploit host-provided nutrients.

We also observed a low level of *pGAL1* activation *in vivo*, even without the addition of galactose. We rationalize this observation based on the known presence of galactose within mucus polysaccharides, as well as the presence of gut bacteria (which are known to be present in all antibiotic-treated mouse models) which may liberate galactose monosaccharides from the mucus. A testable hypothesis based on this observation is that *pGAL1* would exhibit reduced leaky expression in monocolonized mice, or in mice without mucus-degrading bacteria.

We showed that several of these inducible systems are composable into higher-order logical functions, such as AND, thus enabling even more control over *in situ* therapeutic biomanufacturing. We envision that these systems could be used to activate therapeutic production (via co-administration of inducer) at the onset of treatment, followed by cessation of inducer once symptoms have resolved, thereby providing tighter control over drug dose than provided by intestinal washout alone. However, the timescale of protein induction *in vivo* remains to be measured and potentially shortened. Looking forward, we expect that the use of inducible systems with very low “off” states, coupled with biosensors that can recognize therapeutically relevant biomarkers (57), will enable tightly-controlled, autonomous disease treatment using engineered live biotherapeutics.

Serendipitously, we found that expanding *S. boulardii*’s galactose metabolism, which was necessary to enable simultaneous growth and gene expression, also conferred a colonization advantage in the mouse gut during inducer administration. While *pGAL1* seemed to be slightly activated during gut passage in the absence of additional galactose, the high sensitivity of *pGAL1* implies that naturally bioavailable galactose within the intestines is insufficient to provide a meaningful colonization advantage to *SbGal⁺*. However, further experiments are necessary to determine whether this effect is reproducible in mice that have not been treated with antibiotics, and in more human-like gut models. Taken together, these results imply that galactose can provide a significant level of control over the gut residence times of both *SbGal⁺* (which can utilize galactose as a carbon source) and *Sb* (for which galactose is toxic).

Taken together, the incorporation and *in vivo* functionality of inducible gene expression systems enable precision control over dosage that *Sb* provides, as well as the location within the gut that the dose is administered. In tandem with *Sb’s* capability for synthesizing small molecules, secreting proteins, or displaying proteins on its cell surface, tunable gene expression advances *S. boulardii*’s utility for therapeutic applications and studies of host-microbiome interactions.

## Materials and Methods

### Strains and Culture Media

*Escherichia coli* NEB Stable, NEB 5ɑ, and NEB 10β were used for plasmid construction and maintenance. *E. coli* cells were grown in lysogeny broth (LB) (5g/L yeast extract, 10 g/L tryptone, 10g/L NaCl) (at 37 °C, 250 rpm) supplemented with ampicillin (100 µg/mL), kanamycin (50 µg/mL) or chloramphenicol (34 µg/mL). *Saccharomyces boulardii* ATCC-MYA796*ΔURA3ΔHIS3* was used to construct *SbGal*⁺, and *SbGal⁺* was used for subsequent inducible promoter characterization, yeast surface display, and logic gate experiments. *Sb* strain ATCC-MYA797 was used as a control for growth characterization experiments. *Saccharomyces cerevisiae* strain BY4741 was the source for the *PGM2* gene and was used as a control for growth characterization experiments. Yeast cultures for genome editing were grown in yeast extract-peptone-dextrose (YPD) medium (50 g/L YPD Broth (Sigma-Aldrich)). For all other experiments, yeast cultures were grown in synthetic complete media containing 0.67% (w/v) Yeast Nitrogen Base Without Amino Acids (Sigma-Aldrich), 1.92 g/L Yeast Synthetic Media Dropout Mix (uracil, histidine, or both), and glucose (2% (w/v)) or raffinose (2% (w/v)) as a carbon source unless otherwise indicated. All *Sb* strains were grown at 37 °C and 250 rpm and all *S. cerevisiae* strains were grown at 30 °C and 250 rpm. Anaerobic cultivations were carried out in an anaerobic chamber (Coy lab, gas mix 90% Nitrogen, 5% carbon dioxide, 5% hydrogen) at 37 °C with agitation at 900 rpm provided by BioShake iQ.

### Plasmid and Strain Construction

A list of the strains, gene fragments, and primers used to collect the data presented in this work is shown in **SI Appendix, Tables S1-S3**, while detailed sequences and maps are presented in **SI Appendix, Datasets S1-S3**. *SbGal⁺* was constructed by replacing the *Sb PGM2* gene with the *S. cerevisiae* version via CRISPR-Cas9 genome editing. Plasmid ISA1041 provided guide RNA and Cas9 nuclease to carry the edit. A 300 bp fragment of *S. cerevisiae PGM2* gene containing the position to be corrected was amplified via PCR and co-transformed with ISA1041 to wild-type *Sb* as described below. Selection was performed by growing transformants in synthetic media containing galactose as the sole carbon source. Editing of *PGM2* was confirmed by Sanger Sequencing.

A synthetic toolkit (MoClo-YTK) containing yeast parts were gifts from the Dueber Lab (Addgene #1000000061). Expression vectors for *yeGFP*, *mKate*, and *CaFbFP,* assembled via Golden Gate Assembly, consisted of two connectors, an inducible or constitutive promoter, the fluorescent protein coding sequence, the *tENO1* terminator, the *URA3* yeast marker, the *2µ* yeast origin, and *AmpR/ColE1* as an *E. coli* marker and origin. Similarly, expression vectors for cognate repressor proteins included two connectors, the constitutive promoter *pFBAI* (41), the repressor protein coding sequence, the *tENO2* terminator, the *HIS3* yeast marker, the *CEN* yeast origin, and *AmpR/ColE1* as an *E. coli* marker and origin. All yeast parts were included in the MoClo-YKT kit except for the following parts, which were ordered as gBlocks from Integrated DNA Technologies, Inc (IDT) with appropriate type-2 restriction sites and overhangs for Golden Gate Assembly: *pFBAI, tetR, lacI, xylR, CaFbFP,* and *mKate2.* The following parts were ordered as plasmids from IDT (Gene Synthesis): *pTET, pLAC, pXYL.* Expression vectors were assembled according to Deuber lab YTK protocols via Golden Gate cloning, with the Golden Gate reaction mixture containing 0.5 µL of 40 nM of each DNA part (20 fmol), 0.5 µL T7 ligase (EB), 1.0 µL T4 Ligase Buffer (NEB), and 0.5 µL BsaI (10,000 U/mL, NEB), with water to bring the final volume to 10 µL. Assembly was performed on a thermocycler using the following program: 30 cycles of digestion ( 37 °C for 2 min) and ligation (16 °C for 5 min), followed by a final digestion (60 °C for 10 min) and heat inactivation (80 °C for 10 min).

To construct condensed plasmids for orthogonality experiments, Gibson Assembly was used to assemble both the inducible promoter-fluorescent protein transcriptional unit and constitutive promoter-cognate repressor transcriptional unit into the same backbone, separated by a connector. For each condensed plasmid, three fragments were amplified, consisting of the two transcriptional units and yeast backbone with *E. coli* marker and origin, with ∼20 bp homology between fragments. The Gibson Assembly mixture consisted of the following: 100 ng backbone fragment, additional insert fragments in 2:1 molar ratio to backbone fragment, 10 µL HiFi 2X Master Mix (NEB), and water up to 20 µL. The reactions were incubated in a thermocycler at 50 °C for 30 min prior to transformation to *E. coli*.

The pYD1 plasmid was a gift from the Wittrup Lab (Addgene #73447) and the *TRP1* marker on pYD1 was swapped with *HIS3* marker from MoClo-YTK via 2-part Gibson cloning, resulting in DD580. Then, SA1 was ordered as primers and was inserted into MCS on DD580 via Q5 mutagenesis (DD608).

The pGAL1-AGA1-URA3 integration cassette (DD579) was constructed via Golden Gate assembly with the ISA086 backbone. AGA1 was amplified from the *Sb* genome and pGAL1 and tENO1 were amplified from the YTK.

The nourseothricin resistance gene *natR* was obtained from pYTK078 and assembled into an integration cassette (ISA186) by Golden Gate assembly as described above. The resulting integration cassette provided the DNA repair template and was transformed with CRISPR-Cas9 and gRNA targeting integration site 1 (23) (ISA1045) for CRISPR-based insertion of *natR* into the *Sb* MYA 796 ΔURA ΔHIS and *SbGal⁺* genomes, producing Sb*::NatR* and *SbGal⁺::NatR,* respectively. Transformation reactions were plated in YPD supplemented with nourseothricin and positive integrations were screened via Sanger sequencing.

Integration cassettes for logic gate constructs were constructed via Golden Gate assembly of the following parts: The integration cassette backbone, the logic gate, NanoLuc, and a transcriptional terminator. The pGalTet logic gate and the NanoLuc gene were both ordered as gBlocks with BsaI overhangs. The resulting integration cassette provided the DNA repair template and was transformed with CRISPR-Cas9 and gRNA targeting integration site 5 (23) for CRISPR-based insertion of *natR* into the *Sb* MYA 796 ΔURA ΔHIS and *SbGal⁺* genomes.

### Yeast Competent Cells and Transformations

We used the yeast competent cell preparation and transformation protocol from Durmusoglu et. al (2021) (23), which is based on the protocol from Gietz et. al (56, 57). To prepare competent cells, yeast colonies were inoculated into 1 mL YPD and incubated in a shaking incubator overnight at 37 °C, 250 rpm. This culture was diluted into fresh 25 ml YPD (with OD600 ≅ 0.25) and grown to OD 0.8-1.0. Cells were pelleted by centrifugation for 5 minutes at 3,000 x g and resuspended in 25 ml autoclaved water before being centrifuged again under the same conditions. The cells were then resuspended in 1 mL lithium acetate (100 mM, Sigma-Aldrich) and centrifuged again under the same conditions. The cells were resuspended in 250 µL lithium acetate (100 mM) and divided into transformation tubes, with 50 µL/tube. Cells were washed again in 1 mL lithium acetate (100 mM) before being spun down and the supernatant removed. The cell pellet was then gently resuspended in 50 µL boiled salmon sperm DNA (2 mg/ml), and transformation reagents were added in the following order: 2 µg DNA repair template if applicable, 1 µg of any yeast plasmids (either for expression or for gRNA and Cas12a expression), 36 µL lithium acetate (1.0 M), and 260 µL PEG3350 (50%, Fisher Scientific). To produce the salmon sperm DNA, double-stranded salmon sperm DNA (Invitrogen, 15632011) at 10 mg/ml was diluted to 2 mg/mL and incubated at 95 °C for 5 min to denature the DNA. The transformation mix was gently vortexed for less than 5 seconds at low speed before being heat shocked at 42 °C for one hour. The transformation reactions were then centrifuged for 3 minutes at 3,000 x g, and the supernatant was removed and discarded. The cells were resuspended in 1 mL YPD by gentle pipetting, and recovered for 1 hour at 37 °C, 250 rpm (or 30 °C for *S. cerevisiae*). In the case of genome editing reactions, this recovery period was extended to a total of 3 hours. Finally, the cell suspension was centrifuged for 1 minute at 3,000 x g and the pellet was resuspended in 100 µL. 50 µL of the suspension was plated on appropriate growth media.

### S. boulardii Colony PCR

Yeast genome edits were confirmed using Phire Plant Direct PCR Master Mix from Thermo Fisher. The protocol for performing PCR amplification directly from yeast colonies is described by the manufacturer. Briefly, 10 µL of master mix was combined with 1 µL of each primer and water up to 20 µL. Using a pipette tip, a small part of a yeast colony was picked and resuspended in the PCR reaction. If the PCR failed, a modified protocol was used, as follows. A small colony was resuspended in 8 µL of 20 mM NaOH and incubated at 98 °C for 10 minutes. Then 10 µL of PCR master mix was added to the lysed cells, combined with 1 µL of each primer. The PCR reaction then proceeded according to the supplier’s specifications. Primers were designed to bind outside the linear repair template’s homology arms.

### Construction of Growth Curves

Three biological replicates of each strain were grown overnight at 30°C or 37 °C, 250 rpm in their corresponding media. The cultures were then subinoculated to OD 0.1 in 96-well-plates (Costar, Corning™ 3788) in appropriate media (single and combination carbon sources) and grown for 36 hours in a plate reader (BioTek Synergy™ H1, Shake Mode: Double Orbital, Orbital Frequency: continuous shake 365 rpm, Interval: 10 min).

### Flow Cytometry Experiments

#### Dose-Response Curve Construction

Yeast strains were inoculated from single colonies on plates into 1 mL of appropriate media and grown overnight at 37 °C, 250 rpm. Cultures were then subinoculated to OD 0.1 in media containing any appropriate inducer molecules. Cultures were induced for 24 hr.s and incubated at 4 °C for 1-2 hours to facilitate protein folding (anaerobic cultures were not incubated at 4 °C). Anaerobic cultures were incubated in the anaerobic chamber. The *pCUP1*-yeGFP strain was induced in sulfate free media to reduce non-specific induction. The *pGAL1-yeGFP* strain was induced in media with raffinose as the carbon source to eliminate repression of promoter activity glucose. Both aerobic and anaerobic cultures were then diluted to OD 0.1-0.5 in flat-bottom 96-well plates and run on a BD Accuri™ C6 Plus Flow Cytometer. For each replicate, 10,000 events were collected under settings of FSCH-H < 20,000 and SSC-H < 600 and medium to low flow (500-2000 events/seconds). Fluorescence was detected on the FITC channel for yeGFP and CaFbFP. No gating was performed.

#### Orthogonality

Yeast strains were inoculated from single colonies on plates into 1 mL of appropriate media and grown overnight at 37 °C, 250 rpm. Cultures were then subinoculated to OD 0.1 in 96-well deep well plate wells with media containing each of the inducer molecules at their highest concentration. Cultures were induced for 24 hours and incubated at 4 °C for 1-2 hours to facilitate protein folding. Cultures were then diluted to OD 0.1-0.5 in flat-bottom 96-well plates and run on a BD Accuri™ C6 Plus Flow Cytometer. For each replicate, 10,000 events were collected under settings of FSCH-H < 20,000 and SSC-H < 600 and medium to low flow (500-2000 events/seconds). yeGFP fluorescence was detected on the FITC channel and mKate2 fluorescence was detected on the PerCP channel. No gating was performed.

#### Induced surface display detection

Yeast strains were inoculated from single colonies on plates into 1 mL of CSM-U media and grown overnight at 37 °C, 250 rpm. Cells were subinoculated to OD 0.5 in 5 mL CSM-U media with 1.8% galactose and 0.2% glucose and cultured 16 hours at 30 °C to induce the surface display of peptide. 5×10^6^ cells were processed for flow cytometry analysis. After media removal, cells were washed in 0.1% BSA 1X PBS. Then, they were labeled with 200 μL anti-V5 Antibody, FITC (1:250) at 800 rpm rotation at 4°C. Cells were collected and washed in 0.1% BSA 1X PBS and resuspended in 200 µL PBS and run on a BD Accuri™ C6 Plus Flow Cytometer in 96-well plate format. For each replicate, 10,000 events were collected under settings of FSCH-H < 20,000 and SSC-H < 600 and medium flow (2000 events/seconds). Fluorescence was detected on the FITC channel. No gating was performed.

### Microscopy Imaging

Yeast suspensions were prepared by pipetting 5 µL onto a glass slide. Another glass slide coverslip was placed immediately on top of the yeast suspension droplets to create a thin film for imaging and avoid evaporation while imaging. All images were acquired using an inverted Leica DMi8 microscope with a 63x oil-immersion objective (NA = 1.40) and a Hamamatsu Orca-Fusion camera with a 60 ms exposure time. Spinning disk confocal microscopy and imaging was accomplished by using an 89 North LDI-7 Laser Diode Illuminator at 470 nm at 20% power for excitation and 510 nm for emission. When conducting confocal microscopy, maximum intensity projection images were acquired using 10 slices with a 1 µm step size. Brightfield images used a single imaging plane. Confocal and brightfield images were obtained sequentially.

### Chow/Mouse diet experiment

100g of chow oval pellets (Laboratory Rodent Diet catalog #001319) was resuspended in 1 L of DI water and filtered through a 0.22 µL filter (Catalog number: 567-0020). Filtered supernatant was used to grow Sb MYA 796 ΔURA ΔHIS and *SbGal⁺* with or without NanoLuciferase constructs. Expression of NanoLuciferase was measured in chow diet growth media with or without 2% galactose.

### Nanoluciferase activity detection (*in vitro*)

Nanoluciferase expressing strains were inoculated in triplicates from single colonies on plates into 1 mL of complete synthetic media (CSM) lacking uracil (-URA) (for *pGAL*) or complete synthetic media (CSM) lacking uracil and histidine (-URA-HIS) (for AND-gate) and grown overnight at 37 °C, 250 rpm. Cultures were then subinoculated to OD 0.1 in 1 mL CSM-URA or CSM-URA-HIS with respective inducers (galactose only for *pGAL1,* and either galactose only, aTc only, or both inducers for *pGalTet-NanoLuc*) and grown for 24 hours at 37 °C, 250 rpm.10 µl of each culture was collected for the luciferase assay (Promega, Nano-Glo Luciferase Assay System). Cells were diluted in 90 µl media, optical densities were measured in a plate reader (BioTek Synergy™ H1, absorbance 600 nm) and then mixed with 100 µl NanoGlo Assay Reagent (10 µl Nano-Glo Luciferase Assay Substrate and 90 µl sterile 1X PBS) in a 96 well black opaque plate. Cells were incubated for 5 minutes for luminescent reaction to take place. Luminescence was measured in a plate reader (BioTek Synergy™ H1, emission 420 nm). Luminescence values were normalized by OD600 values for each replicate.

### Mouse Experiments

All mouse experiments were approved by the NC State University Institutional Animal Care and Use Committee (IACUC).

#### *Sb* and *SbGal⁺* Experiments

Six-week old female C57BL/6J mice were obtained from Jackson Laboratories and hosted at the NCSU Biological Resources Facility (BRF) for 3−4 days before experiments. Mice were housed in groups of three and their cages were changed before treatment with antibiotic cocktail, sugar administration and before treatment with *Sb* or *SbGal⁺*. Groups with galactose administration were administered with galactose (20 mg/mL) *ad libitum* in filter sterilized drinking water starting from day 0 and refreshed daily. Antibiotic administration was started 3 days prior to the yeast gavage and continued during the experiment and refreshed daily. Antibiotic cocktail consisted of ampicillin (0.5 mg/mL), gentamicin (0.5 mg/mL), metronidazole (0.5 mg/mL), neomycin (0.5 mg/mL), vancomycin (0.25 mg/mL) and sucralose (4 mg/mL). Mice were gavaged with 10^8^ CFU *Sb* ( *SbGal⁺* or *Sb*) every day for 4 days (D0, D1, D2, D3). Fecal samples were collected every 24 h from day 1 to day 9. On Day 9, the mice were sacrificed and intestinal contents (small intestine, cecum, colon) were collected.

1−2 pieces of stool or intestinal matter were collected in preweighed 1.5 mL centrifuge tubes and then weighed again to determine fecal mass. Fecal matter was then resuspended in 1 mL PBS per 10 mg feces. Fecal suspensions were plated on YPD media containing 50 μg/mL nourseothricin and 0.25 mg/mL streptomycin. Plates were sealed with parafilm and incubated at 37 °C for 2−3 days.

#### SbGal⁺-NLuc Experiments

Six-week old female C57BL/6J mice were obtained from Jackson Laboratories and hosted at the NCSU Biological Resources Facility (BRF) for 3−4 days before experiments. Mice were housed in groups of four and their cages were changed before treatment with antibiotic cocktail, inducer administration and before treatment with *SbGal⁺* or *SbGal⁺-NLuc*. Antibiotic cocktail (ampicillin (0.5 mg/mL), gentamicin (0.5 mg/mL), metronidazole (0.5 mg/mL), neomycin (0.5 mg/mL), vancomycin (0.25 mg/mL)), sucralose (4 mg/mL) was administered *ad libitum* in filter sterilized drinking water starting from 3 days prior to *Sb* gavage and continued throughout the experiment and refreshed daily. Galactose induction (20 mg/mL) was administered via *ad libitum* in filter sterilized drinking water starting from Day 0 and refreshed daily. Mice were gavaged with 10^8^ CFU *Sb* ( *SbGal⁺* or *SbGal⁺-NLuc*) Day 0 and Day 1 for 2 days. Fecal samples were collected daily (Day 1 and Day 2). On Day 2, the mice were sacrificed on Day 2 and intestinal contents (small intestine, cecum, colon) were collected.

1−2 pieces of stool or intestinal matter were collected in preweighed 1.5 mL centrifuge tubes and then weighed again to determine fecal mass. Fecal matter was then resuspended in 1 mL PBS per 10 mg feces. Serial dilutions of fecal suspensions were plated on YPD media containing 50 μg/mL nourseothricin and 0.25 mg/mL streptomycin. Plates were sealed with parafilm and incubated at 37 °C for 2−3 days.

For Nanoluciferase activity detection, 100 μl of fecal suspension was serially diluted (1:10, 1:100 and 1:1000) in 1X PBS then mixed with 100 µl NanoGlo Assay Reagent (10 µl Nano-Glo Luciferase Assay Substrate and 90 µl sterile 1X PBS) in a 96 well black opaque plate. Suspensions were incubated for 5 minutes for luminescent reaction to take place. Luminescence was measured in a plate reader (BioTek Synergy™ H1, emission 420 nm). Luminescence values were normalized by CFU values for each replicate.

#### pGAlTet-NLuc Experiments

Six-week old female C57BL/6J mice were obtained from Jackson Laboratories and hosted at the NCSU Biological Resources Facility (BRF) for 3−4 days before experiments. Mice were housed in groups of four and their cages were changed before treatment with antibiotic cocktail, inducer administration and before treatment with *SbGal⁺* with *pGalTet-NLuc (*AND-gate) plasmid. Antibiotic cocktail (ampicillin (0.5 mg/mL), gentamicin (0.5 mg/mL), metronidazole (0.5 mg/mL), neomycin (0.5 mg/mL), vancomycin (0.25 mg/mL)), sucralose (4 mg/mL) was administered *ad libitum* in filter sterilized drinking water starting from 3 days prior to *Sb* gavage and continued throughout the experiment and refreshed daily. Galactose induction (20 mg/mL) and aTc induction (0.25 mg/mL) was administered via *ad libitum* in filter sterilized drinking water starting from Day 0 and refreshed daily. Mice were gavaged with 10^8^ CFU *Sb* on Day 0 and Day 1 for 2 days. Fecal samples were collected daily (Day 1 and Day 2). On Day 2, the mice were sacrificed on Day 2 and intestinal contents (small intestine, cecum, colon) were collected.

1−2 pieces of stool or intestinal matter were collected in preweighed 1.5 mL centrifuge tubes and then weighed again to determine fecal mass. Fecal matter was then resuspended in 1 mL PBS per 10 mg feces. Serial dilutions of fecal suspensions were plated on YPD media containing 50 μg/mL nourseothricin and 0.25 mg/mL streptomycin. Plates were sealed with parafilm and incubated at 37 °C for 2−3 days.

For Nanoluciferase activity detection, 100 μl of fecal suspension was serially diluted (1:10, 1:100 and 1:1000) in 1X PBS then mixed with 100 µl NanoGlo Assay Reagent (10 µl Nano-Glo Luciferase Assay Substrate and 90 µl sterile 1X PBS) in a 96 well black opaque plate. Suspensions were incubated for 5 minutes for luminescent reaction to take place. Luminescence was measured in a plate reader (BioTek Synergy™ H1, emission 420 nm). Luminescence values were normalized by CFU values for each replicate.

One set of lower gastrointestinal tract tissue per group was collected in a petri dish and processed for tissue imaging at BRIC Small Animal Imaging Facility core facility at UNC Chapel Hill. After collection, the tissue was transferred in ice to the facility and incubated in 10 mL NanoGlo Assay Reagent (200 µl Nano-Glo Luciferase Assay Substrate and 9.8 mL sterile 1X PBS) for 5 minutes shaking at 50 rpm, at room temperature. After incubation, the reagent was discarded and the tissue was imaged for bioluminescent photon emission using IVIS® Spectrum (PerkinElmer Inc.) with exposure times 1-5 minutes. with exposure times ranging from 1 to 5 min, depending on the signal intensity.

## Acknowledgements

We gratefully acknowledge experimental assistance from Dr. Jonathan Frank at the BRIC Small Animal Imaging Facility core facility at UNC Chapel Hill, Sandy Elliot at the NCSU Biological resources facility, and members of the Crook lab for helpful discussions.

DD and NC were supported by the National Science Foundation (CBET-1934284) and the Novo Nordisk Foundation (NNF19SA0035474). I.S.A. is supported by NCSU CBE startup funds and the Ministry of Higher Education - Oman. DJH was supported by the NC State Park Scholarship. AC and ASM were supported by the National Science Foundation (MCB-1947498). CS, RVU, and MOAS were supported by the Novo Nordisk Foundation (NNF20CC0035580, NNF17CO0028232, and NNF19SA0035438).

## Author Contributions

DD, DJH, ISA, and NC designed and conceived the study. DD, DJH, ISA, CS and KD performed all yeast engineering and cultivation experiments. DD led all mice experiments with help from ISA. AC and DD performed and analyzed microscopy imaging for yeast display. NC supervised the research. DD, DJH, ISA and NC wrote the manuscript. NC, ASM, RVU, and MOAS solicited funding. All authors read and approved the final manuscript.

