## Supplementary figures and images for "Programming Probiotics: Diet-responsive gene expression and colonization control in engineered *S. boulardii*"

### DD576-pGAL1-AGA1-tENO1-INT-1-URA-AmpR-ColE1-sequence.pdf

#### DD576 pGAL1, AGA1, tENO1, INT 1, URA, AmpR, Col...

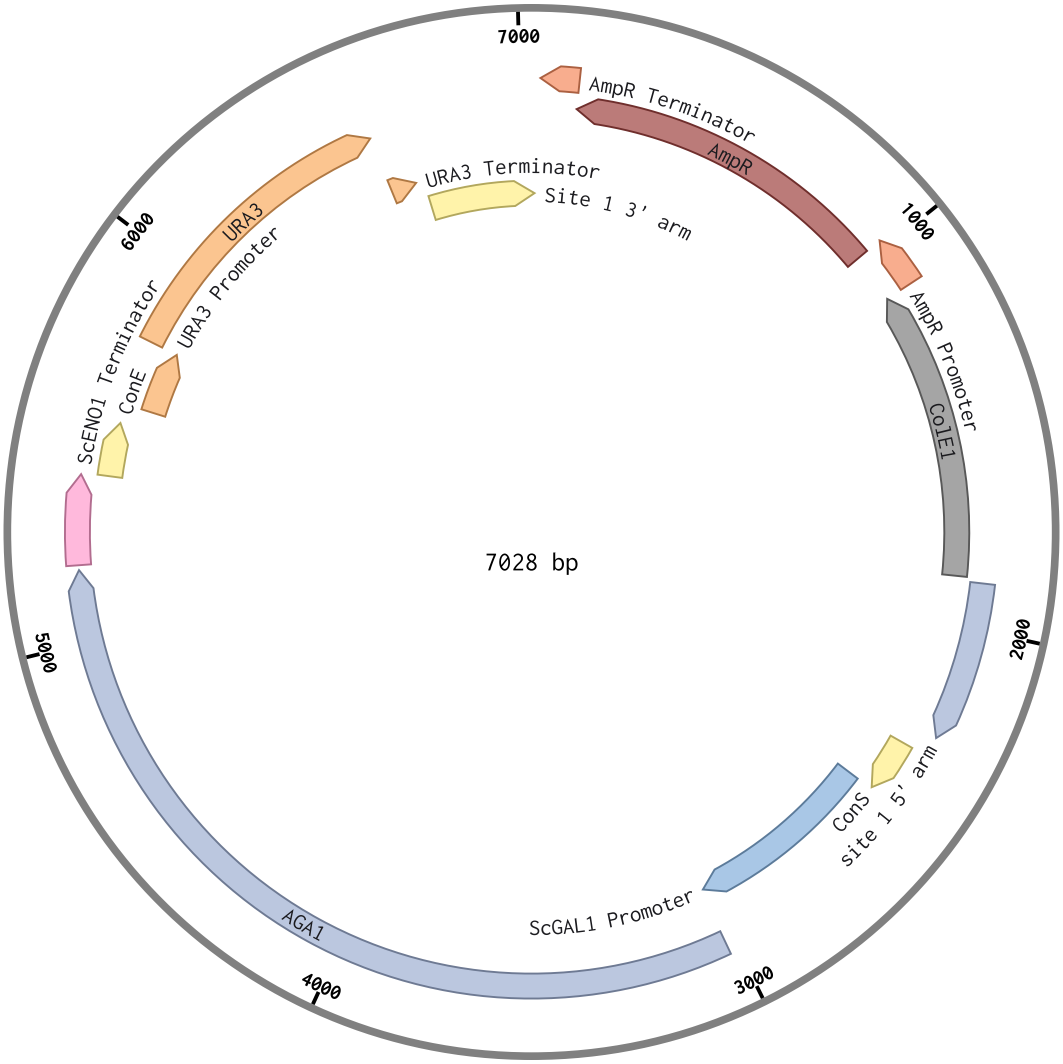

### DD580-pGAL-AGA2-tMATa-HIS-CEN-AmpR-ColE1-sequence.pdf

#### DD580 pGAL, AGA2, tMATa, HIS, CEN, AmpR, ColE1 ...

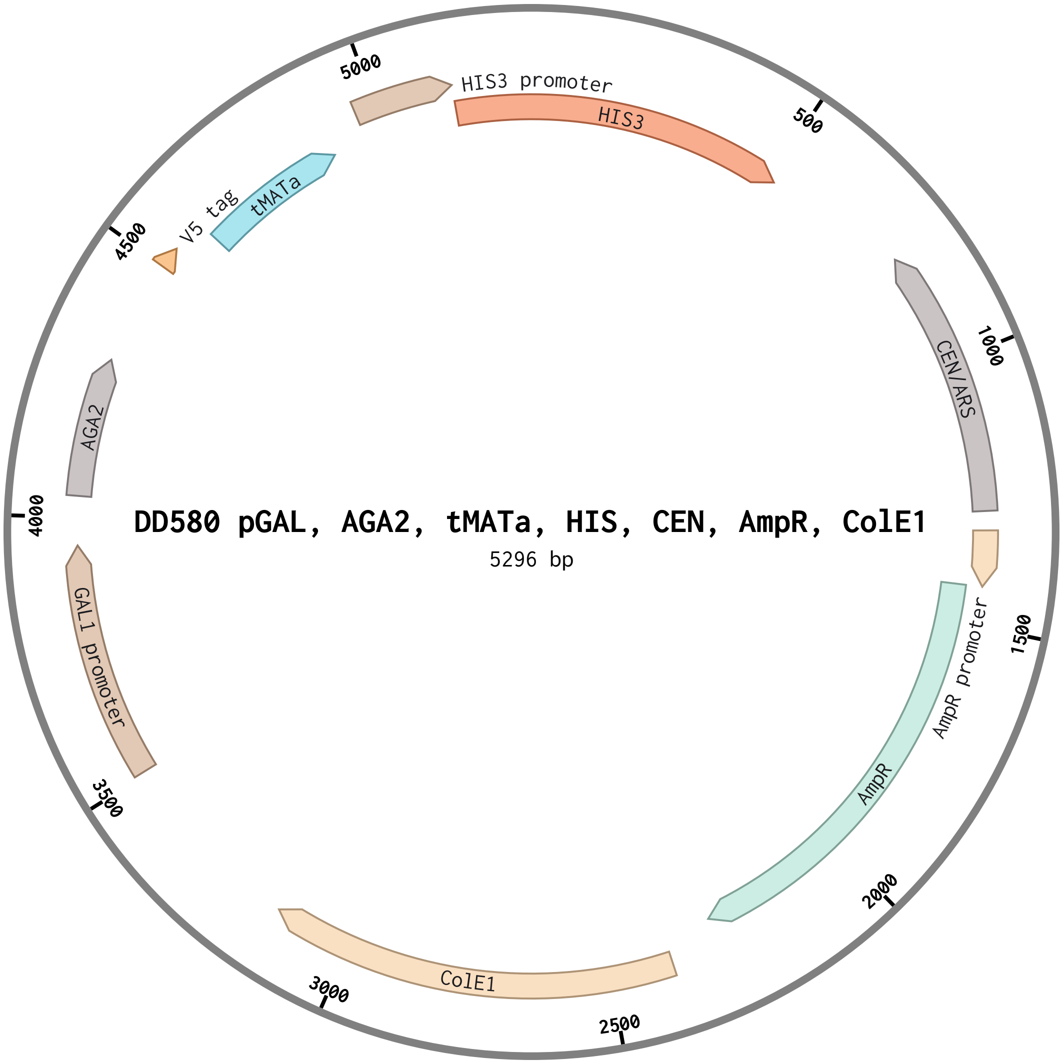

### DD592-pGAL-mKate2-tENO1-pTET-yeGFP-tENO1-URA-2micron-AmpR-sequence.pdf

#### DD592 pGAL, mKate2, tENO1, pTET, yeGFP, tENO1, ...

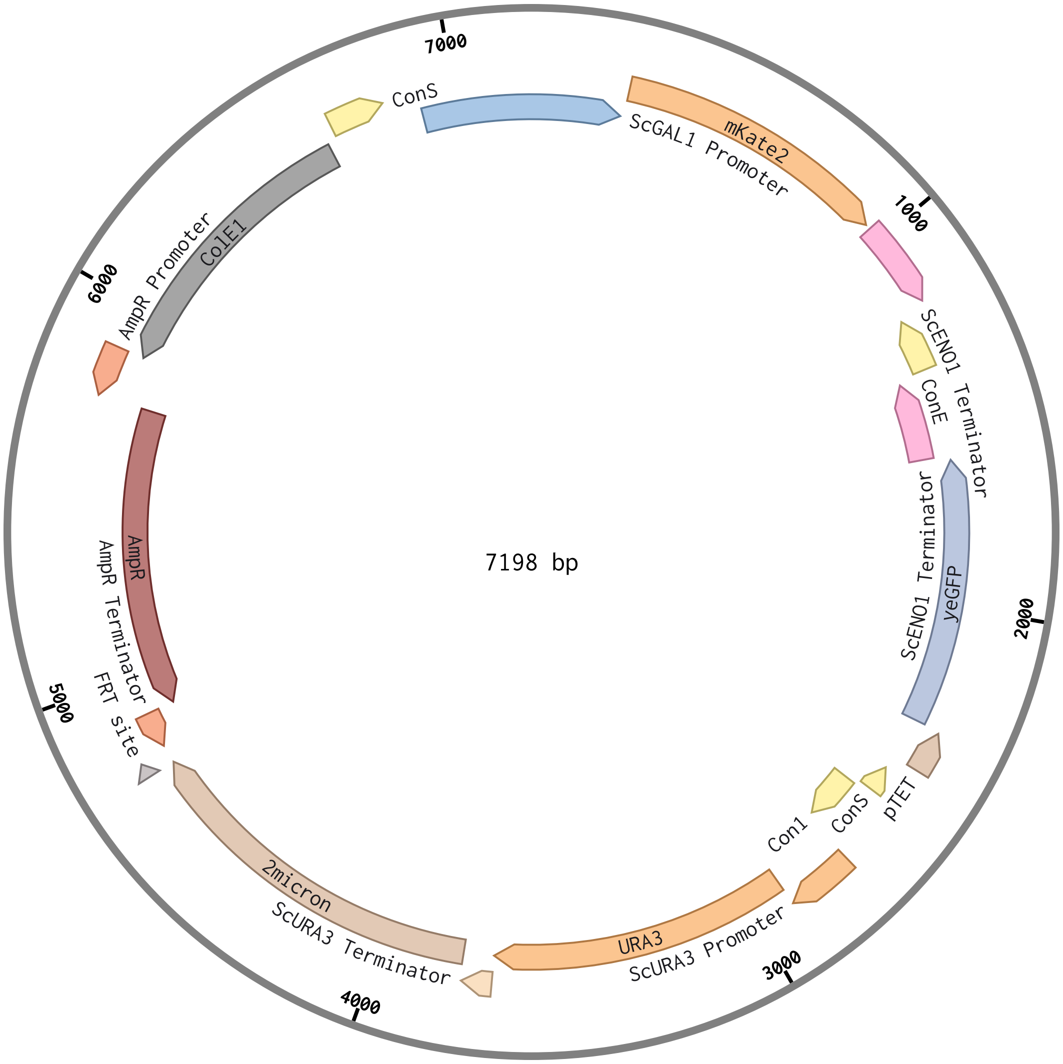

### DD593-pGalTet-NanoLuc-tSSA1-URA-2micron-AmpR-sequence.pdf

#### DD593 pGalTet, NanoLuc, tSSA1, URA, 2micron, Am...

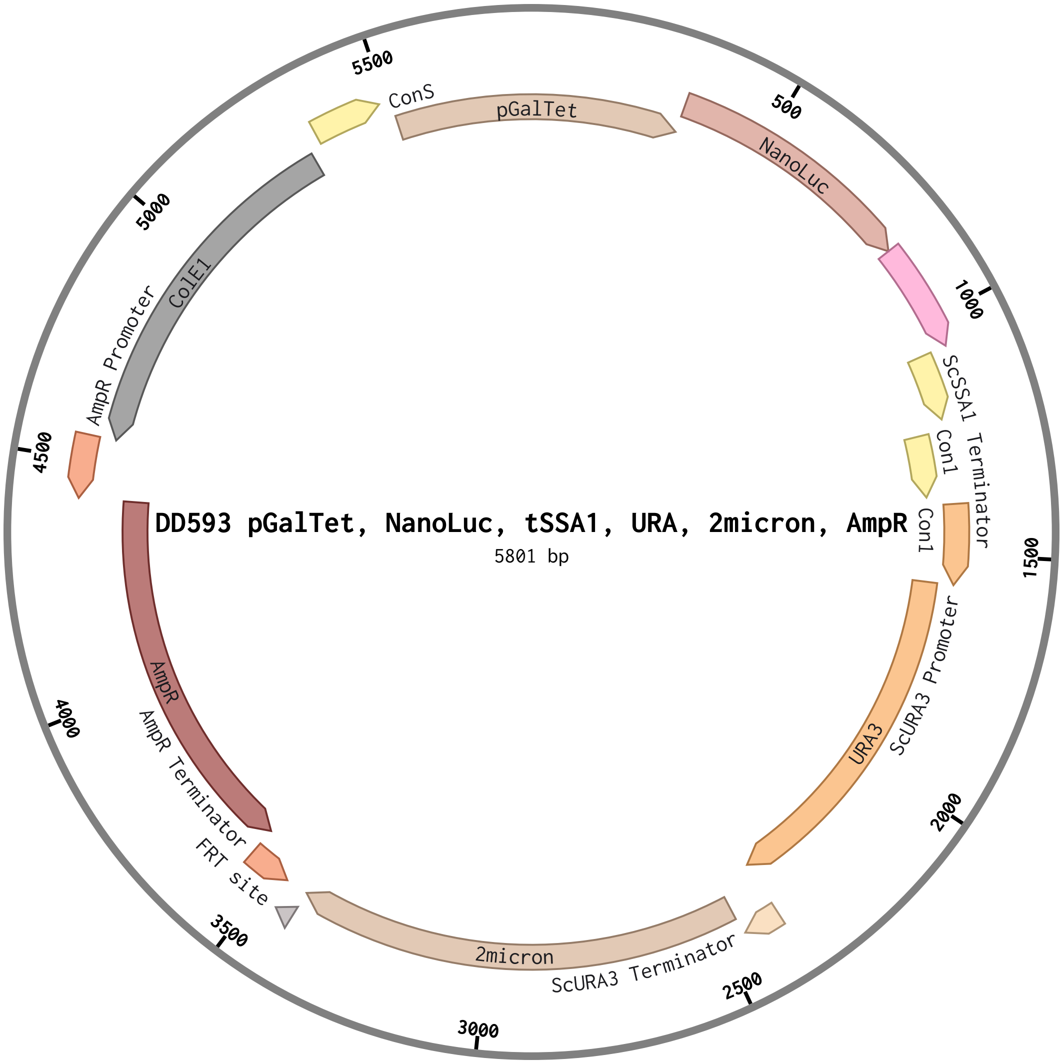

### DD594-pGalTet-yeGFP-tENO1-URA-2micron-AmpR-ColE1-sequence.pdf

#### DD594 pGalTet, yeGFP, tENO1, URA, 2micron, AmpR...

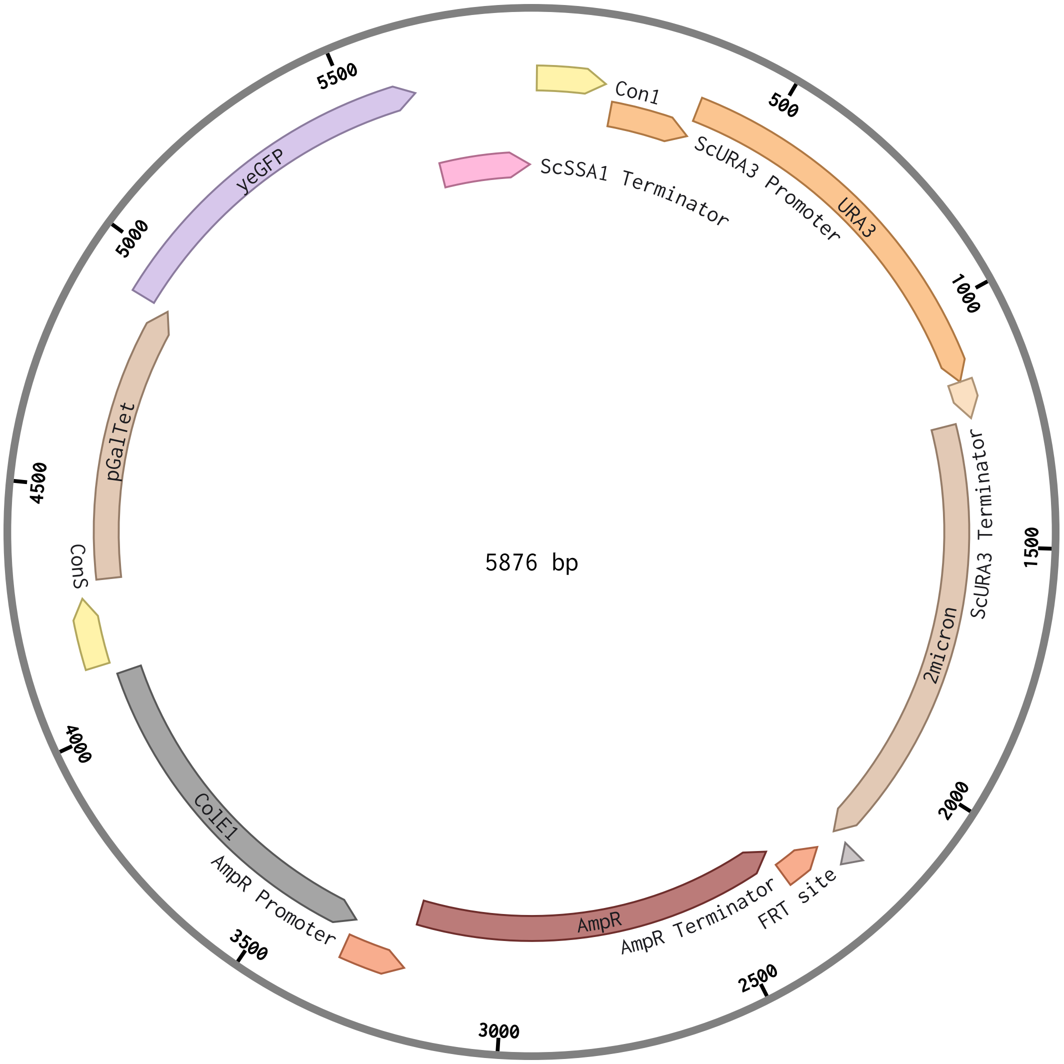

### DD608-pGal-AGA2-SA1-tMATa-HIS-CEN-AmpR-ColE1-sequence.pdf

#### DD608 pGal, AGA2, SA1, tMATa, HIS, CEN, AmpR, C...

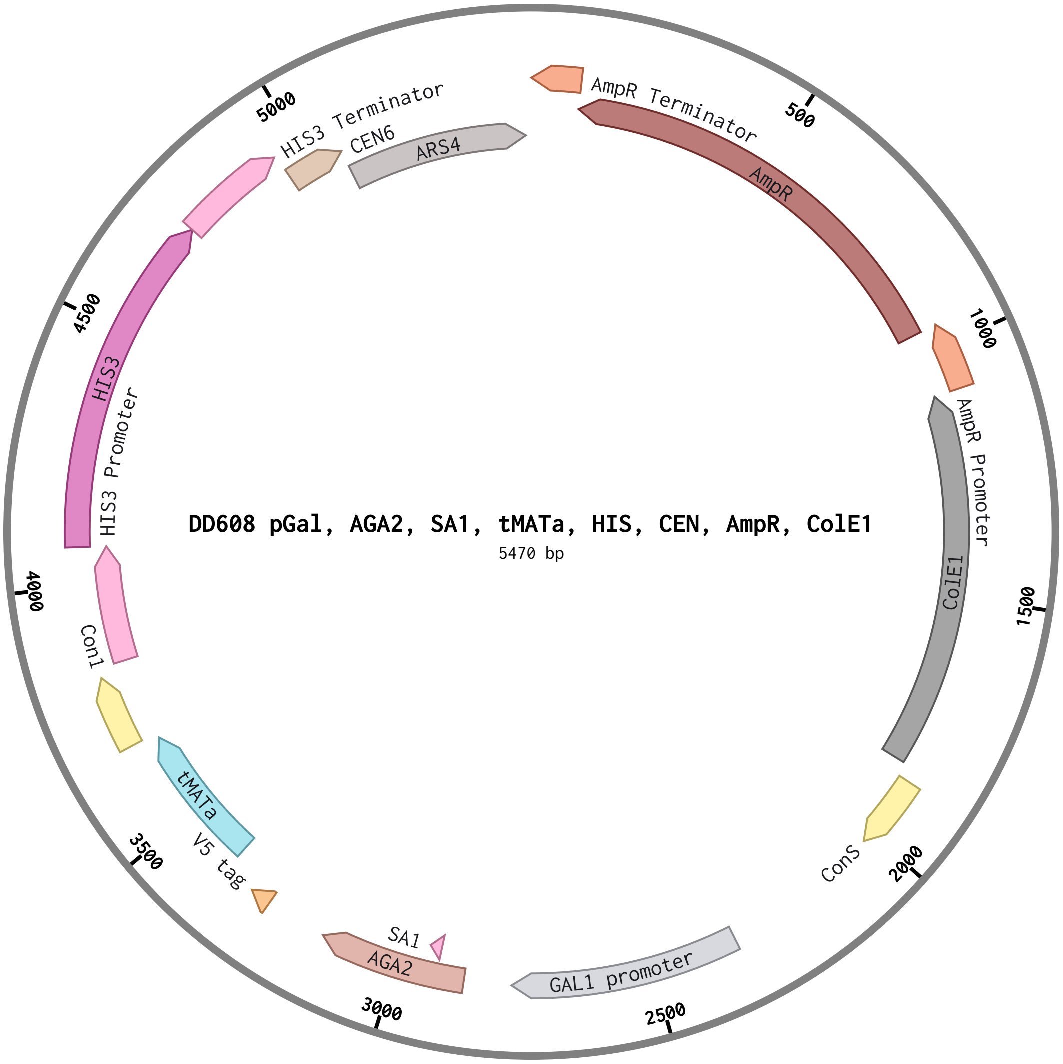

### DJH088-pXYL-yeGFP-tENO1-URA-2micron-Amp-sequence.pdf

#### DJH088 pXYL, yeGFP, tENO1, URA, 2micron, Amp (5...

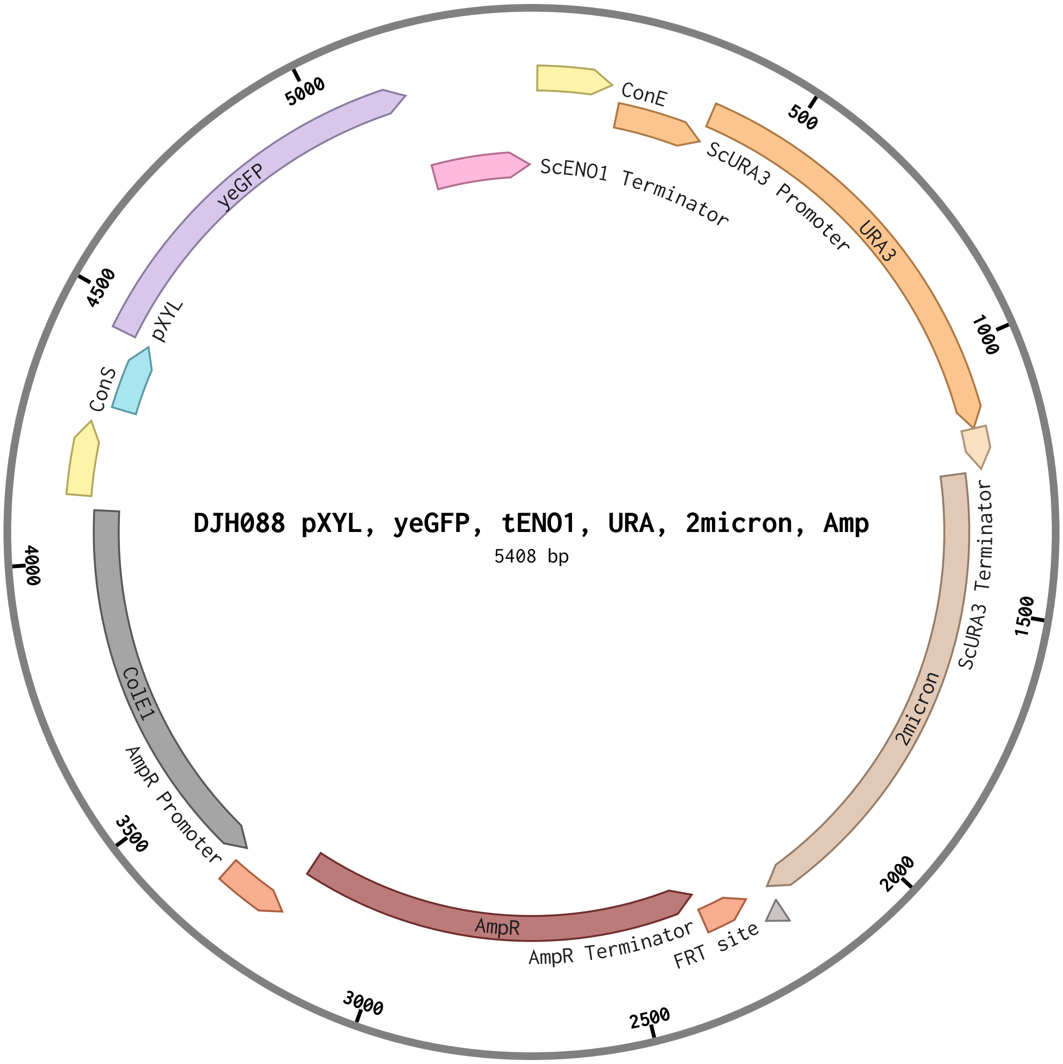

### DJH089-pLac-yeGFP-tENO1-URA-2micron-Amp-sequence.pdf

#### DJH089 pLac, yeGFP, tENO1, URA, 2micron, Amp (5...

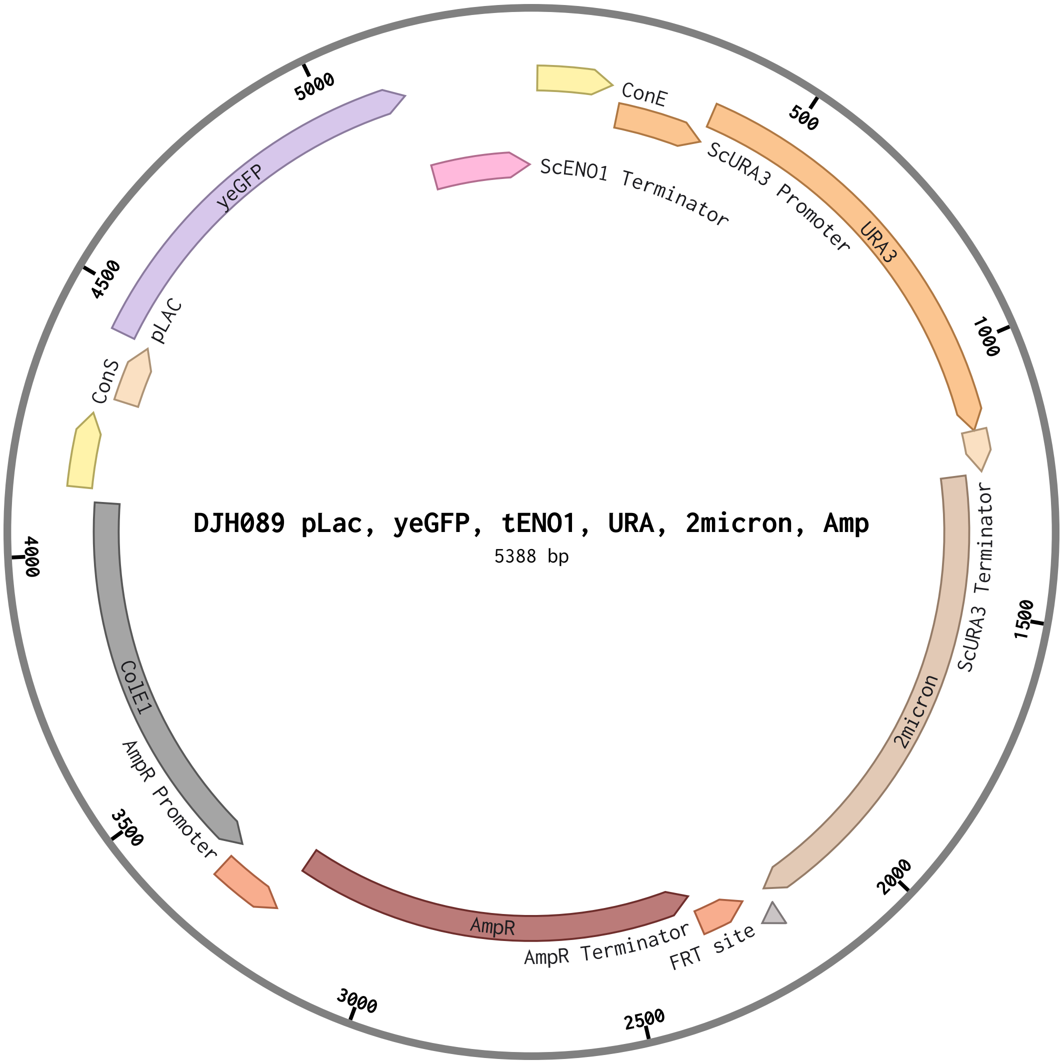

### DJH090-pTet-yeGFP-tENO1-URA-2micron-Amp-sequence.pdf

#### DJH090 pTet, yeGFP, tENO1, URA, 2micron, Amp (5...

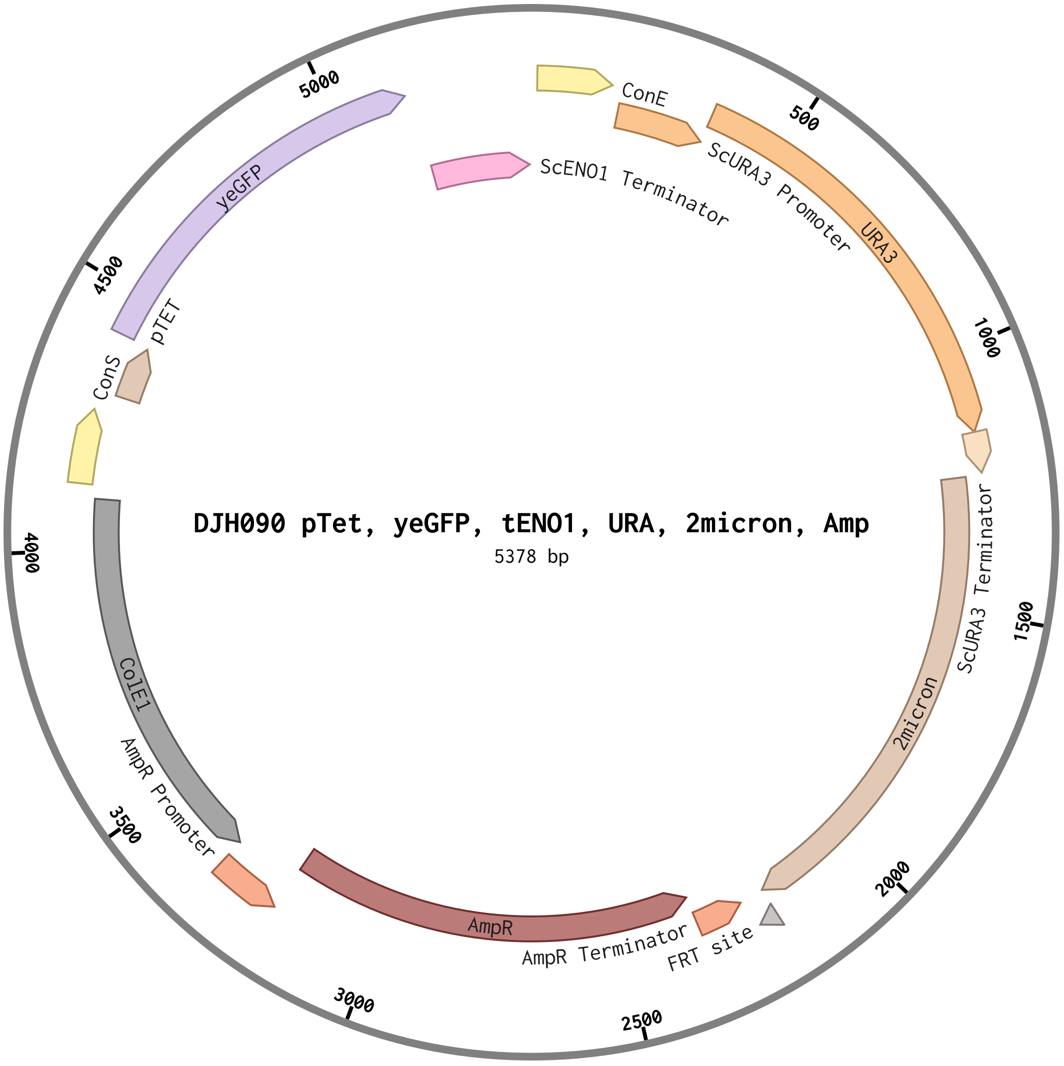

### DJH091-pXYL-CmR-ColE1-sequence.pdf

#### DJH091 pXYL, CmR, ColE1 (1811 bp)

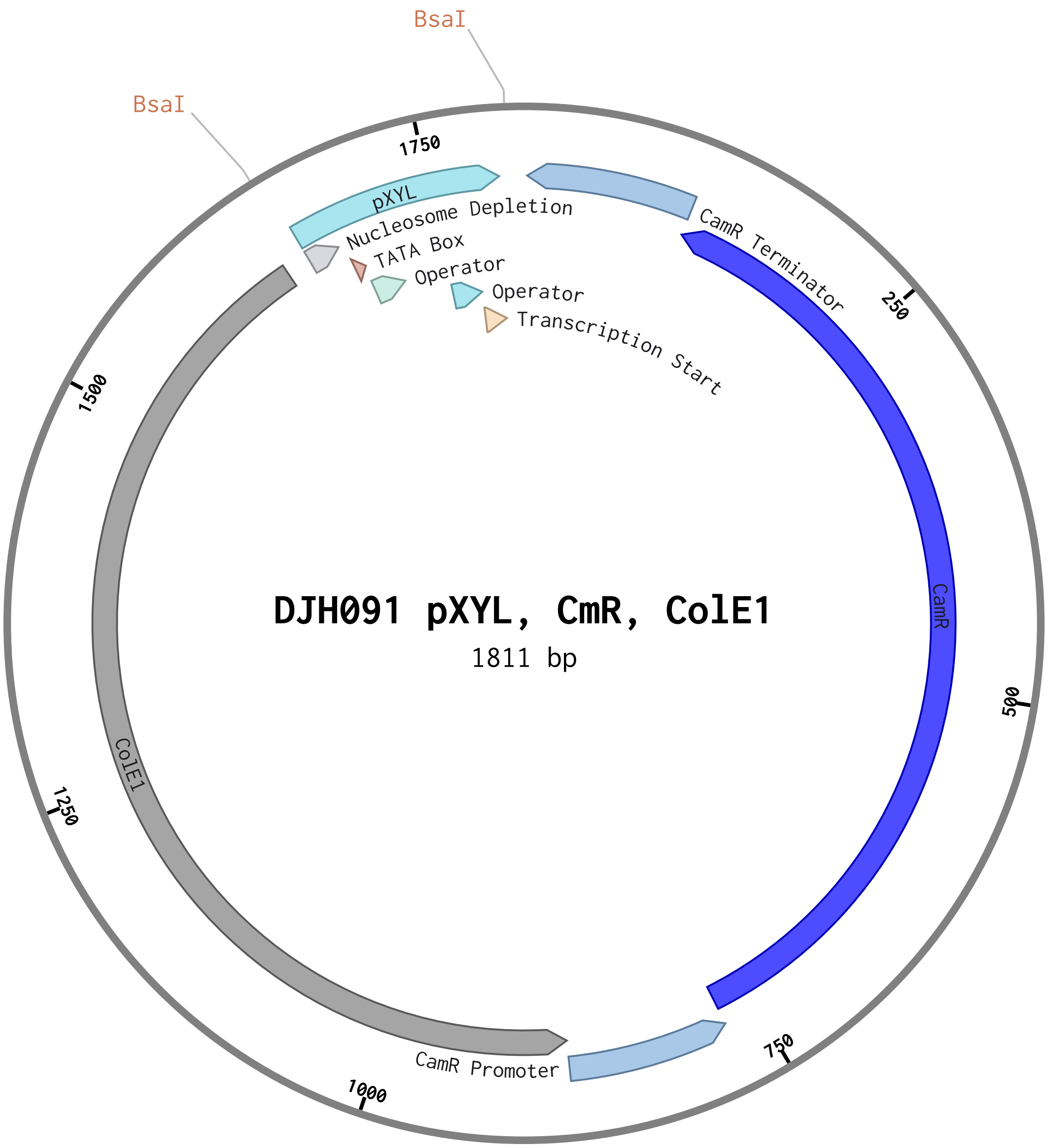

### DJH092-pLAC-CmR-ColE1-sequence.pdf

#### DJH092 pLAC, CmR, ColE1 (1791 bp)

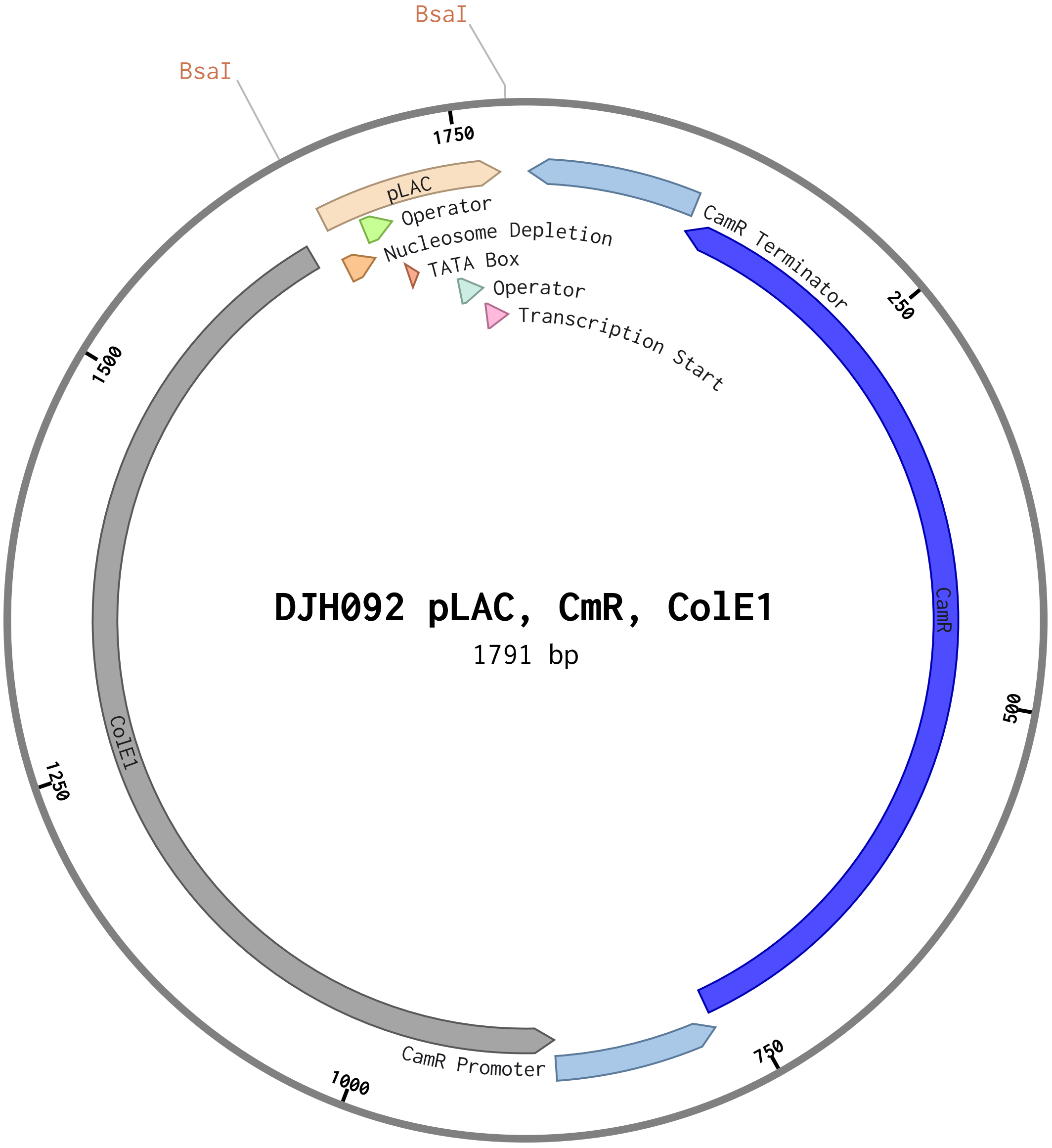

### DJH093-pTET-CmR-ColE1-sequence.pdf

#### DJH093 pTET, CmR, ColE1 (1781 bp)

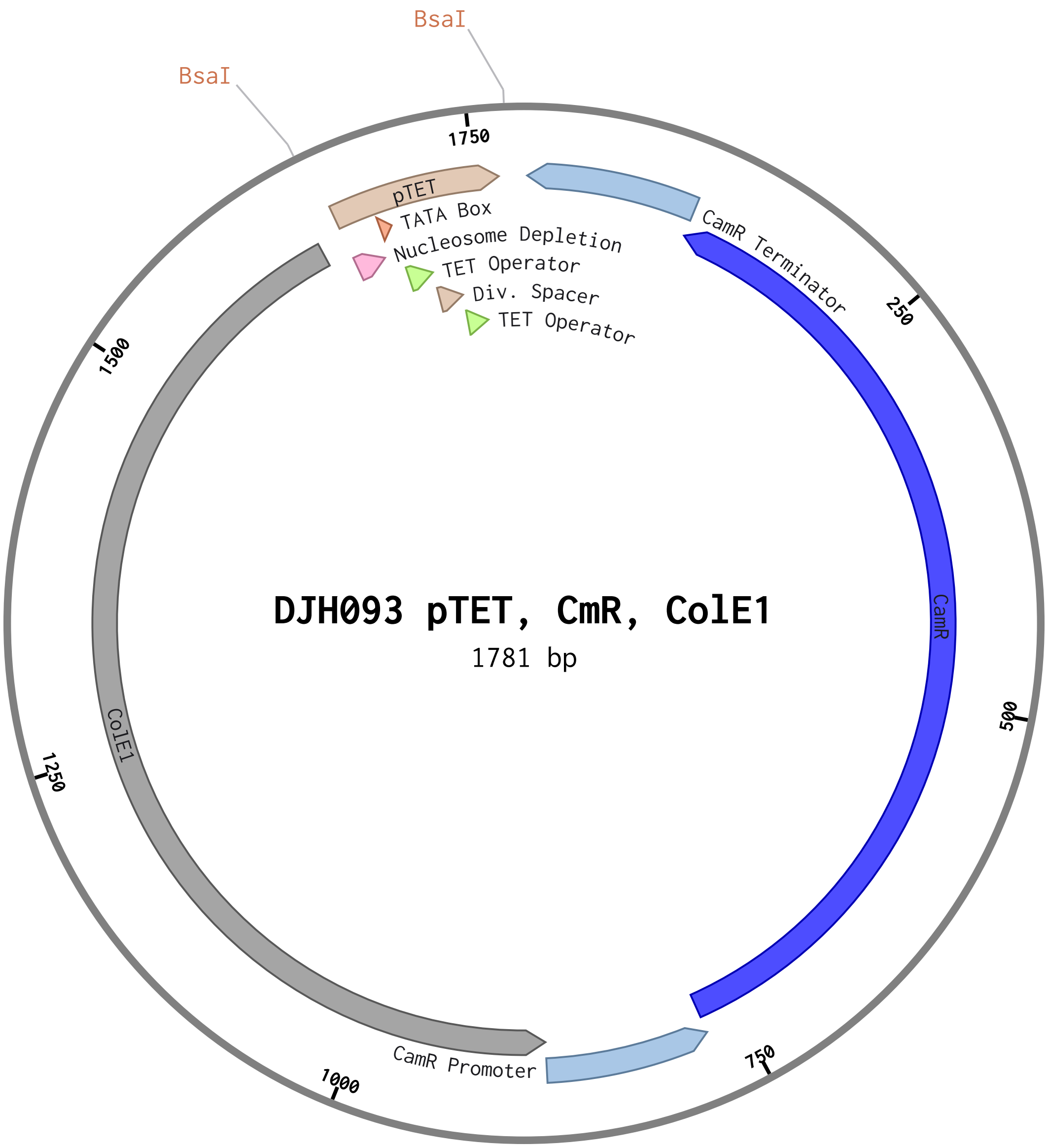

### DJH094-pFBAI-xylR_NLS-tENO2-HIS-CEN-Amp-sequence.pdf

#### DJH094 pFBAI, xylR_NLS, tENO2, HIS, CEN, Amp (5...

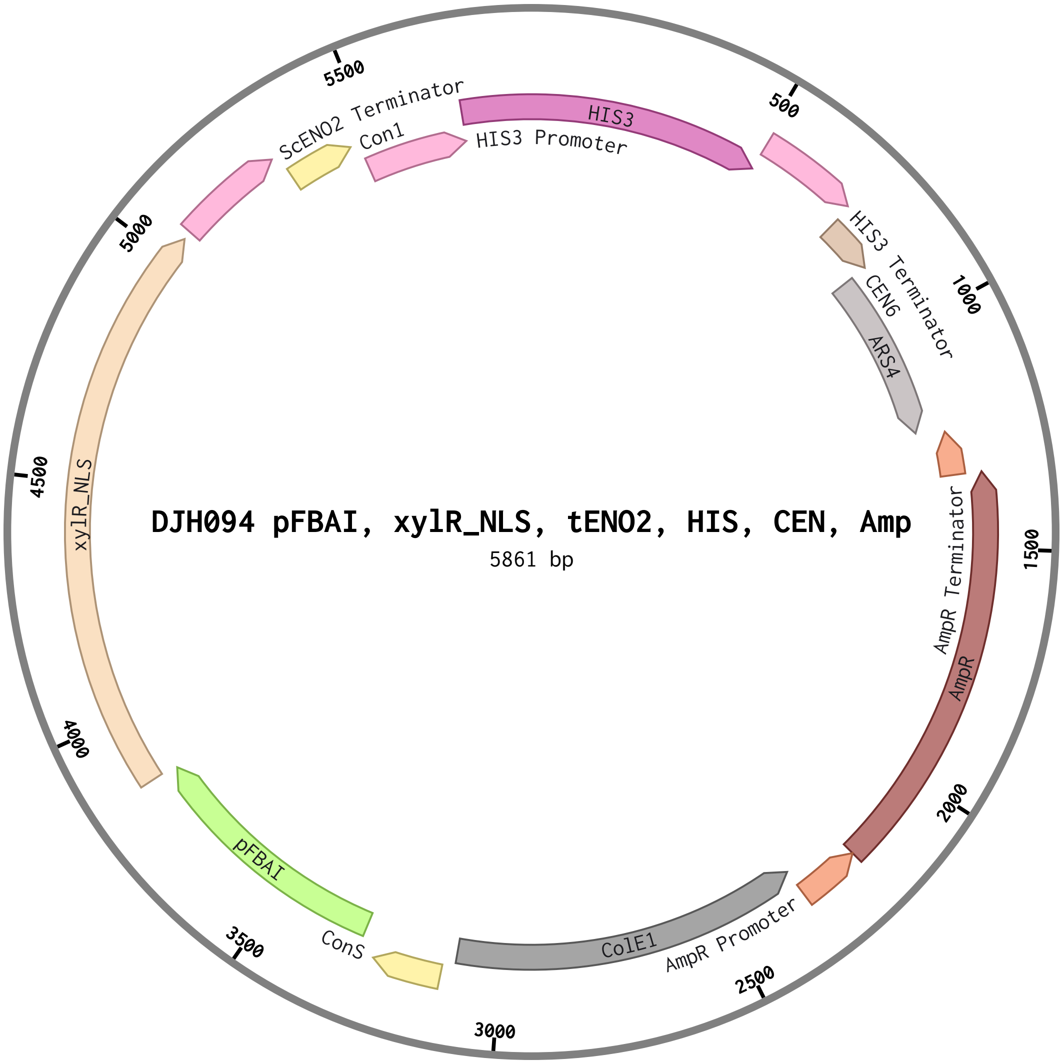

### DJH095-pFBAI-lacI_NLS-tENO2-HIS-CEN-Amp-sequence.pdf

#### DJH095 pFBAI, lacI_NLS, tENO2, HIS, CEN, Amp (5...

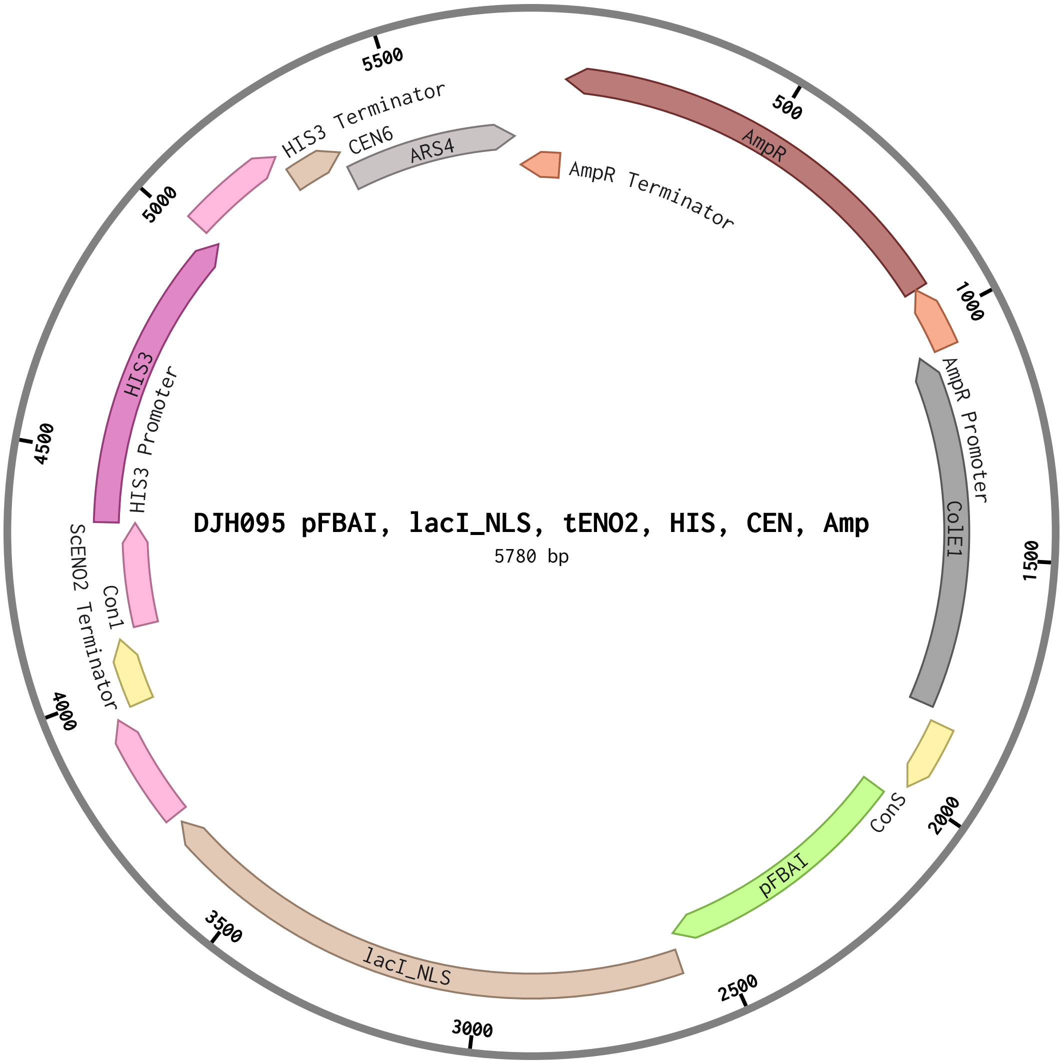

### DJH096-pFBAI-tetR_NLS-tENO2-HIS-CEN-Amp-sequence.pdf

#### DJH096 pFBAI, tetR_NLS, tENO2, HIS, CEN, Amp (5...

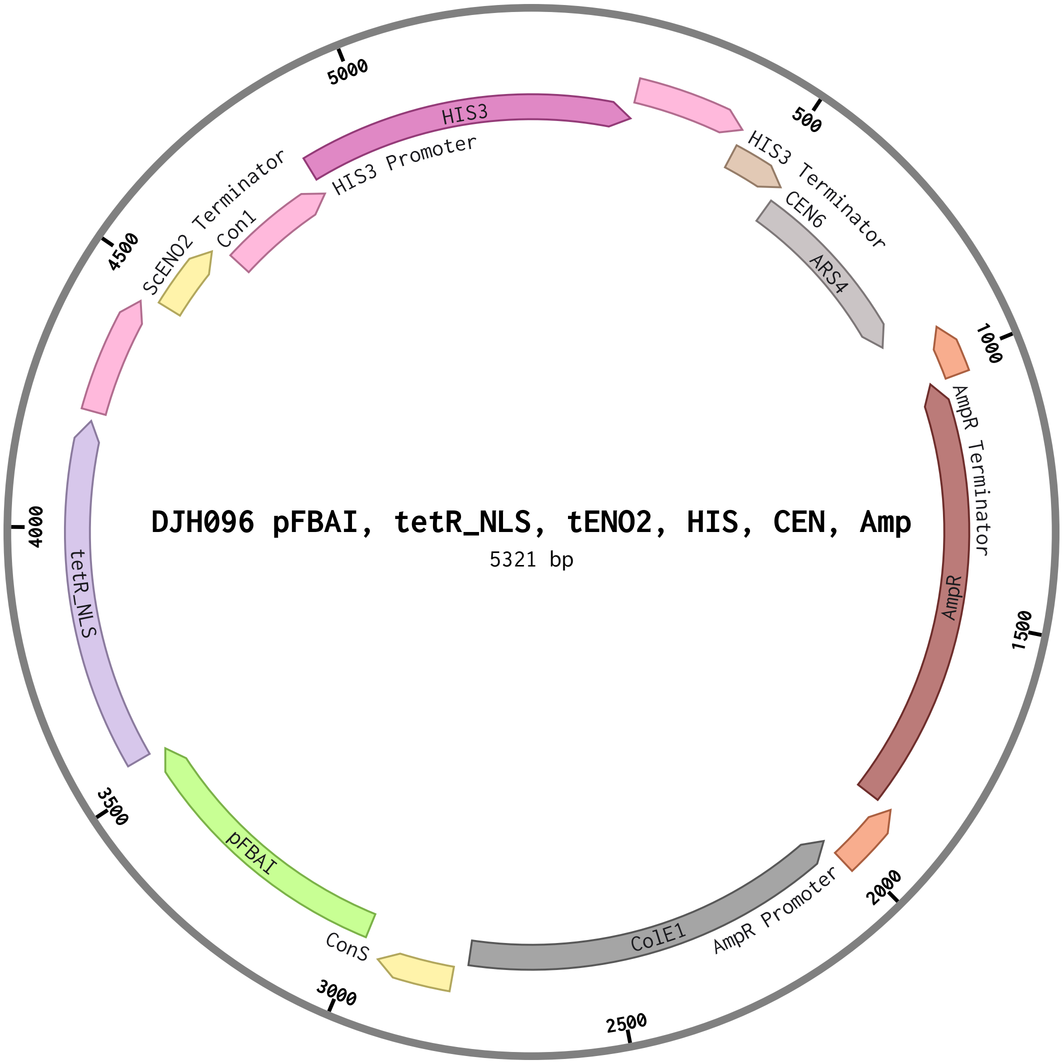

### DJH097-pCUP1-yeGFP-tENO1-URA-2micron-Amp-sequence.pdf

#### DJH097 pCUP1, yeGFP, tENO1, URA, 2micron, Amp (...

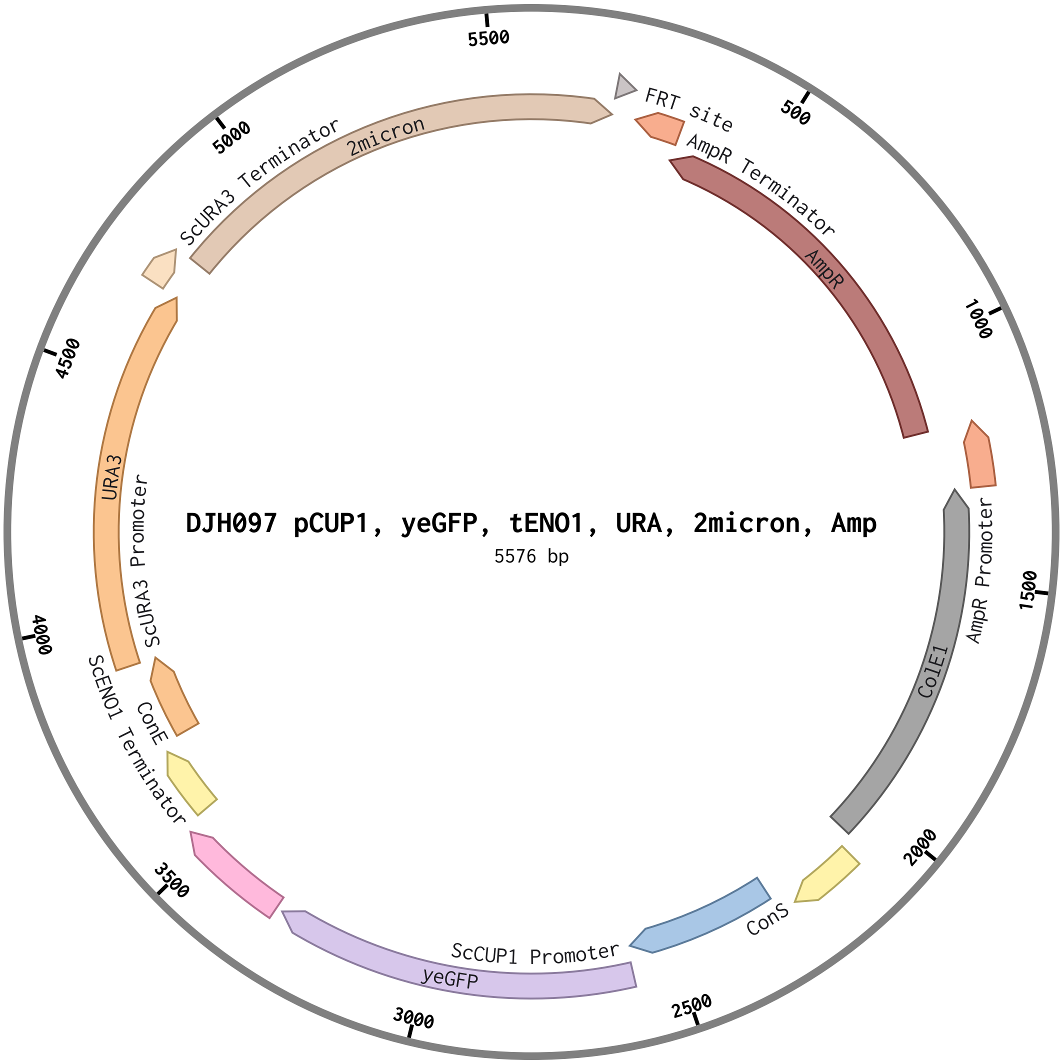

### DJH098-pGAL-yeGFP-tENO1-URA-2micron-Amp-sequence (5).pdf

#### DJH098 pGAL, yeGFP, tENO1, URA, 2micron, Amp (5...

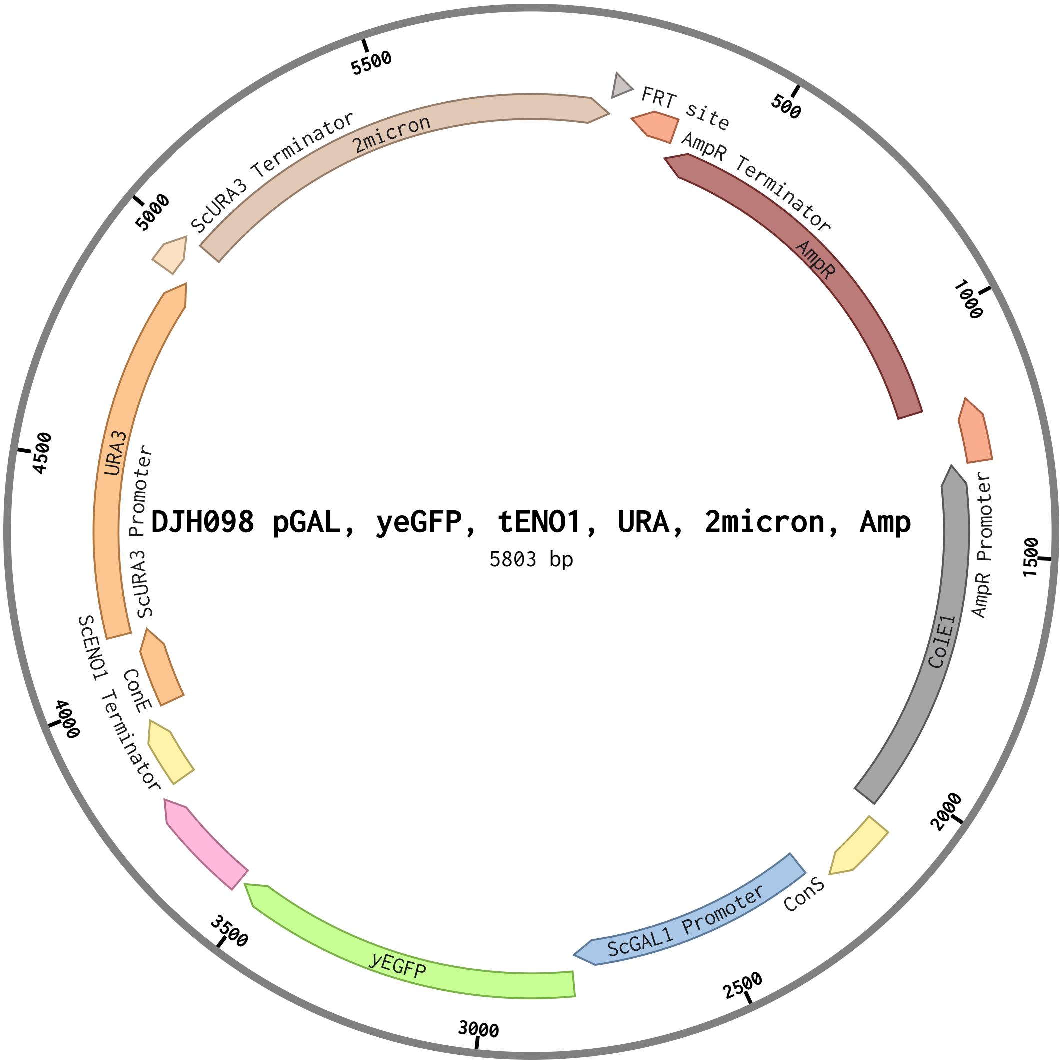

### DJH099-pGAL-CaFbFP-tENO1-URA-2micron-AmpR-sequence.pdf

#### DJH099 pGAL, CaFbFP, tENO1, URA, 2micron, AmpR ...

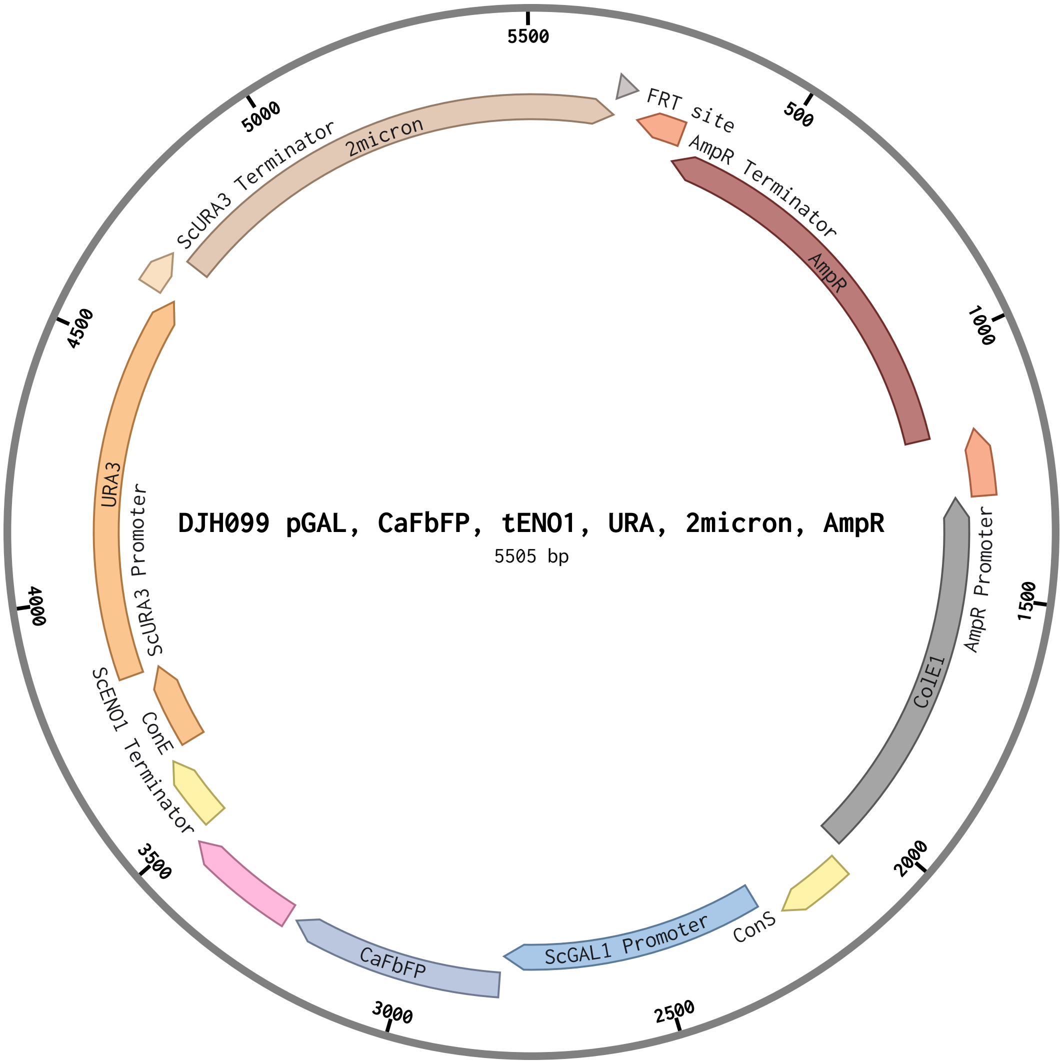

### DJH100-pCUP1-CaFbFP-tENO1-URA-2micron-AmpR-sequence.pdf

#### DJH100 pCUP1, CaFbFP, tENO1, URA, 2micron, AmpR...

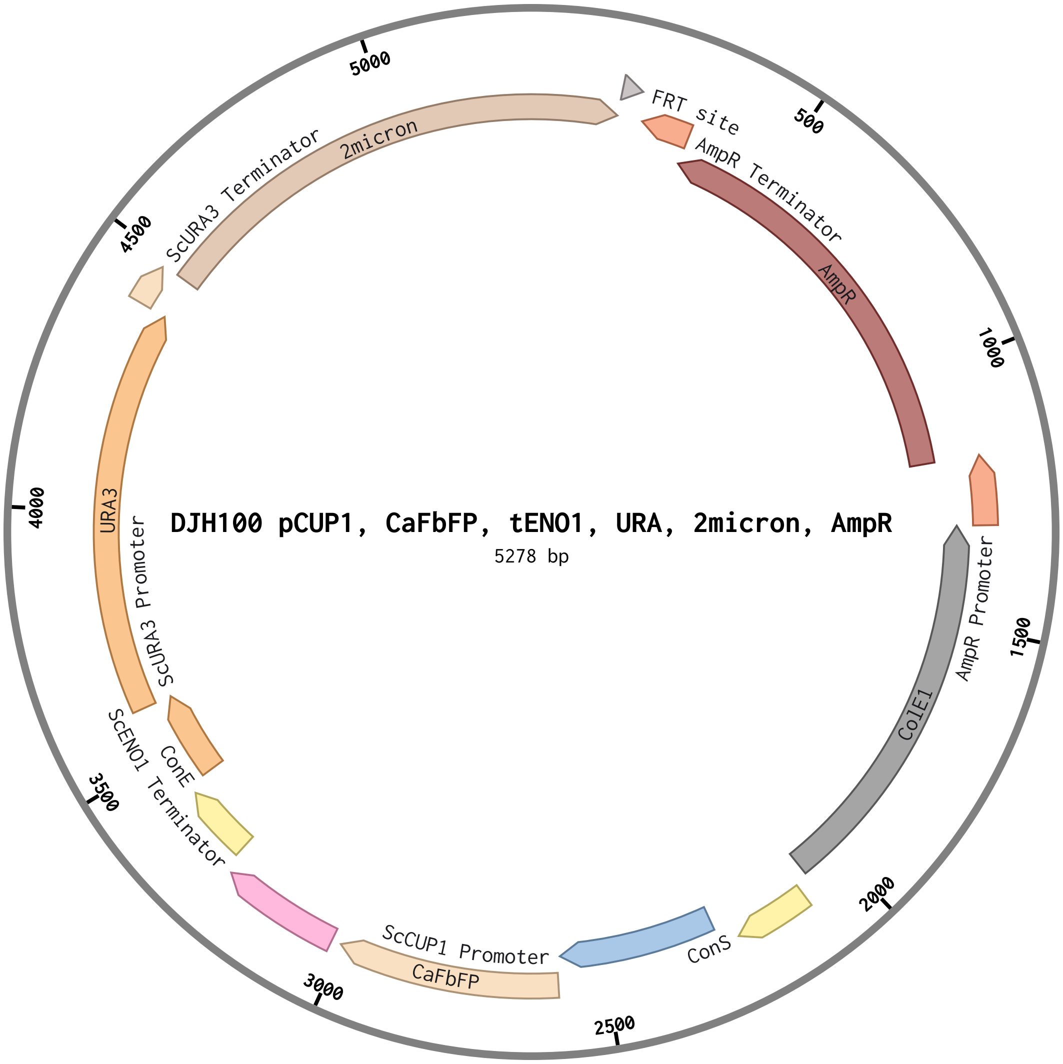

### DJH101-pXYL-CaFbFP-tENO1-URA-2micron-AmpR-sequence.pdf

#### DJH101 pXYL, CaFbFP, tENO1, URA, 2micron, AmpR ...

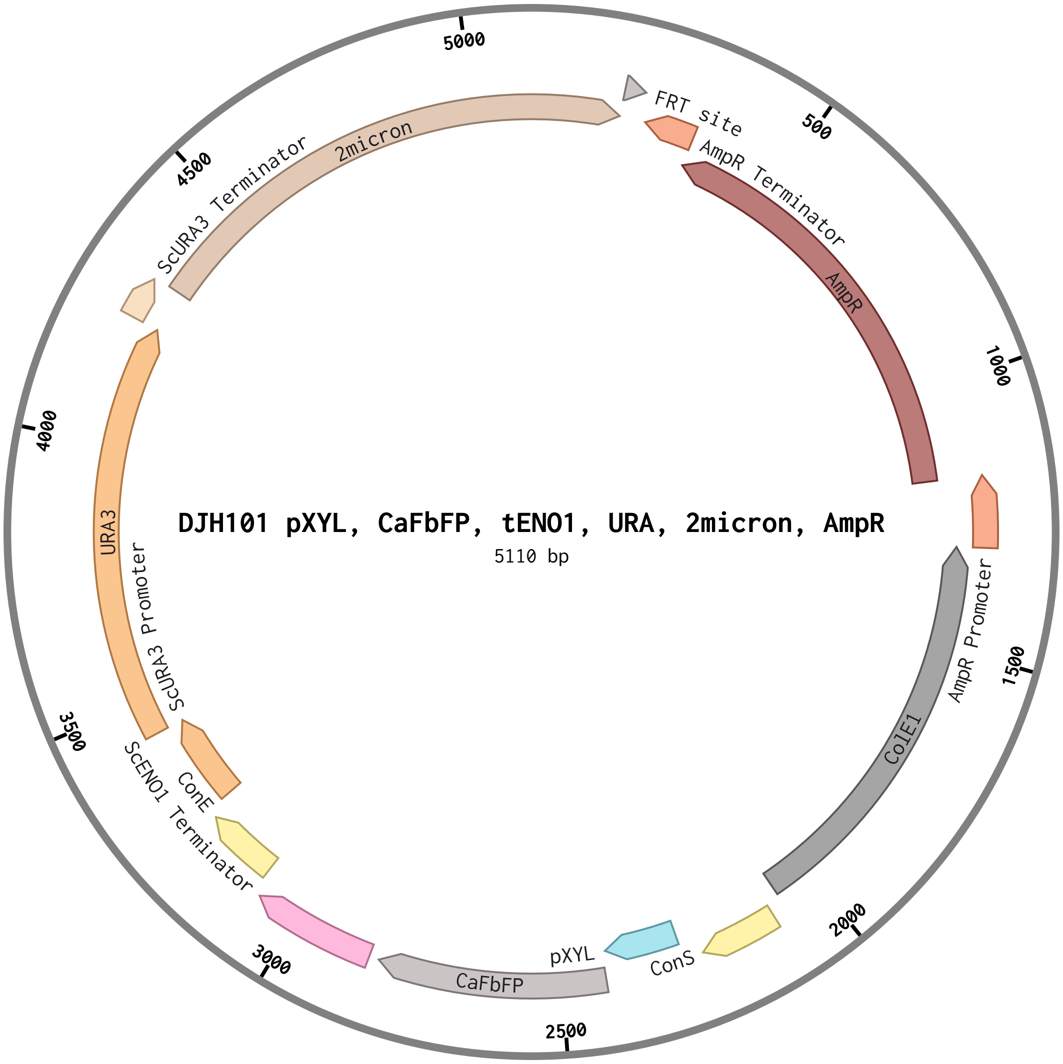

### DJH102-pLAC-CaFbFP-tENO1-URA-2micron-AmpR-sequence.pdf

#### DJH102, pLAC, CaFbFP, tENO1, URA, 2micron, AmpR...

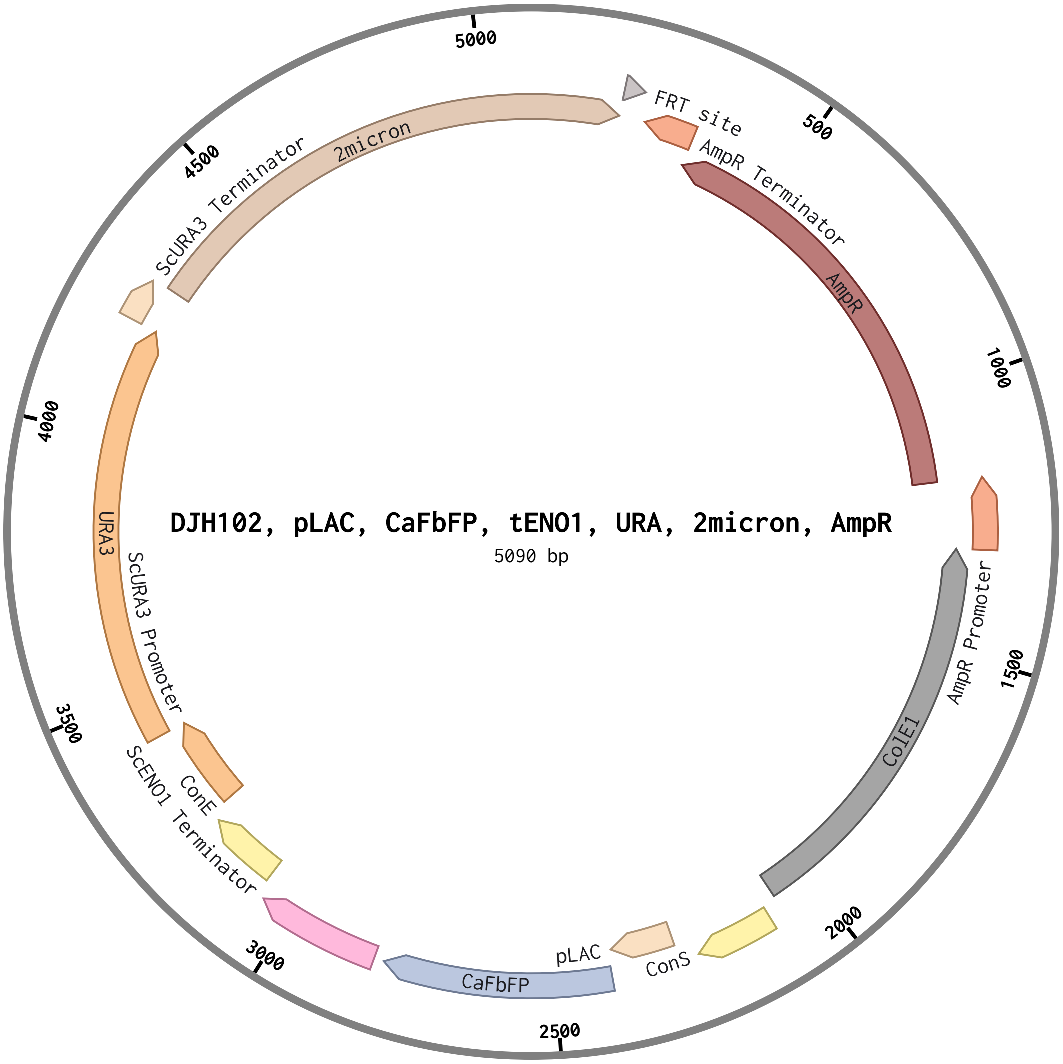

### DJH103-pTET-CaFbFP-tENO1-URA-2micron-AmpR-sequence.pdf

#### DJH103 pTET CaFbFP, tENO1, URA, 2micron, AmpR (...

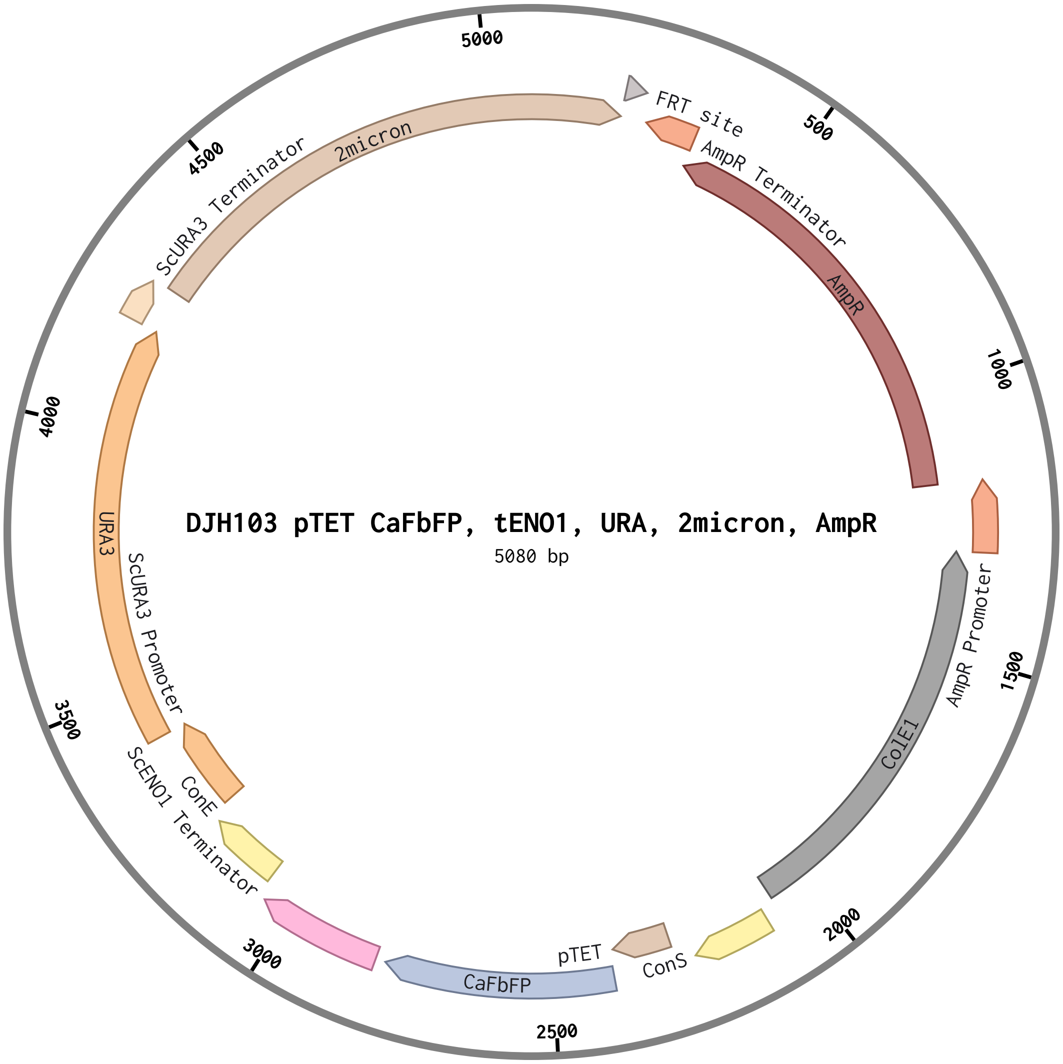

### gb_CaFbFP-sequence.pdf

#### gb_CaFbFP (460 bp)

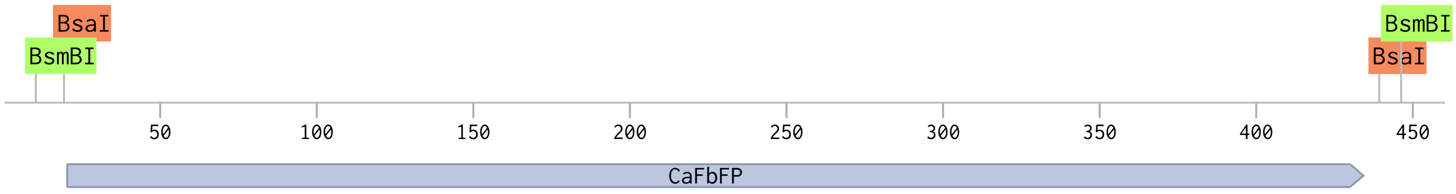

### gb_FBAI-sequence.pdf

#### gb_FBAI (599 bp)

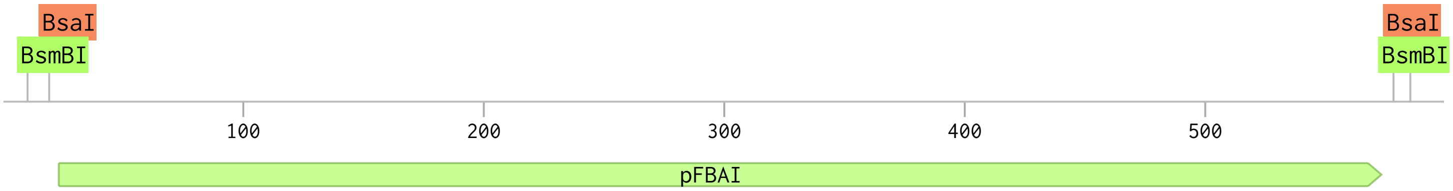

### gb_lacI_NLS-sequence.pdf

#### gb_lacI_NLS (1160 bp)

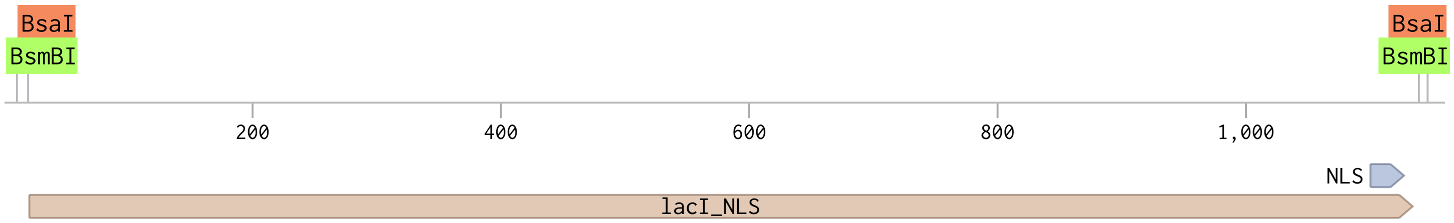

### gb_mKate2-sequence.pdf

#### gb_mKate2 (709 bp)

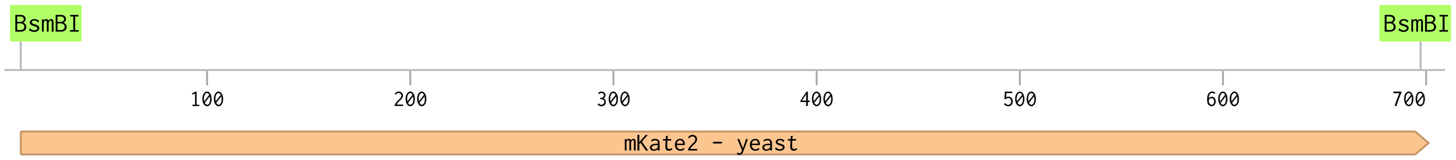

### gb_NanoLuc-sequence.pdf

#### gb_NanoLuc (562 bp)

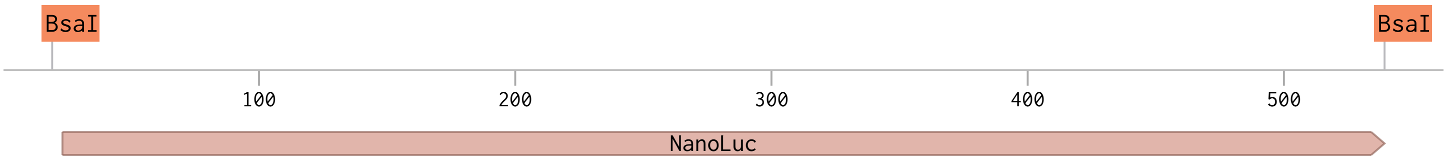

### gb_pGalTet-sequence.pdf

#### gb_pGalTet (637 bp)

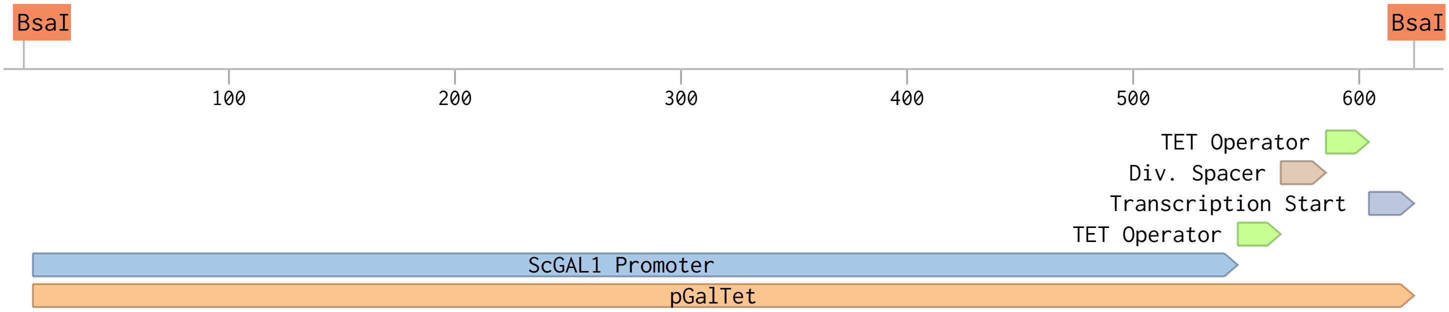

### gb_tetR_NLS-sequence.pdf

11/13/2023 10:50:28 AM

#### gb_tetR_NLS (701 bp)

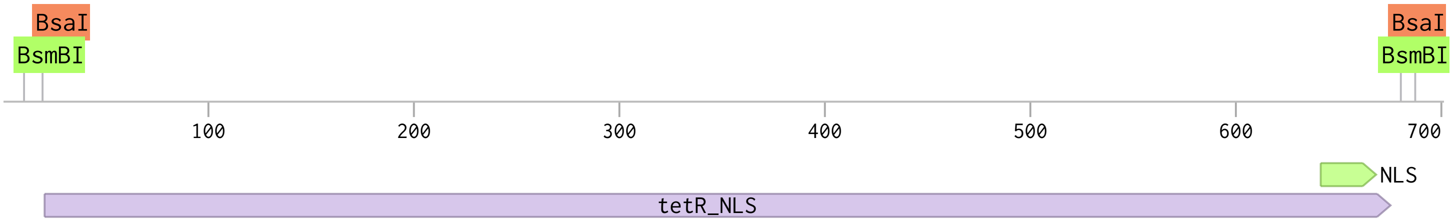

### gb_xylR_NLS-sequence.pdf

11/13/2023 10:58:12 AM

#### gb_xylR_NLS (1241 bp)

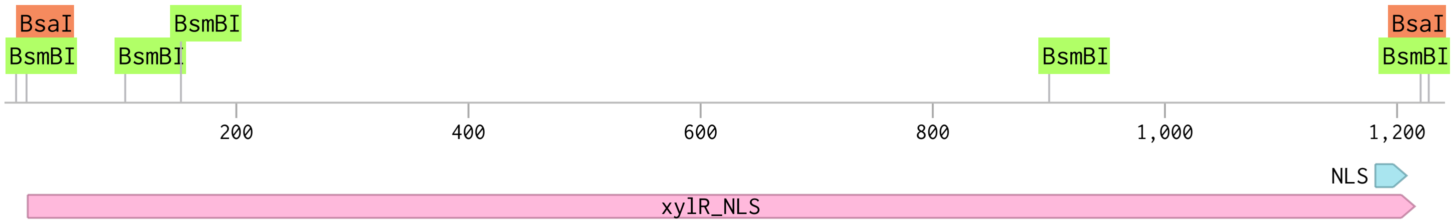

### ISA086-234-GFP-dropout-INT-1-URA-AmpR-ColE1-sequence.pdf

#### ISA086 234 GFP dropout, INT 1, URA, AmpR, ColE1...

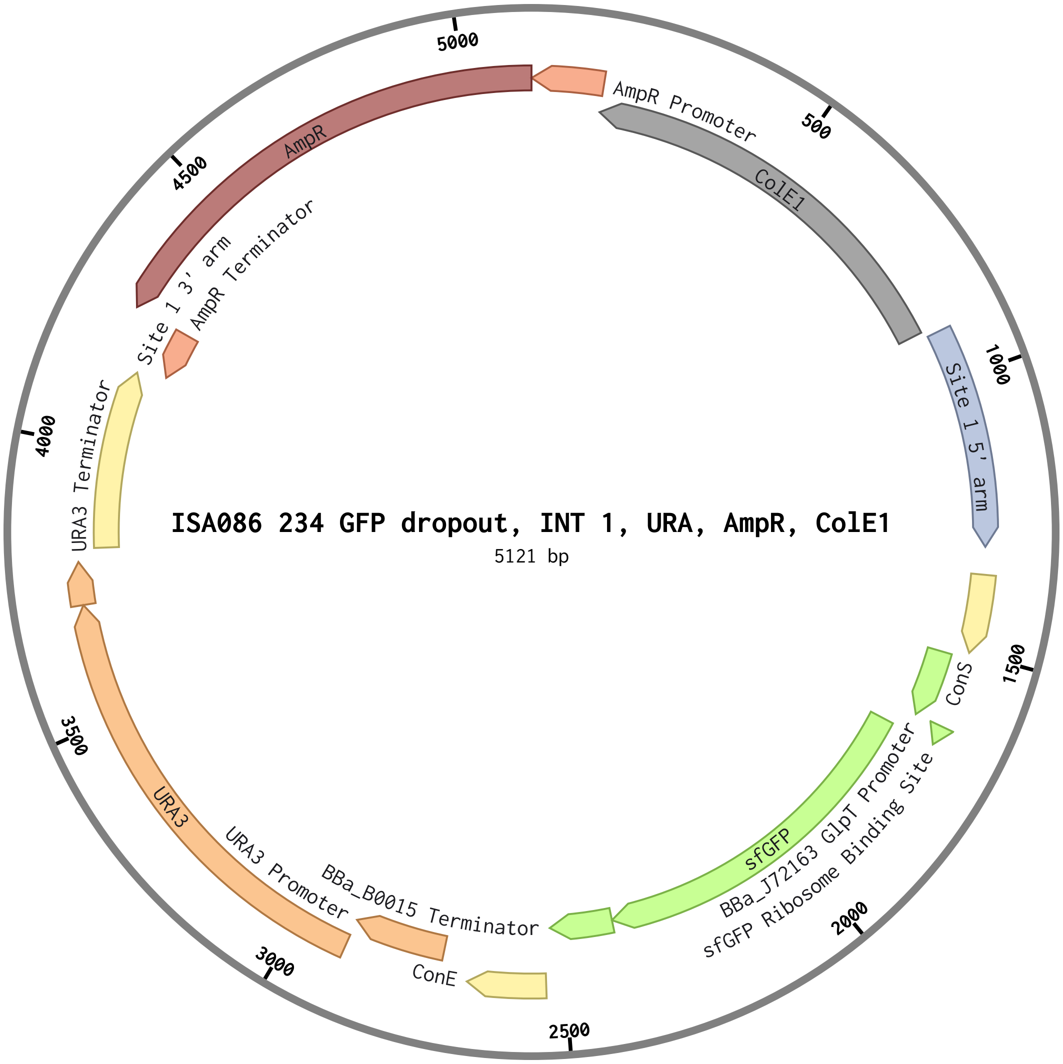

### ISA186-NatR-INT-1-site-AmpR-ColE1-sequence.pdf

#### ISA186 NatR, INT 1 site, AmpR, ColE1 (3992 bp)

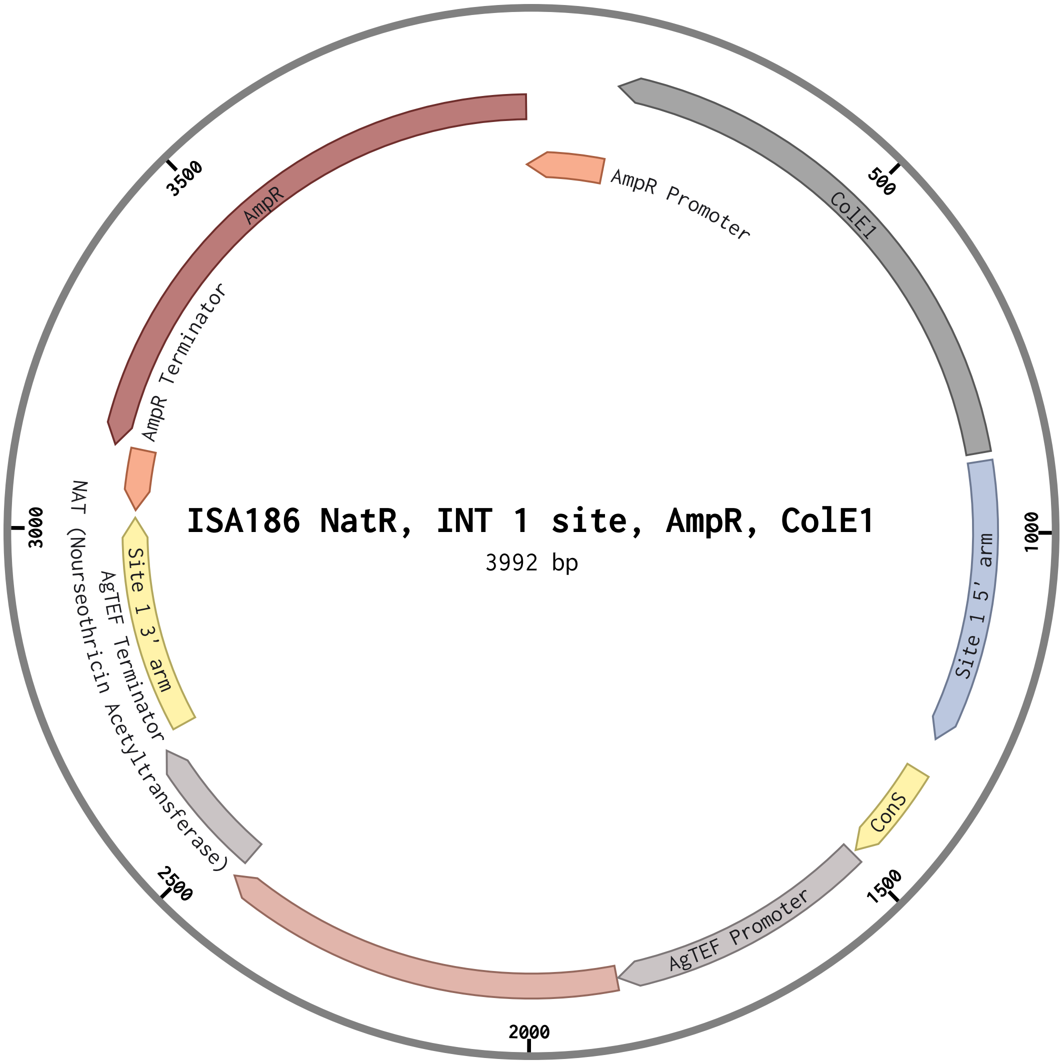

### ISA1041-pCCW12-Cas9-tSNR52-sgRNA-for-PGM2-fix-NatR-CEN-KanR-ColE1-sequence.pdf

#### ISA1041 pCCW12, Cas9, tSNR52, sgRNA for PGM2 f...

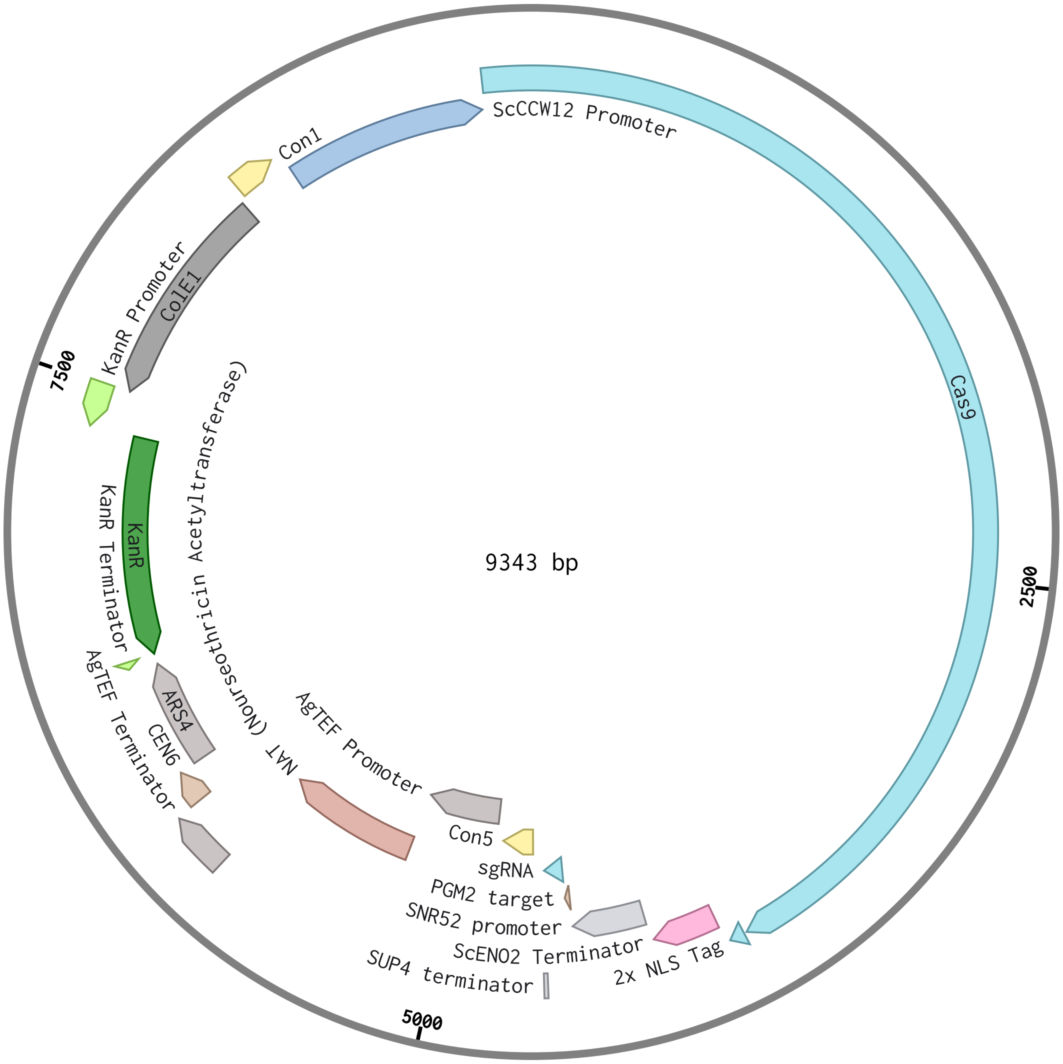

### ISA1045-pCCW12-Cas9-tSNR52-sgRNA-for-Site-1-NatR-CEN-KanR-ColE1-sequence.pdf

#### ISA1045 pCCW12, Cas9, tSNR52, sgRNA for Site 1...

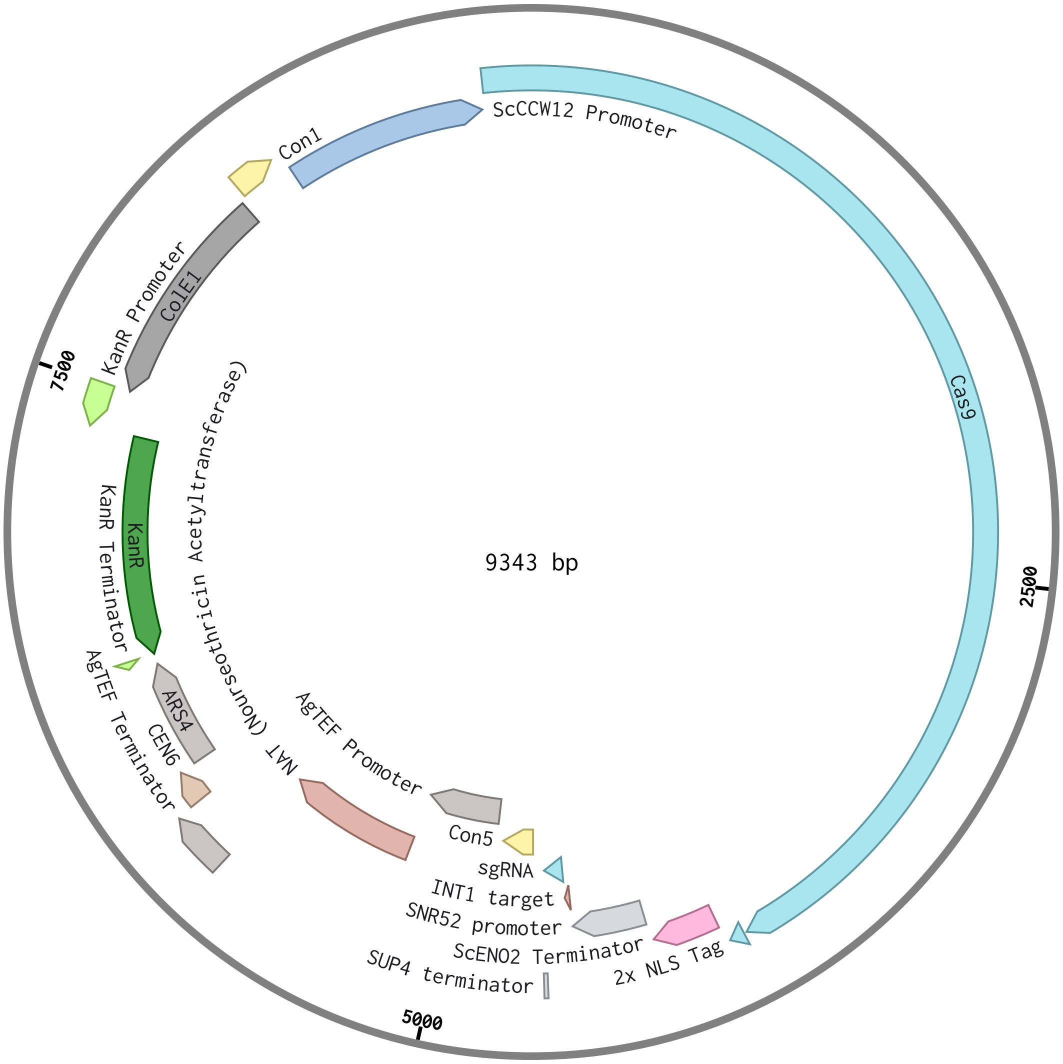

### ISA1130-pCCW12-Cas9-tSNR52-sgRNA-for-Site-5-NatR-CEN-KanR-ColE1-sequence.pdf

#### ISA1130 pCCW12, Cas9, tSNR52, sgRNA for Site 5,...

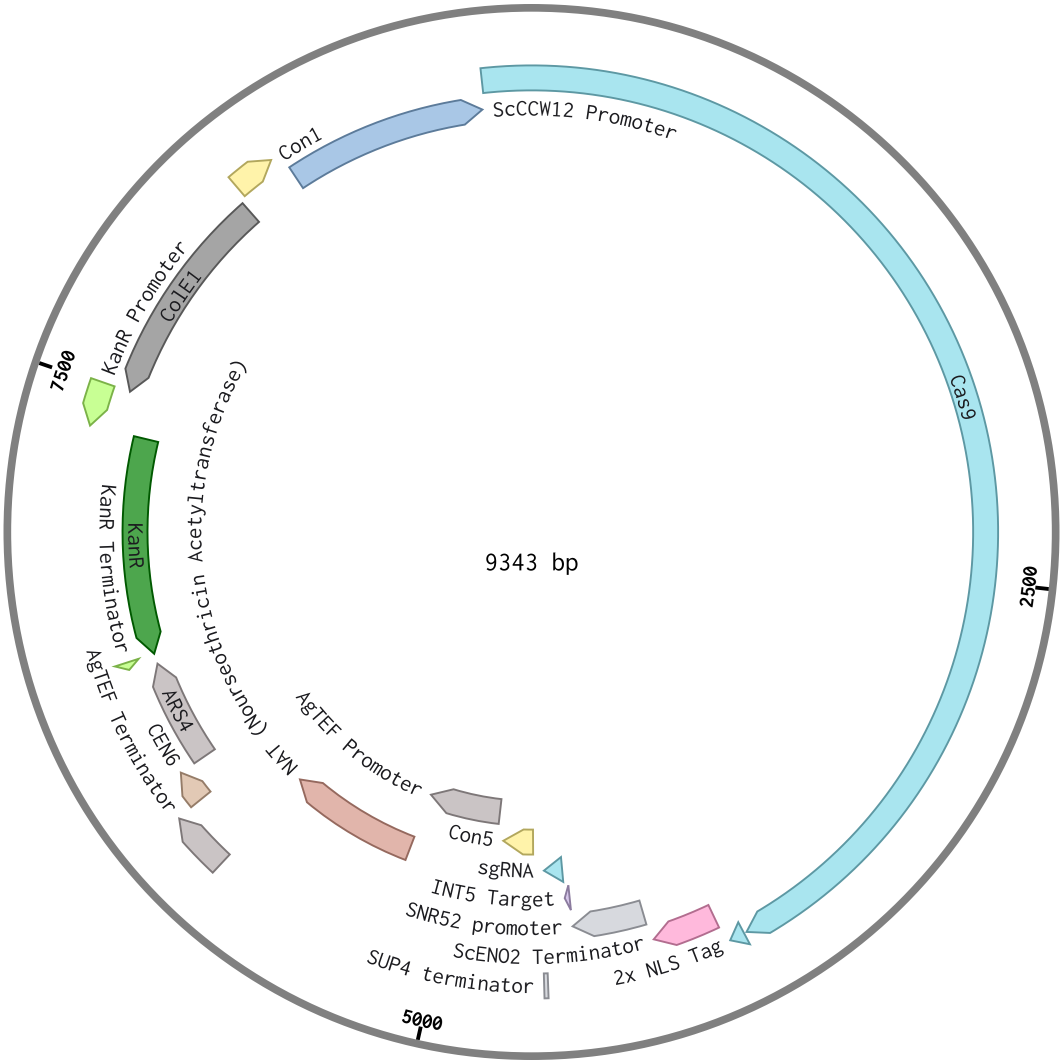

### ISA1132-pCCW12-mKate2-tENO2-URA-2micron-AmpR-ColE1-sequence.pdf

#### ISA1132 pCCW12, mKate2, tENO2, URA, 2micron, Am...

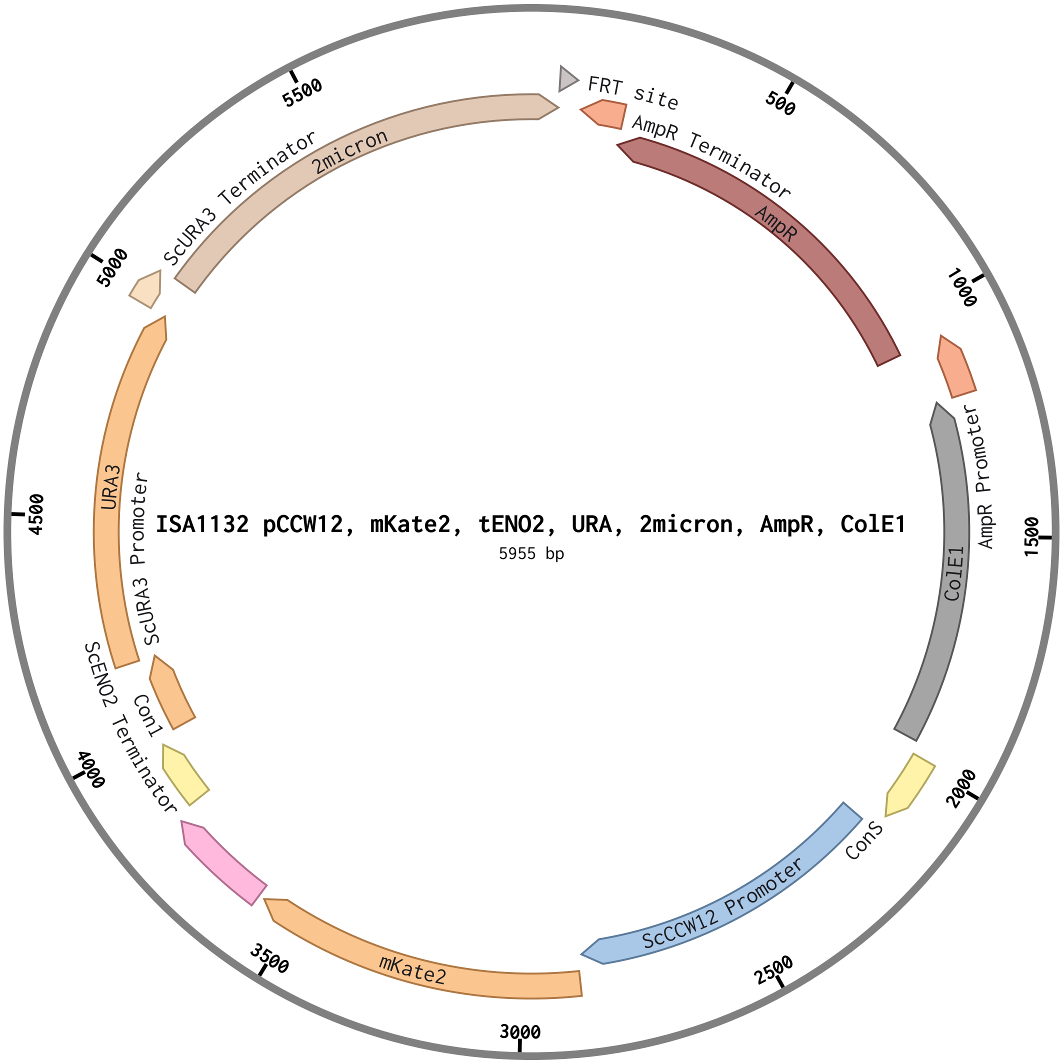

### ISA1149-pGAL-NanoLuc-tTDH1-URA-2micron-AmpR-ColE1-sequence.pdf

#### ISA1149, pGAL, NanoLuc, tTDH1, URA, 2micron, Am...

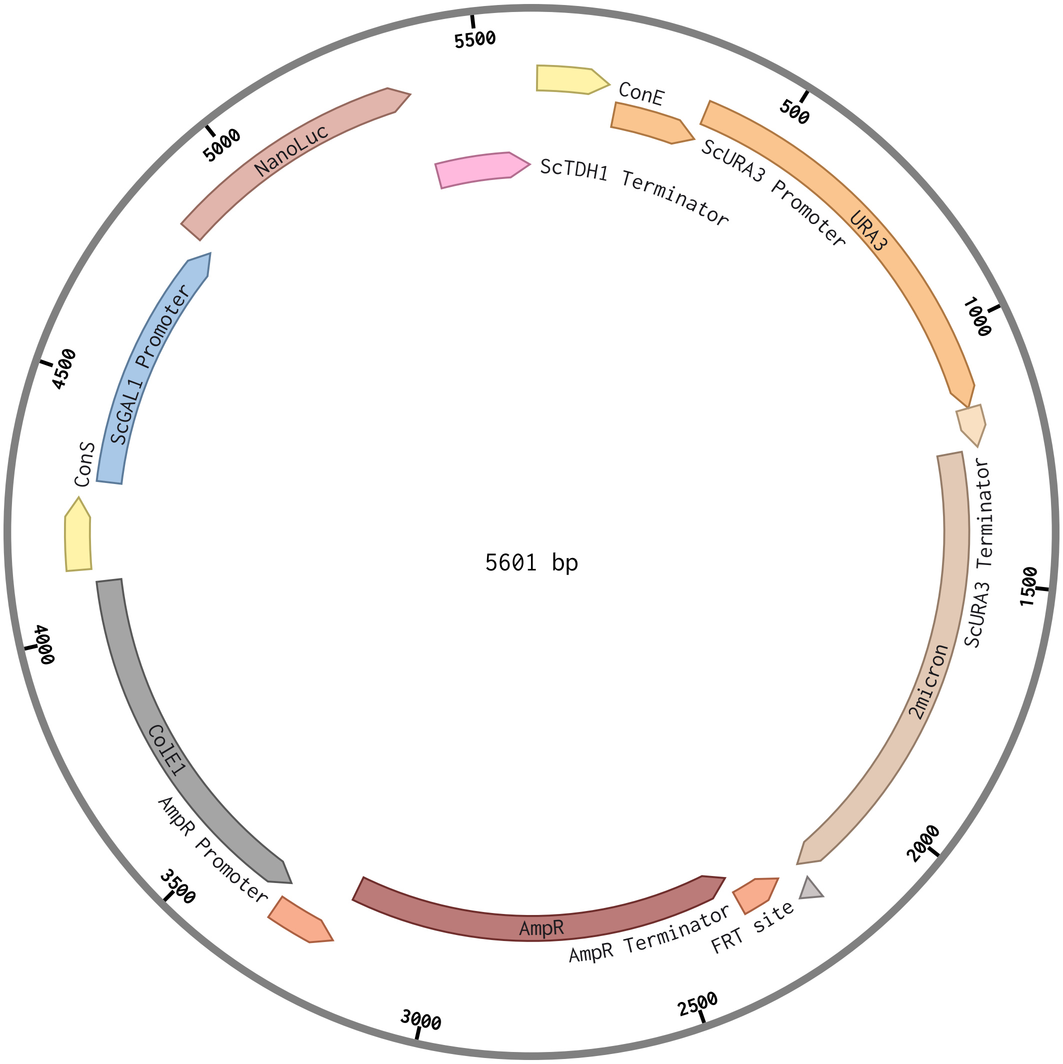

### ISA1150-pGAL-NanoLuc-tPGK1-INT-5-URA-AmpR-ColE1-sequence.pdf

#### ISA1150 pGAL, NanoLuc, tPGK1, INT 5, URA, AmpR,...

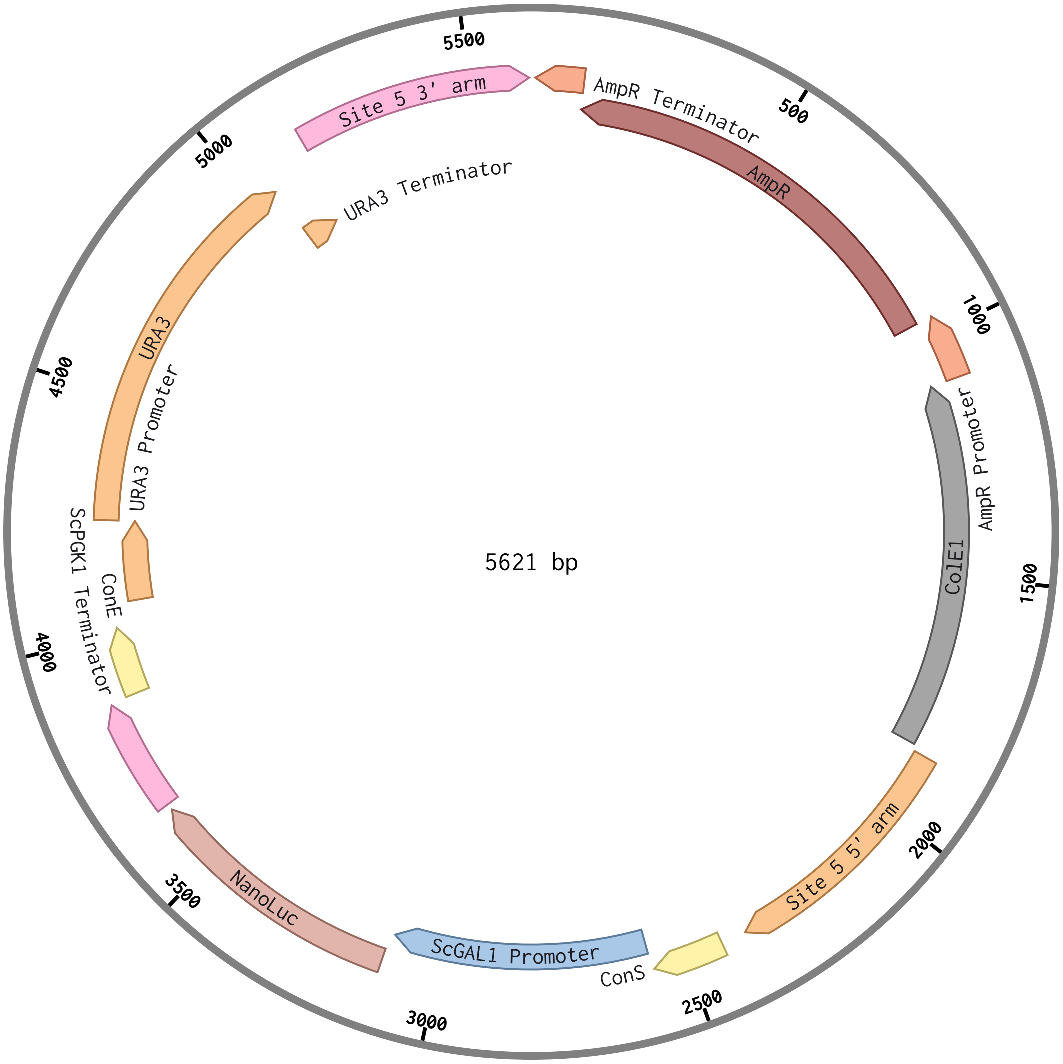

### pYD1-pGAL1-AGA2-TRP1-CEN-AmpR-ColE1-sequence.pdf

#### pYD1 pGAL1, AGA2, TRP1, CEN, AmpR-ColE1 (5562 bp)

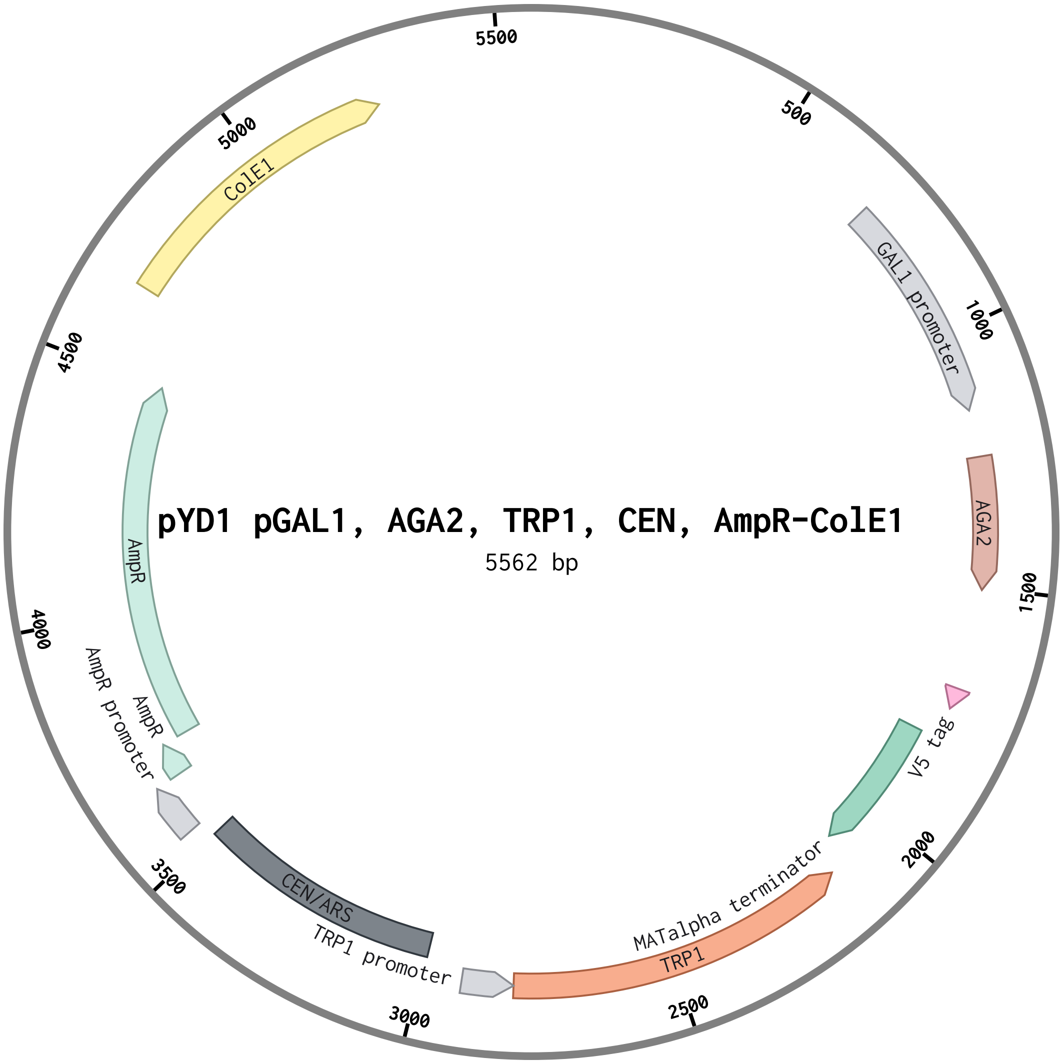
