## Supporting Information for "Programming Probiotics: Diet-responsive gene expression and colonization control in engineered *S. boulardii*"

**This PDF file includes:**

Figures S1 to S7
Tables S1 to S3
Legends for Datasets S1 to S3
SI References

**Other supporting materials for this manuscript include the following:**

Datasets S1 to S3

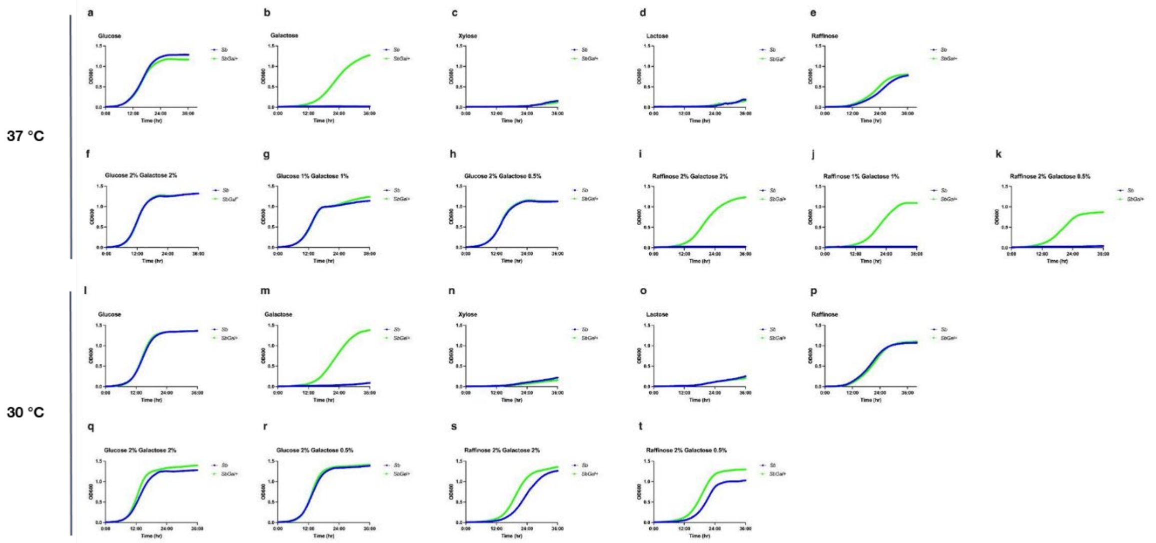

**Figure S1. Repaired galactose metabolism enables growth in inducing sugars at 37 °C and 30 °C.** (**a-k**) Growth of *Sb* and *SbGal*$^+$ in complete synthetic media (CSM) with various carbon sources at 37 °C. (**a**) *Sb* and *SbGal*$^+$ grow equivalently in glucose-containing media. (**b**) *SbGal*$^+$ exhibits a marked proliferation advantage in galactose-containing media. (**c-d**) Both strains show minimal growth on xylose or lactose, respectively. (**e**) *SbGal*$^+$ reveals a slightly enhanced ability to utilize raffinose at 37 °C. (**f-g**) Examination of growth in mixed glucose and galactose, underscoring the diauxic shift behavior of *SbGal*$^+$ when presented with limited glucose and abundant galactose. (**h**) Both strains grow equally well when glucose is in excess. (**i-k**) A stark contrast in growth is observed at 37°C with only raffinose and galactose available, highlighting the superior growth of *SbGal*$^+$. (**l-t**) Comparative growth analyses of *Sb* and *SbGal*$^+$ in complete synthetic media (CSM) with various carbon sources at 30°C. (**l-p**) Growth *Sb* and *SbGal*$^+$ in complete synthetic media (CSM) with various single carbon sources. (**q-t**) Examination of growth in mixed carbon sources at 30°C, again showing underscoring the diauxic shift behavior of *SbGal*$^+$ when presented with limited glucose or raffinose and abundant galactose. Three biological replicates were used for all growth curve measurements.

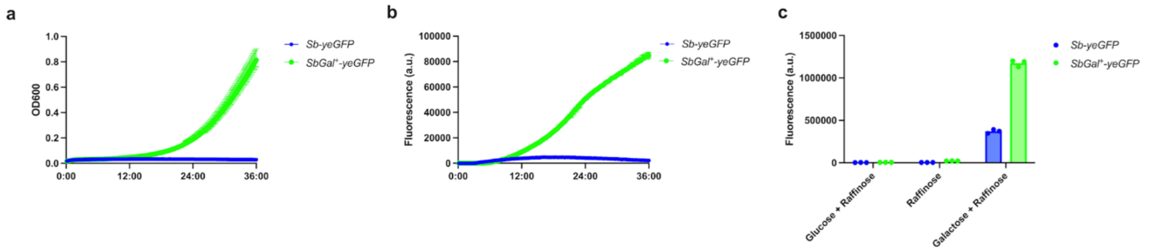

**Figure S2. Comparison of growth and expression of yeGFP in *Sb* and *SbGal⁺*.** (**a**) Growth of *Sb* and *SbGal*$^+$ containing *pGAL1-yeGFP* in complete synthetic media without uracil (CSM-U) with galactose. (**b**) Fluorescence levels over time of the strains in (**a**). (**c**) Per-cell fluorescence values (measured via flow cytometry) of both strains grown in glucose+raffinose, raffinose alone, or galactose+raffinose. Three biological replicates were used for all growth curve measurements.

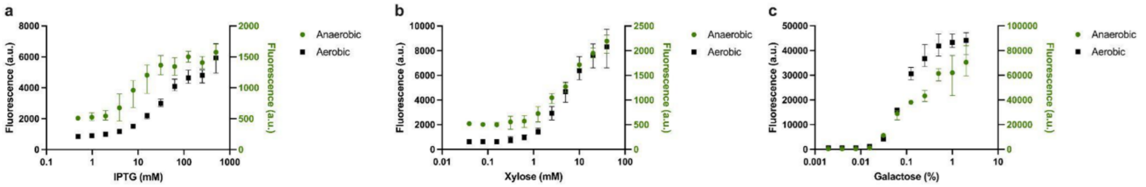

**Figure S3. Inducible systems enable tunable gene expression in aerobic and anaerobic environments.** Dose-response curves of yeast expressing CaFbfp under aerobic and anaerobic conditions using three inducible systems. Systems with the highest fold induction: *pLAC* (**a**), *pXYL* (**b**), and *pGAL1* (**c**) were cultured under aerobic and anaerobic conditions. (**a**) Strains expressing CaFbfp under the control of the *pLAC* promoter were cultured in synthetic media lacking uracil and histidine supplemented with glucose (2%) and 0.1-1000 mM IPTG. (**b**) Strains expressing CaFbfp under the control of the *pXYL* promoter were cultured in synthetic media lacking uracil and histidine supplemented with glucose (2%) and 0.01-100 mM xylose. (**c**) Strains expressing CaFbfp under the control of the *pGAL1* promoter were cultured in synthetic media lacking uracil supplemented with raffinose (2%) and 0.001-10% galactose. Three biological replicates were cultured for each condition and CaFbfp expression was measured for 24 hours.

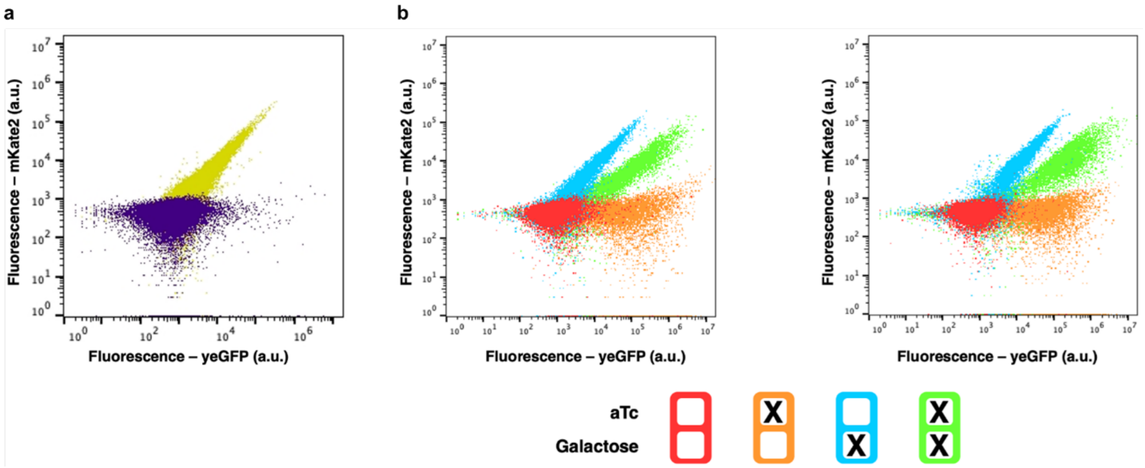

**Figure S4. Validation and replication of independent expression of dual fluorescent reporters in *S. boulardii.*** (**a**) Scatterplot of *Sb* harboring an empty vector(purple) or a constitutive mKate2 fluorescent reporter (**b**) Replicate scatter plots confirming the results shown in Figure 4.

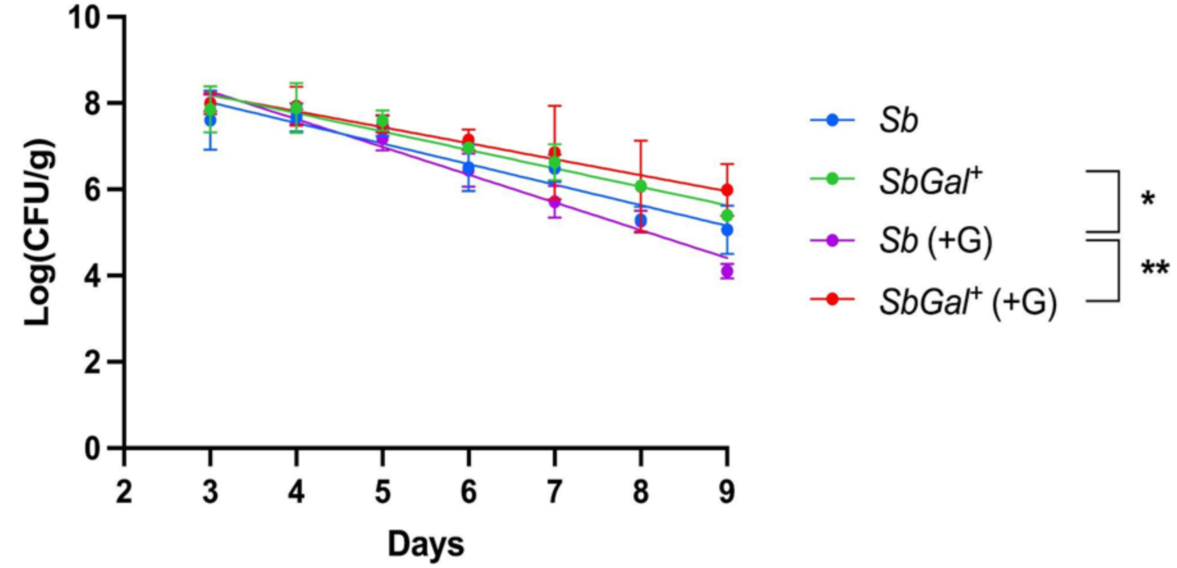

| | Equation | $R^2$ |
|---|---|---|
| *Sb* | Y = -0.4769*X + 9.448 | 0.8006 |
| *SbGal*$^+$ | Y = -0.4275*X + 9.479 | 0.8550 |
| *Sb* (+G) | Y = -0.6435*X + 10.20 | 0.9505 |
| *SbGal*$^+$ (+G) | Y = -0.3712*X + 9.297 | 0.6392 |

**Figure S5: Longitudinal colonization dynamics of *Sb* and *SbGal*$^+$ in antibiotic-treated mice.** This graph displays the average logarithmic counts of colony-forming units per gram (Log(CFU/g)) of fecal samples over time, depicting the colonization profiles of wild-type *Sb* and *SbGal*$^+$ in the presence and absence of galactose (G). Best fit lines to this logarithmically transformed data are provided below the plot. One-way ANOVA with Sidak's multiple comparisons test were conducted between the slopes of *Sb*, *SbGal*$^+$, Sb (+G) and *SbGal*$^+$ (+G) treatment groups. (ns P> 0.05, *P< 0.05, **P< 0.005).

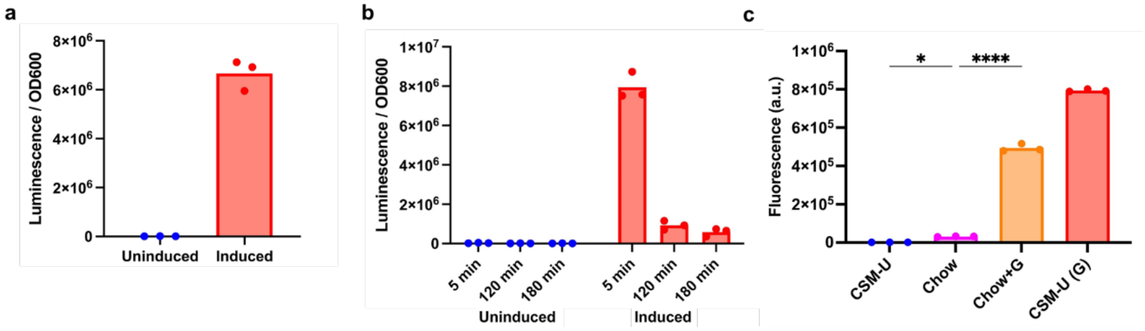

**Figure S6. Gene expression in yeast synthetic media and mouse chow diet.** (**a**) *Sb* expressing NanoLuciferase under the control of *pGAL1* in CSM-U with and without galactose. (**b**) Measurement of NanoLuciferase luminescence at 5, 120 and 180 minutes after Luciferase substrate addition. Each dot represents a biological replicate. (**c**) Fluorescence from *Sb* expressing yeGFP under the control of *pGAL1* in yeast complete synthetic media without uracil (CSM-U) and chow mice diet (Chow) with and without additional galactose. One-way ANOVA with Sidak's multiple comparisons test were conducted between CSM-U, Chow and Chow+G groups. (ns P> 0.05, *P< 0.05, **P< 0.005, ***P< 0.0005, ****P< 0.0001)

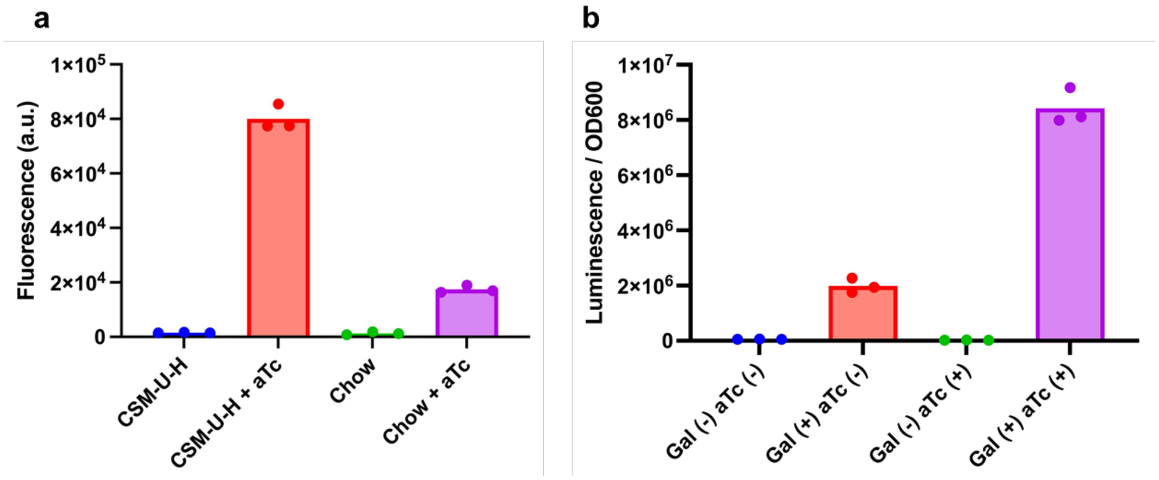

**Figure S7. *in vitro* characterization of genetic logic gates in *Sb*.** (**a**) *Sb* expressing yeGFP under the control of anhydrotetracycline inducible promoter (*pTET*) was tested in yeast complete synthetic media without uracil and histidine (CSM-U-H) and chow mice diet (Chow) with and without added aTc. (**b**) *Sb* expressing NanoLuciferase under the control of *pGAL-TET* in CSM-U-H in various combinations of inducers. Each dot represents a biological replicate.

**Table S1.** List of strains and plasmids

| Figure | Strain name | E.coli/S.b strain | Description | Selection Marker | Source |
|---|---|---|---|---|---|
| *Figure 2* | DJH077 | SbGal+ ΔURA ΔHIS | | | This study |
| | DD313 | Sb MYA 796 ΔURA ΔHIS | | | This study |
| | ISA061 | Sc BY4741 | | | |
| *Figure S2* | DJH098 | NEB 5 α | ConS, pGAL, yeGFP, tENO1, ConE, URA, 2μ, AmpR-ColE1 | AmpR | This study |
| | DJH107 | Sb MYA 796 ΔURA ΔHIS | ConS, pGAL, yeGFP, tENO1, ConE, URA, 2μ, AmpR-ColE1 | URA | This study |
| | DJH118 | SbGal+ ΔURA ΔHIS | ConS, pGAL, yeGFP, tENO1, ConE, URA, 2μ, AmpR-ColE1 | URA | This study |
| *Figure 3* | DJH088 | NEB 5 α | ConS, pXYL, yeGFP, tENO1, ConE, URA, 2μ, AmpR-ColE1 | AmpR | This study |
| | DJH089 | NEB 5 α | ConS, pLac, yeGFP, tENO1, ConE, URA, 2μ, AmpR-ColE1 | AmpR | This study |
| | DJH090 | NEB 5 α | ConS, pTet, yeGFP, tENO1, ConE, URA, 2μ, AmpR-ColE1 | AmpR | This study |
| | DJH091 | NEB 5 α | pXYL, CmR-ColE1 (holder) | CmR | This study, pXYL from [1] |
| | DJH092 | NEB 5 α | pLAC, CmR-ColE1 (holder) | CmR | This study, pLAC from [1] |
| | DJH093 | NEB 5 α | pTET, CmR-ColE1 (holder) | CmR | This study, pTET from [1] |
| | DJH094 | NEB Stable | ConS, pFBAI, xylR_NLS, tENO2, Con1, HIS3, CEN6/ARS4, AmpR-ColE1 | AmpR | This study |
| | DJH095 | NEB 5 α | ConS, pFBAI, lacI_NLS, tENO2, Con1, HIS3, CEN6/ARS4, AmpR-ColE1 | AmpR | This study |
| | DJH096 | NEB Stable | ConS, pFBAI, tetR_NLS, tENO2, Con1, HIS3, CEN6/ARS4, AmpR-ColE1 | AmpR | This study |
| | DJH097 | NEB 5 α | ConS, pCUP1, yeGFP, tENO1, ConE, URA, 2μ, AmpR-ColE1 | AmpR | This study |
| | DJH119 | SbGal+ ΔURA ΔHIS | ConS, pCUP1, yeGFP, tENO1, ConE, URA, 2μ, AmpR-ColE1 | URA | This study |
| | DJH120 | SbGal+ ΔURA ΔHIS | ConS, pXYL, yeGFP, tENO1, ConE, URA, 2μ, AmpR-ColE1; ConS, pFBAI, xylR_NLS, tENO2, | URA, HIS | This study |

| | | | | | |
|---|---|---|---|---|---|
| | | | Con1, HIS3, CEN6/ARS4, AmpR-ColE1 | | |
| | DJH121 | SbGal+ ΔURA ΔHIS | ConS, pLac, yeGFP, tENO1, ConE, URA, 2μ, AmpR-ColE1; ConS, pFBAI, lacI_NLS, tENO2, Con1, HIS3, CEN6/ARS4, AmpR-ColE1 | URA, HIS | This study |
| | DJH122 | SbGal+ ΔURA ΔHIS | ConS, pTet, yeGFP, tENO1, ConE, URA, 2μ, AmpR-ColE1; ConS, pFBAI, tetR_NLS, tENO2, Con1, HIS3, CEN6/ARS4 | URA, HIS | This study |
| *Figure S3* | DJH099 | NEB Stable | ConS, pGAL, CaFbFP, tENO1, ConE, URA, 2μ, AmpR-ColE1 | AmpR | This study |
| | DJH100 | NEB Stable | ConS, pCUP1, CaFbFP, tENO1, ConE, URA, 2μ, AmpR-ColE1 | AmpR | This study |
| | DJH101 | NEB Stable | ConS, pXYL, CaFbFP, tENO1, ConE, URA, 2μ, AmpR-ColE1 | AmpR | This study |
| | DJH102 | NEB Stable | ConS, pLAC CaFbFP, tENO1, ConE, URA, 2μ, AmpR-ColE1 | AmpR | This study |
| | DJH103 | NEB Stable | ConS, pTET, CaFbFP, tENO1, ConE, URA, 2μ, AmpR-ColE1 | AmpR | This study |
| | DJH123 | SbGal+ ΔURA ΔHIS | ConS, pGAL, CaFbFP, tENO1, ConE, URA, 2μ, AmpR-ColE1 | URA | This study |
| | DJH124 | SbGal+ ΔURA ΔHIS | ConS, pCUP1, CaFbFP, tENO1, ConE, URA, 2μ, AmpR-ColE1 | URA | This study |
| | DJH125 | SbGal+ ΔURA ΔHIS | ConS, pXYL, CaFbFP, tENO1, ConE, URA, 2μ, AmpR-ColE1; ConS, pFBAI, xylR_NLS, tENO2, Con1, HIS3, CEN6/ARS4, AmpR-ColE1 | URA, HIS | This study |
| | DJH126 | SbGal+ ΔURA ΔHIS | ConS, pLAC CaFbFP, tENO1, ConE, URA, 2μ, AmpR-ColE1; ConS, pFBAI, lacI_NLS, tENO2, Con1, HIS3, CEN6/ARS4, AmpR-ColE1 | URA, HIS | This study |
| | DJH127 | SbGal+ ΔURA ΔHIS | ConS, pTET, CaFbFP, tENO1, ConE, URA, 2μ, AmpR-ColE1; ConS, pFBAI, tetR_NLS, tENO2, Con1, HIS3, CEN6/ARS4, AmpR-ColE1 | URA, HIS | This study |
| *Figure 4* | DD576 | NEB 5 α | ConS, pGAL1, AGA1, tENO1, ConE, URA, Site 1 integration arms, AmpR-ColE1 | AmpR | This study |
| | DD579 | SbGal+ ΔURA ΔHIS ΔPrb1 ΔPep4 | ConS, pGAL1, AGA1, tENO1, ConE, URA (integrated Site 1) | | This study |
| | pYD1 | NEB 5 α | pGAL1, AGA2, TRP1, CEN, AmpR-ColE1 | AmpR, TRP | [2] |

| | | | | | |
|---|---|---|---|---|---|
| | DD580 | NEB 5 α | pGAL1, AGA2, tMATa, HIS3, CEN6/ARS4, AmpR-ColE1 | AmpR | This study |
| | DD608 | NEB 5 α | ConS, pGAL1, AGA2, SA1, tMATa, Con1, HIS3, CEN6/ARS4, AmpR-ColE1 | AmpR | This study |
| | DD613 | SbGal+ ΔURA ΔHIS ΔPrb1 ΔPep4 | ConS, pGAL1, AGA1, tENO1, ConE, URA (integrated site 1); pGAL1, AGA2, SA1, tMATa, Con1, HIS3, CEN6/ARS4, AmpR-ColE1 | HIS | This study |
| | ISA086 | NEB 10 β | ConS, GFP 234 Dropout, ConE, URA, Site 1 integration arms, AmpR-ColE1 | AmpR | This study |
| | ISA186 | NEB Top10 | ConS, pAgTef, NatR, tAgTef, Site 1 integration arms, NatR, AmpR-ColE1 | AmpR | This study |
| *Figure 5* | DD592 | NEB 5 α | ConS, pGAL1, mKate2, tENO1, ConE, tENO1, yeGFP, pTET, Con1, URA3, 2μ, AmpR-ColE1 | AmpR | This study |
| | DD598 | SbGal+ ΔURA ΔHIS | ConS, pGAL1, mKate2, tENO1, ConE, tENO1, yeGFP, pTET, Con1, URA3, 2μ, AmpR-ColE1; ConS, pFBAI, tetR_NLS, tENO2, Con1, HIS3, CEN6/ARS4, AmpR-ColE1 | URA, HIS | This study |
| *Figure S4* | ISA1132 | NEB 5 α | ConS, pCCW12, mKate2, tENO2, Con1, URA3, 2μ, AmpR-ColE1 | AmpR | This study |
| | ISA1135 | Sb MYA 7976 ΔURA | ConS, pCCW12, mKate2, tENO2, Con1, URA3, 2μ, AmpR-ColE1 | URA | This study |
| | ISA1155 | SbGal+ ΔURA ΔHIS | ConS, pCCW12, mKate2, tENO2, Con1, URA3, 2μ, AmpR-ColE1 | URA | This study |
| *Figure 6* | ISA1151 | Sb MYA 796 ΔURA ΔHIS | NatR (integrated Site 1) | NatR | This study |
| | DJH154 | SbGal+ ΔURA ΔHIS | NatR (integrated Site 1) | NatR | This study |
| *Figure S5/Figure 7* | ISA1149 | NEB 5 α | ConS, pGAL1, NanoLuc, tTDH1, ConE, URA3, 2μ, AmpR-ColE1 | AmpR | This study |
| | ISA1150 | NEB 5 α | ConS, pGAL1, NanoLuc, tPGK1, ConE, URA3, Site 5 integration arms, AmpR-ColE1 | AmpR | This study |
| | ISA1154 | SbGal+ ΔURA ΔHIS | ConS, pGAL1, NanoLuc, tPGK1, ConE, URA3 (integrated Site 5) | | This study |
| *Figure 8* | DD593 | NEB 5 α | ConS, pGalTet, NanoLuc, tSSA1, Con1, URA3, 2μ, AmpR-ColE1 | AmpR | This study |
| | DD599 | SbGal+ ΔURA ΔHIS | NatR (integrated Site 1); ConS, pGalTet, NanoLuc, tSSA1, Con1, URA3, 2μ, AmpR-ColE1; ConS, pFBAI, tetR_NLS, tENO2, Con1, HIS3, CEN6/ARS4, AmpR-ColE1 | URA, HIS, NatR | This study |

| | | | | | |
|---|---|---|---|---|---|
| *Figure S6* | DD594 | NEB 5 α | ConS, pGalTet, yeGFP, tENO1, ConE, URA3, 2u, AmpR-ColE1 | AmpR | This study |
| | DD601 | SbGal+ ΔURA ΔHIS | NatR (Site 1); ConS, pGalTet, yeGFP, tENO1, ConE, URA3, 2µ, AmpR-ColE1; ConS, pFBAI, tetR_NLS, tENO2, Con1, HIS3, CEN6/ARS4, AmpR-ColE1 | URA, HIS, NatR | This study |
| *Yeast Integration* | ISA1041 | NEB 5 α | pCCW12, Cas9, tENO2, pSNR52, sgRNA for PGM2, tSUP4, Con5, pAgTef, NatR, tAgTef, CEN6/ARS4, KanR, ColE1a | KanR | This study |
| | ISA1045 | NEB 5 α | pCCW12, Cas9, tENO2, pSNR52, sgRNA for Site 1, tSUP4, Con5, pAgTef, NatR, tAgTef, CEN6/ARS4, KanR, ColE1a | KanR | This study |
| | ISA1130 | NEB 5 α | pCCW12, Cas9, tENO2, pSNR52, sgRNA for Site 5, tSUP4, Con5, pAgTef, NatR, tAgTef, CEN6/ARS4, KanR, ColE1a | KanR | This study |

**Table S2.** List of gBlocks

| gBlock name | Description |
|---|---|
| gb_pFBAI | pFBAI |
| gb_tetR_NLS | tetR_NLS |
| gb_lacI_NLS | lacI_NLS |
| gb_xylR_NLS | xylR_NLS |
| gb_CaFbFP | CaFbFP |
| gb_mKate2 | mKate2 |
| gb_NanoLuc | NanoLuc |
| gb_pGalTet | pGalTet |

**Table S3**. List of primers

| **Primer Sequence** | **Description** |
|---|---|
| GTATCAATTCGCATTCTTATATTTAATACATA | Check integration, site 1 |
| CTGAGAAACTGGTGAATAATTCGATAA | Check integration, site 1 |
| GATCTAAAGTAATCGTCGAGAC | Check integration, site 5 |
| GTTGAAGGGGTTTCTTAAGG | Check integration, site 5 |
| CCCAGTTGTTTGTAGCTGGTT | Amplify linear donor DNA, site 1 |
| TTGTTGGCATTCCATTGTTG | Amplify linear donor DNA, site 1 |
| GGTCGTTTTTGTGCAGCATATTG | Amplify linear donor DNA, site 5 |
| GAGCTTACTCTATATATTCATTCTTGTCAGTTTC | Amplify linear donor DNA, site 5 |
| GTGGTGAAGAATCGTTTGGT | Amplify *PGM2* fragment from S. cerevisiae |
| GACCTTAAGAGACTCATCGG | Amplify *PGM2* fragment from S. cerevisiae |

**Dataset S1 (separate file).** Plasmid maps for all plasmids generated in this study in PDF format. The plasmid names correspond to the names listed in Table S1.

**Dataset S2 (separate file).** Maps for all gene fragments generated in this study in PDF format.

**Dataset S3 (separate file).** Genbank files for all plasmids and gene fragments generated in this study in genbank format.
